# Evolution of the non-visual and visual opsin gene repertoire in ray-finned fishes

**DOI:** 10.1101/2024.10.03.616438

**Authors:** Maxime Policarpo, Lily G. Fogg, Fabio Cortesi, Walter Salzburger

## Abstract

Photoreception – the detection of light for image formation (vision) as well as for non-image-forming purposes (circadian regulation, DNA repair) – is critical to the survival of most animals. In vertebrates, photoreception is mediated by opsin proteins, which are classified, according to their function, into visual and non-visual opsins. Here, we provide the most comprehensive study to date on the evolution of the opsin gene family in the largest class of vertebrate, fishes, with a particular focus on the understudied non-visual opsins. Based on an in-depth analysis of 535 high-quality genomes, we document great variation in gene numbers in the different opsin gene subfamilies across ray-finned fishes and show that visual opsins are more prone to duplications and losses than non-visual opsins. We provide compelling evidence that visual and non-visual opsins co-evolve in ray-finned fishes, both in terms of copy numbers and selective pressures acting on their coding sequences, probably in response to the different photic environments they inhabit. Species that live in dim light or all in the dark (such as in caves or the deep sea) convergently reduced their visual and non-visual opsin gene repertoires, while polar species feature accelerated evolution in both. Fishes that rely on electroreception show a reduction in the number of visual and non-visual opsin genes and accelerated evolution of the remaining genes. We further found that genes of the phototransduction cascade co-evolve with opsins. Finally, that non-visual opsins are mainly expressed in testes and ovaries (next to the eyes) supports a function in gamete biology.

## Introduction

The sensory systems of animals play fundamental roles in vital tasks such as orienting themselves in space and time, tracking down food, avoiding predators, finding shelter, and selecting mating partners (1–4). Animal sensory systems, and the genes underlying them, are thus under strong Darwinian selection (2, 5, 6). One essential sensory system, especially in vertebrates, is photoreception, *i*.*e*., the detection of photons for image-forming (visual) and non-image-forming (non-visual) purposes (7). Among the core elements of photoreception are opsin genes, which encode for G protein-coupled receptors (8) and are classified, according to whether or not they are involved in image formation, into visual and non-visual opsins (8).

The function and evolution of visual opsin genes have been well characterized in many vertebrate clades (4, 9–11). Most vertebrates possess two types of visual opsins, rhodopsin(s) (*rh1*) that mediate scotopic (dim-light) vision and cone opsins that are responsible for photopic (bright-light) and color vision (12). The vertebrate ancestor most probably possessed a single *rh1* gene and members of four cone opsin gene subfamilies, each permitting the detection of a specific waveband of the visible light spectrum: the short wavelength sensitive opsins *sws1* and *sws2*, the mid-wavelength sensitive *rh2*, and the long wavelength sensitive *lws* (12). Vision is initiated by a photon-induced conformational change of a visual opsin protein covalently bound to a vitamin-A derived chromophore, forming the photopigment. This activates transducins, which in turn activate phosphodiesterases that degrade cGMP, ultimately leading to the closure of cGMP-gated ion channels and a hyperpolarization of the photoreceptor cell (13). Other genes required for vision include regulators of this cascade (such as arrestins and guanylyl cyclases), GC-activating proteins, and retinoid isomerohydrolases (13, 14), but also crystallins, which are necessary for a proper transparency and refractive index of the lens (15, 16).

Much less is known about the evolution of non-visual opsin genes and the associated phototransduction cascade in vertebrates. This is surprising, given that non-image forming photoreception initiates several critical biological processes, including circadian clock photoentrainment (8), DNA repair activation (9, 10), neuronal development and activity (11), and seasonality, *e*.*g*., in reproduction (12). It is most likely because of these diverse functions that the non-visual opsin gene repertoire is much larger than that of visual opsin genes (7, 17), with more than twenty subfamilies of non-visual opsins known to date (17, 18), including *pinopsin* (19), melanopsins (*opn4*) (20), teleost multiple tissue opsins (*tmt*) (21), retinal G protein receptors (*rgr*) (22), and neuropsin (*opn5*) (23).

The most species-rich class of vertebrates, ray-finned fishes (actinopterygians), are particularly interesting in the context of opsin evolution, as members of this group experience a diverse range of photic environments (4), depending on water depth and turbidity, latitude, and diel activity patterns, among others (24–32). Accordingly, actinopterygians feature substantial interspecific variation in the extent and composition of their visual opsin gene repertoire (33, 34). This is partially due to differences in gene retention after the teleost-specific whole genome duplication (18, 35) and recent genome duplications in certain groups (36). On the other hand, the dynamics of non-visual opsin evolution in fishes remains elusive.

Here, we provide the most comprehensive examination to date of opsin gene evolution in ray-finned fishes, focusing on the understudied non-visual opsin genes. By analyzing 535 high-quality fish genomes, we characterize their opsin gene repertoire and show that visual and non-visual opsins co-evolve in this group, driven mainly by the prevailing light-environment. The evolution of other genes involved in photoreception and in circadian clock regulation is tightly linked to the opsin gene repertoire. We also analyzed gene expression patterns in 75 species, and found that, next to the eye, non-visual opsins are highly expressed in gonadal tissues, suggesting that – just like in jellyfishes, mice and humans (37–39) – they also function in gamete release, maturation and guidance in fish.

## Results

### Characterization of the opsin gene repertoire in ray-finned fishes

We first characterized the entire opsin gene repertoire – comprising the five subfamilies of visual opsins and 22 subfamilies of non-visual opsins – in 535 high-quality genome assemblies representing the taxonomic and ecological diversity of ray-finned fishes (Fig. 1, Fig. S1-S35). Applying a standardized gene mining procedure, we detected a mean number of 25.1 non-visual opsin genes per actinopterygian genome. Tetraploid fishes featured about twice as many non-visual opsins as diploid species (mean of 47 and 24 genes, respectively, Fig. S36), with up to 60 genes found in the goldfish (*Carassius auratus*). Hence, they represent the vertebrates with the by far most extensive repertoires of non-visual opsin genes known to date (18). Among diploid species, the migratory hilsa herring (*Tenualosa ilisha*) had the highest number of non-visual opsins (38 genes) (Fig. S36, Dataset S1). The smallest non-visual opsin gene repertoires were found in the abyssal Mariana snailfish (*Pseudoliparis swirei*) with seven genes, followed by two mormyrids (*Mormyrops zanclirostris* and *Stomatorhinus walker*) with eight genes each (Fig. S36, Dataset S1).

**Figure 1.**
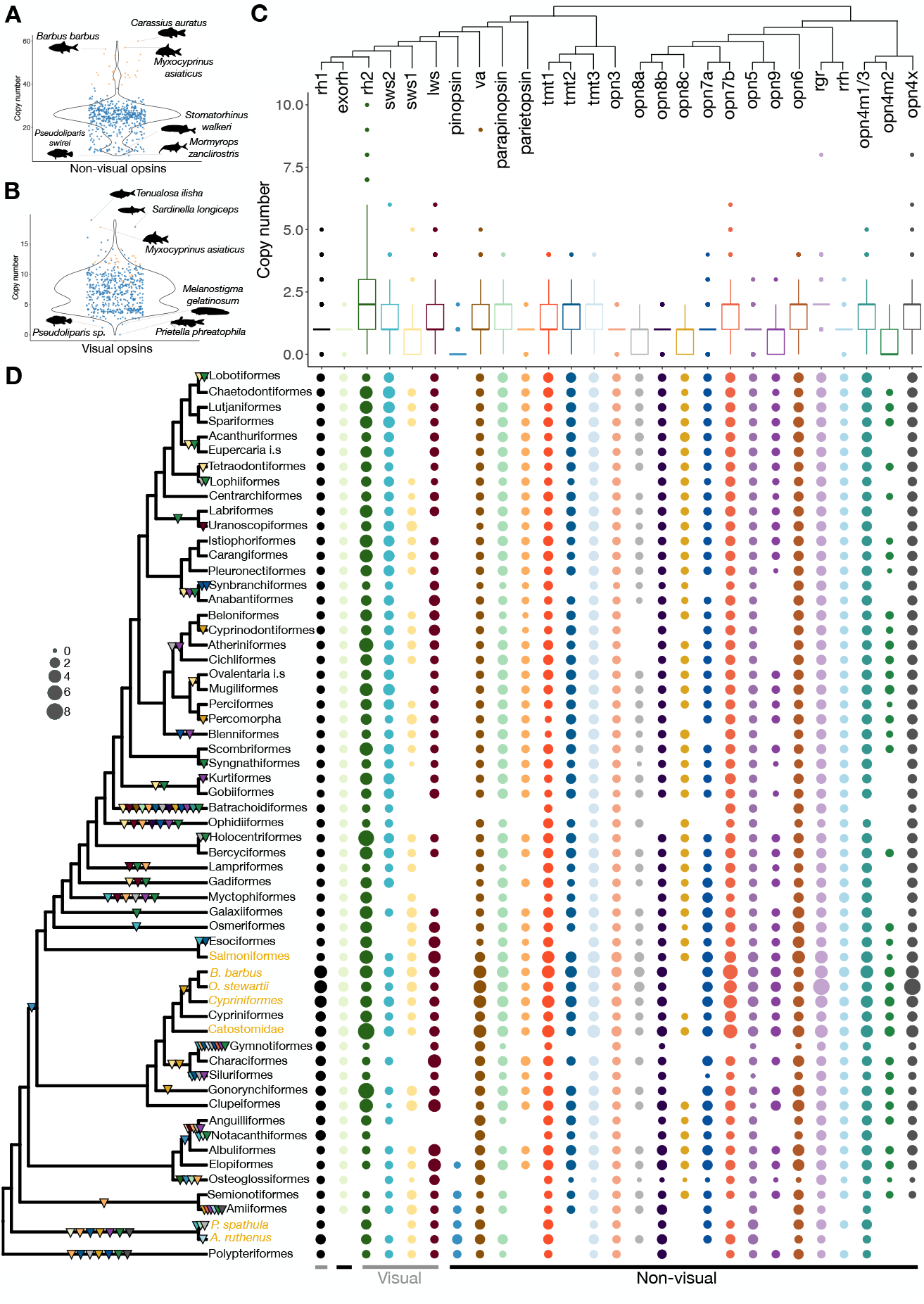
Evolution of the opsin gene repertoire in ray-finned fishes. Number of non-visual opsin genes (A) and visual opsin genes (B) per species represented as violin plots. Blue dots indicate diploid species, orange dots correspond to species with additional whole genome duplications. The names of the three species with the highest and the lowest gene numbers are shown. (C) Gene-tree of opsin gene subfamilies, with the number of complete genes per species and subfamily represented as boxplots. (D) Phylogeny of ray-finned fishes inferred using ASTRAL and dated using the least square dating method. Order-or species-specific whole-genome duplications are indicated with orange tip labels. Five branches correspond to Cypriniformes: one black branch for diploid species, one orange branch for tetraploid species of the genus Carassius, Cyprinus and Sinocyclocheilus as well as three orange branches for Catostomidae, O. stewartia and B. barbus that underwent linages-specific WGD. Triangles represent the complete loss of one opsin gene subfamily and are colored accordingly. Circles on the right of the phylogeny represent the presence of a particular opsin gene subfamily in the given lineage, and the size of the circle is scaled to the mean number of complete genes per species in this lineage. The number of opsins per species and the mean number of opsins per order are reported in Supplementary Data 1. Details on the opsin gene families lost at a particular branch can be retrieved from Fig. S35.

In agreement with previous results (40), we found a mean number of 7.3 visual opsin genes per species. The largest visual opsin gene repertoire in our set of species was found in *T. ilisha* (19 genes), followed by the Indian oil sardine (*Sardinella longiceps*; 18 genes) (Fig. S33, Dataset S1), since the teleost species with the greatest number of visual opsins observed to date (*Diretmus argenteus*) (40) was not included in here due to poor genome assembly quality. The species with the fewest visual opsins were the Mexican blind catfish (*Prietella phreatophila*), which lives in constant darkness and entirely lost its visual opsins, two abyssal snailfish species (*Pseudoliparis sp. Yap Trench*: 1 gene; *P. swirei*: 2 genes), and two deep-sea eelpouts (*Melanostigma gelatinosum* and *Ophthalmolycus amberensis*) with two genes each.

At the order level, we found that tarpons (Elopiformes) and herrings and anchovies (Clupeiformes), many of which are migratory and experience diverse photic conditions over their lifespans (41), had the largest non-visual opsin repertoires (mean = 34 and 33.5, respectively). Clupeiformes also had the largest visual opsin repertoire (mean = 12.8) (Fig. 1D, Fig. S33, Dataset S1). On the other hand, knifefishes (Gymnotiformes), which rely on electric communication, are primarily nocturnal, and live in turbid waters (42), had the lowest number of non-visual (mean = 9.8) and visual opsin genes (mean = 3.3), lacking entirely representatives of 9 opsin subfamilies (Fig. 1D, Fig. S35). We also detected a complete loss of 13 opsin subfamilies in Batrachoidiformes and seven in Ophidiiformes (Fig. 1D, Fig. S35). However, these were only represented by a single species each in our data set.

The number of pseudogenes also varied substantially between species and correlated negatively with the number of functional genes (Fig. S37). Species that experienced recent whole-genome duplications had more pseudogenes than diploid ones (Fig. S36), likely due to the non-functionalization of redundant ohnologs (43, 44). The highest number of opsin pseudogenes was detected in two tetraploid and blind cave cyprinids (*Sinocyclocheilus anophthalmus* and *S. anshuiensis* with 29 and 23 pseudogenes, respectively). Many opsin pseudogenes were also found in two diploid deep-sea species (*P. sp. Yap Trench*: 25; *Dissostichus mawsoni*: 21) (Fig. S36).

So far, it has been assumed that *pinopsin* was lost before the most recent common ancestor (MRCA) of teleosts (19, 45). Our finding of a complete *pinopsin* under purifying selection in the elopiform *Megalops cyprinoides* (Fig. S38) refutes this hypothesis. Instead, it supports a more complex evolutionary history of *pinopsin* with at least three independent losses (Fig. 1D, Fig. S35).

Overall, our results show that the non-visual and visual opsin gene repertoires of ray-finned fishes are shaped by recurrent gene duplications and losses and are much more diverse than previously assumed (4, 18).

### Dynamics of opsin gene evolution in actinopterygians

Next, to reconstruct the evolutionary history of each opsin gene subfamily in ray-finned fishes, we used a gene tree-species tree reconciliation method (46). The most dynamic subfamilies among non-visual opsins were *va* (birth rate = 0.0018 and death rate = 0.0064 per gene per million years), *opn4m1/3* (0.0017/0.008) and *opn3* (0.0017/0.006). In contrast, the least dynamic subfamilies were *pinopsin* (0.0004/0.002), *opn8c* (0.001/0.005) and *opn8a* (0.001/0.006) (Fig. 2A). The evolutionary rates of the visual opsins were significantly higher than that of non-visual opsins (Wilcoxon test p-values = 1.7e-4 and 5e-5 for birth and death rates respectively, Fig. S39), with *rh2* having the highest rates (0.0031/0.01), followed by *lws* (0.003/0.0095) and *sws1* (0.0023/0.008) (Fig. 2A). This suggests stronger selective pressures to modulate the gene repertoire involved in image-forming compared to non-image-forming photoreception.

**Figure 2.**
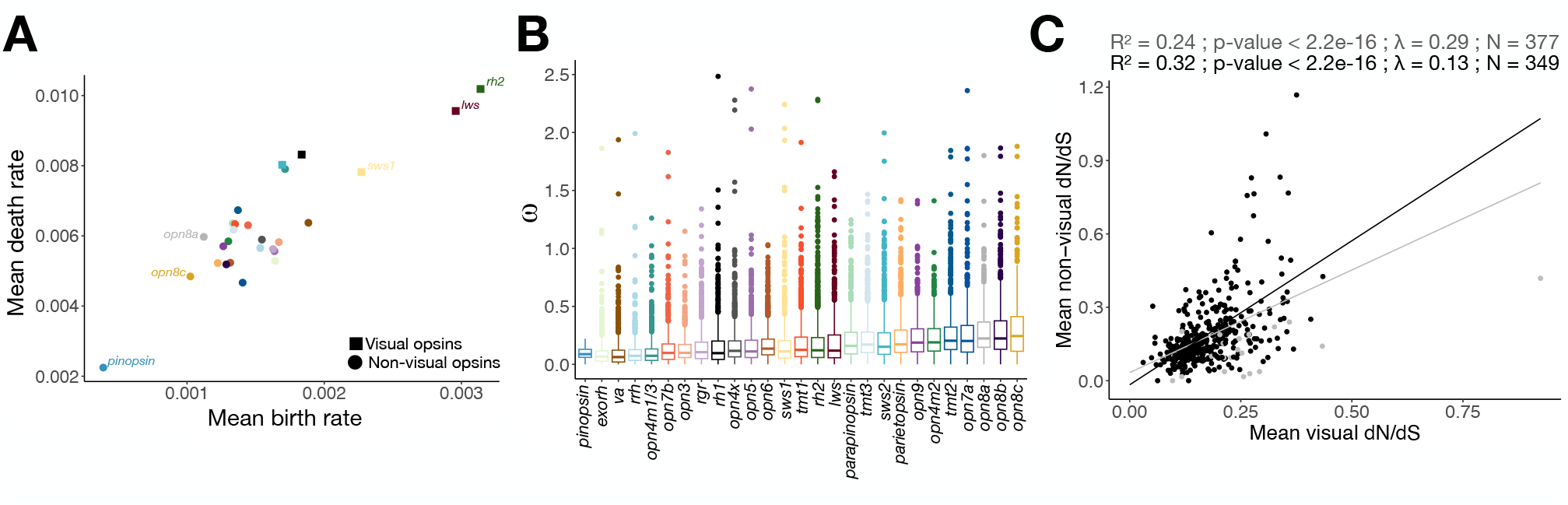
Evolutionary dynamics of opsin genes in ray-finned fishes. (A) Mean birth- and mean death-rate per opsin gene subfamily inferred from gene tree - species tree reconciliations. The name of the three subfamilies with the highest and lowest birth rates are indicated. (B) Distribution of dN/dS (ω) per opsin gene subfamily. (C) Correlation between the mean ω of non-visual opsins and the mean ω of visual opsins. pGLS R^2^, p-value and λ and regression lines considering all ray-finned fishes (gray), or only diploid teleosts (black) are shown. Only species with dN/dS computed for at least three non-visual and three visual opsins were retained (Material and method, Supplementary data 1). N reports the number of genomes used.

We then computed dN/dS (ω) for every opsin gene in each subfamily and found significant differences between subfamilies (Fig. 2B, Fig. S40). The lowest mean ω was found for *pinopsin, exorh* and *va* (0.09, 0.1, and 0.11, respectively), and the highest mean ω was detected for *opn8a, opn8b* and *opn8c* (0.297, 0.303, and 0.33, respectively) (Fig. 2D). No differences were found in the distribution of ω between non-visual and visual opsins (Fig. S41). We found no evidence of an association between birth and death rates and differences in the mean or median ω across opsin subfamilies (Fig. S42). However, we observed that episodes of positive selection typically follow opsin gene duplications, as branches resulting from duplication events had significantly higher ω-values than branches resulting from speciation events (except for *pinopsin* and *opn9*; Fig. S43). This is a strong indication of functional diversification in the aftermath of duplications in opsin genes, for example, by changing spectral sensitivity or thermal stability (24, 34).

Most species had a mean ω-value for their opsin genes of around 0.2 (Fig. S44). Again, tetraploid species had higher ω-values than diploids, likely because many of their opsins either evolve neutrally (43) (pseudogenes with no apparent loss-of-function mutations yet) or are subject to neo-or sub-functionalization (43). The highest mean ω (0.8) was observed in the tetraploid and eyeless cave cyprinid *S. anophthalmus* (Fig. S44). Accelerated opsin sequence evolution was also detected in the weakly electric fish *Mormyrus iriodes* (mean ω = 0.78) and in two deep-sea fishes, the scaly rockcod (*Trematomus loennbergii*; mean ω = 0.56) and the eelpout (*O. amberensis*; mean ω = 0.57) (Fig. S44). Importantly, we observed a positive correlation between the mean ω of non-visual opsins and the mean ω of visual opsins (pGLS: R^2^ = 0.24; p-value < 2.2e-16, Fig. 2C), indicating that the coding sequences of non-visual and visual opsins evolve concertedly and possibly under similar selective constraints. This also implies that visual opsin sequence evolution is predominantly determined by the overall photic environment (light-sensing) rather than more specific functions, such as the detection of the color of mates or prey (color vision), which is consistent with the idea that the latter is more likely regulated through spectral tuning, *e*.*g*., via differential gene expression (47–49).

Our analyses also revealed a positive correlation between the total number of non-visual and visual opsin genes in actinopterygian genomes (pGLS: R^2^ = 0.23; p-value < 2.2e-16), as well as between the number of genes in many of the opsin gene subfamilies (Fig. S45), irrespective of whether all species or only diploid species were considered (Fig. S45). We thus present conclusive evidence that non-visual opsins and visual opsins co-evolve at the macro-evolutionary scale of ray-finned fishes with respect to both sequence evolution (see above) and copy-number.

### Co-evolution of opsins and other genes

To test for co-evolutionary signatures between opsins and other genes, we took advantage of 155 annotated fish genomes, and inspected if the opsin gene numbers and mean ω-values correlated with those of other genes arranged in orthogroups (OGs) (Fig. 3, Fig. S46). We found that the number of non-visual opsins was positively correlated with gene numbers in 48 OGs, and negatively with 115 OGs (Fig. 3A). Among the positively correlated OGs were much more light-perception and circadian clock OGs than expected at random (Fig. S47), including phosphodiesterases (*pde6g*), guanylate cyclases (*gcap7*), two γ-crystallin gene families (one of which had the highest R^2^ value of all OGs), and the cryptochrome circadian regulators (*cry*) involved in the circadian clock feedback loop (50). Functional enrichment analysis of these 48 OGs also revealed an over-representation of genes in three reactome (51) pathways: neuronal system, transmission across chemical synapses, and postsynaptic signal transmission (Table S1). 71 OGs were under stronger purifying selection in species with a larger non-visual opsin gene repertoire (Fig. 3B). Among these were three genes known to be involved in vertebrate eye development: *bicdl1, fzd5*, and *rp2* (52–54). These 71 OGs were enriched for three reactome pathways (transport and synthesis of PAPS, Ca^2+^ pathway, and Beta-catenin independent WNT signaling) and one KEGG pathway (starch and sucrose metabolism) (Table S1).

**Figure 3.**
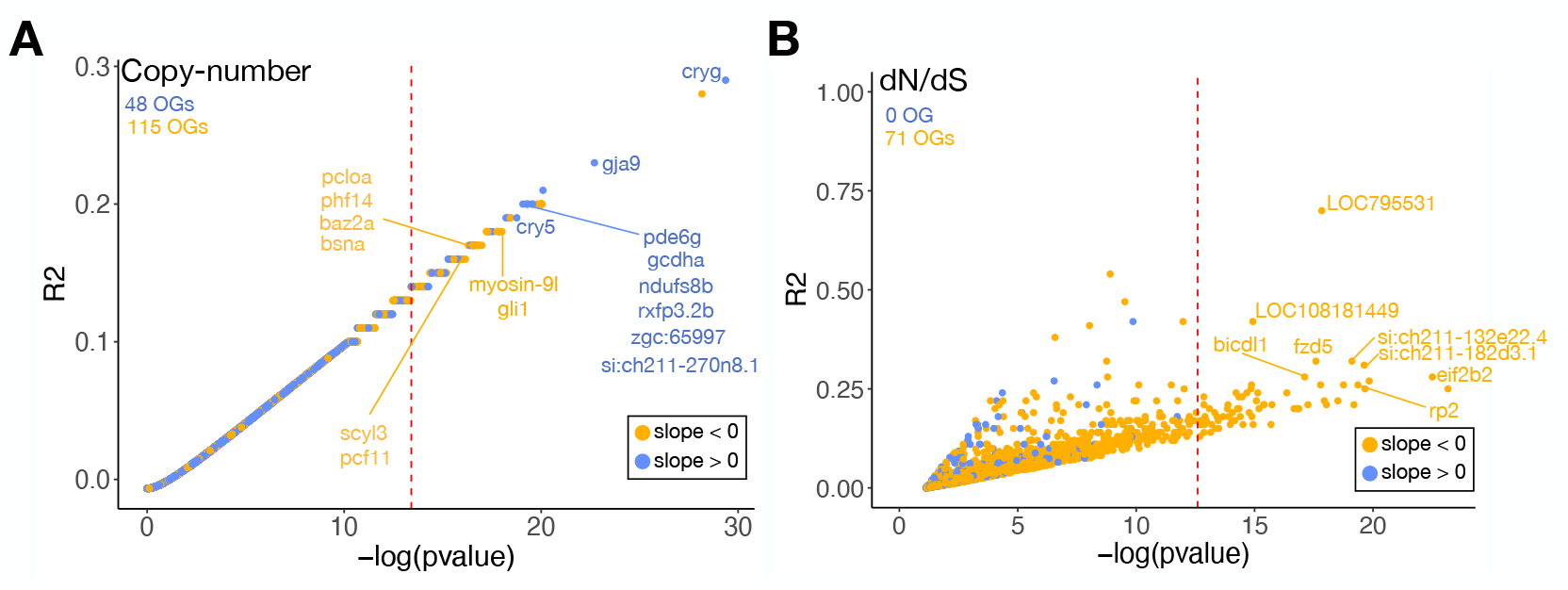
Genome-wide screen for associations between the number of non-visual opsins and (A) the number of genes in 33,018 orthogroups and (B) the mean ω per OG, for 14,740 OGs. The dashed red lines correspond to the Bonferroni significance threshold. The number of positively correlated OGs (blue) and negatively correlated OGs (yellow) are indicated for both analyses. Names of the most significant OGs are indicated. The names assigned to an OG correspond to the name of one zebrafish gene inside this OG. OGs with no zebrafish gene were not included in the enrichment analysis and are not discussed in this study.

The number of visual opsin genes correlated positively with the number of copies in 11 OGs (Fig. S46). The most significant OGs were: (i) *capn15*, known to be involved in eye defects in humans and mice (55); (ii) *crebl2*, known to be involved in retinal neovascularization (56); and (iii) *zdhhc7*, known to be expressed in the primate retina (57) and implicated in human neurological disorders (58). Six OGs were found to have higher ω-values in species with more visual opsins (Fig. S47), and four OGs had lower ω-values (Fig. S47). Among those four, the OG *oflm2* is known to be involved in vertebrate eye development (59).

### Eco-morphological factors associated with the evolution of opsin genes

We then tested for eco-morphological factors related to the number of opsin genes or their ω-values. First, we found that a species’ standard length, diet, and absolute or relative eye diameter could not predict the number of opsin genes nor their ω-values (Fig. 4). In contrast, we observed that species capable of emitting electric discharges had fewer genes in seven different non-visual opsin subfamilies (*tmt3, opn8b*, opn*7b, opn6, rrh, opn4m1/3* and *opn4x*), leading to an overall significant reduction in the number of non-visual opsins and in the total number of opsins in these species (Fig. 4, Fig. S48). This is concordant with the reduced visual sense in such species (60) and likely the result of a sensory trade-off, as these species primarily rely on electric signals, and not photon detection, to perceive their environment. This is further supported by the anatomy of their brains, which exhibit enlarged cerebella at the expense of the optic tecta (61). In addition, we observed that genes in the non-visual opsin subfamilies *tmt3, opn7b* and *opn5* show accelerated evolution in electric fish (Fig. 4, Fig. S49). Using RELAX, a method based on a branch-site model, we found that this shift in ω was due to relaxation of selection for *opn5*, which is involved in short wavelength-mediated photoentrainment of the vertebrate retina (62, 63), but increased positive selection for *tmt3*, likely to be also involved in photoentrainment (64) (Fig. S50). No significant shifts in ω were detected for *opn7b* with this method (Fig. S50).

**Figure 4.**
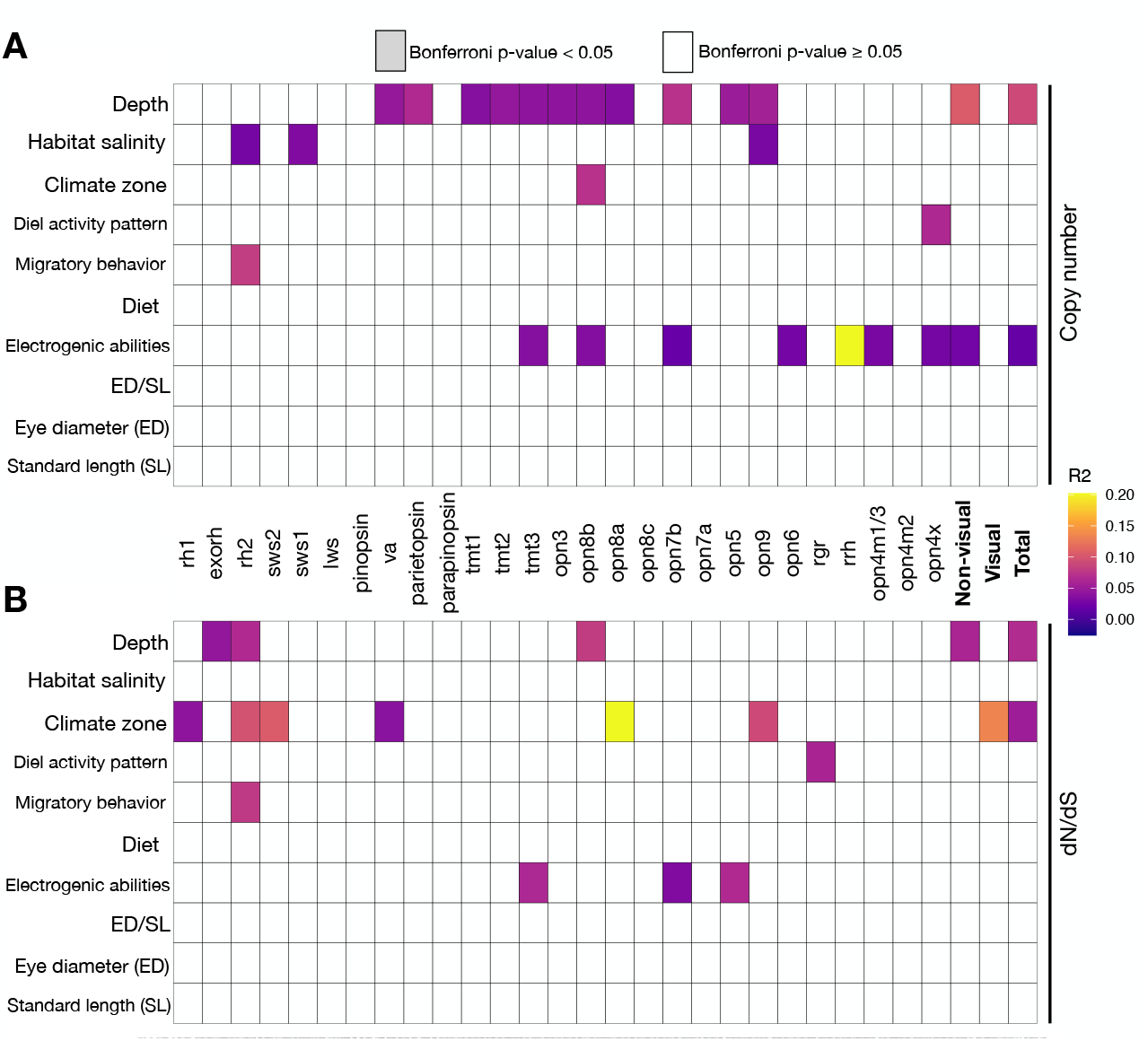
Associations between eco-morphological factors and (A) the number of genes per opsin subfamily or the total number of non-visual opsins, visual opsins and all opsins combined (“total”) and (B) the mean ω per opsin subfamily, or the mean ω of non-visual opsins, visual opsins and all opsins combined (only retaining species with at least three genes). Colored tiles correspond to significant associations (pGLS Bonferroni corrected p-value < 0.05) and the color indicates the pGLS R^2^. Details on the number of species investigated and on ecological parameters are reported in Supplementary data 1.

Next, we found that catadromous fishes had more *rh2* genes than others (Fig 4, Fig. S51). At the same time, anadromous species showed the highest ω-value for this visual opsin subfamily, followed by catadromous species (Fig. 4, Fig. S52). Diel activity patterns were associated with the number of *opn4x* genes (more copies in diurnal species; Fig. S53), in line with the role of *opn4x* in locomotor activity of *D. rerio* during its wake state (65), and with the ω-ratio of *rgr* genes (higher ω-values in crepuscular species; Fig. 4, Fig. S54).

Furthermore, we found that climate zone (temperate, sub-tropical, tropical, boreal, polar or deep-water) correlated with the number of *opn8b* genes (Fig. S55), and salinity (fresh, brackish, salt, or combinations thereof) correlated with the number of genes in one non-visual opsin subfamily, *opn9, and* two visual opsin subfamilies, *rh2* and *sws1* (with freshwater fishes having fewer genes than others, Fig. 4, Fig. S56). Climate zone was related to the mean ω-values of three non-visual opsin subfamilies (*va, opn8a* and *opn9*) and three visual opsin subfamilies (*rh1, rh2* and *sw2*), whereby, in most of these cases, polar species followed by deep-water fishes had the highest ω-ratios (Fig. S57). When considering the entire opsin gene repertoire, polar species again stand out, with much higher mean ω-values than other species (Fig. 4, Fig. S57). This signature of accelerated sequence evolution in the opsins of polar species probably reflects elevated pressures to adapt to the challenging conditions in polar waters, including variable light environments and extreme temperatures (24, 66).

The strongest predictor of the number of non-visual opsins, associated with 11 subfamilies (Fig. 4, Fig. S58), was the water depth at which a species lives. For all these subfamilies, except for *va*, bathypelagic (living below 1000 meters) species had the lowest number of genes, followed by mesopelagic species (300 - 1000 meters) (Fig. S58). We also found that depth was associated with the mean ω-values of the *exorh* and *opn8b* subfamilies, with rariphotic (150 - 300m) and mesopelagic species having slightly higher ω-ratios than others (Fig. 4, Fig. S59). Bathypelagic and mesopelagic species also had higher ω-ratios for *rh2*. Similarly, when considering the mean ω of non-visual opsin genes or the entire opsin repertoire, bathypelagic and mesopelagic species had much higher values than others (Fig. S59).

### Opsin gene expression across ray-finned fishes

Finally, we investigated the gene expression patterns of non-visual and visual opsin genes across tissues and species. We analyzed 582 publicly available transcriptomes representing up to 14 tissues from 75 ray-finned fish species (Fig. S60). As expected, visual opsins were primarily expressed in the eye (mean relative expression [RE] in the eye: 3.95% [95%CI: 2.6-5.3]; RE in other tissues usually less than 0.01%; Fig. 5). The RE of non-visual opsin genes was also highest in the eye (mean = 0.032; 95%CI: 0.023-0.041), followed by the ovary (mean of 0.012%, 95%CI: 0.01-0.014), testis (mean of 0.011%, 95%CI: 0.01-0.012), and the brain (mean of 0.01%, 95%CI: 0.01-0.011) (Fig. 5). The high RE of non-visual opsins in testis and ovaries supports recent evidence for their roles in gamete biology, including sperm thermotaxis (37) and the light-controlled maturation and release of gametes (38, 39). The lowest RE of non-visual opsins was found in the liver (Fig. 5).

**Figure 5.**
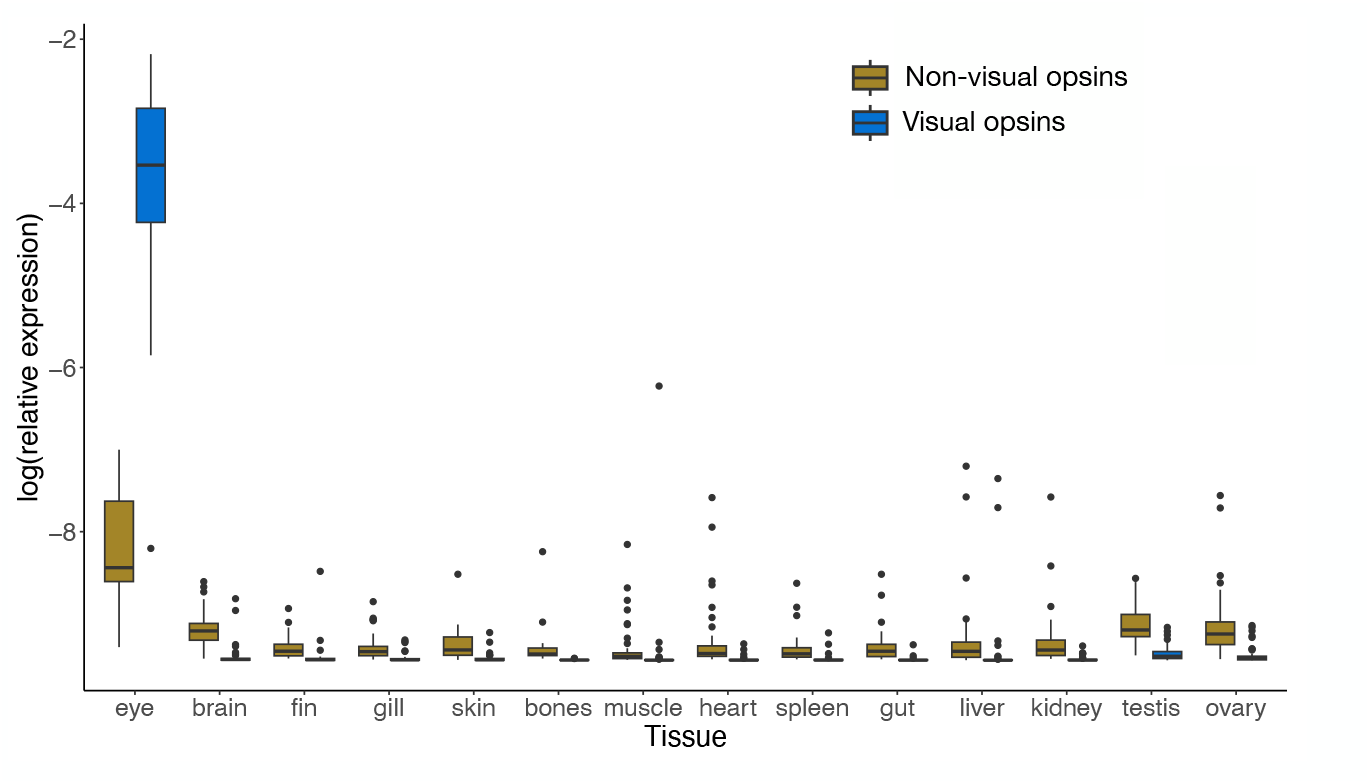
Distribution of the relative expression of visual and non-visual opsins across different tissues in ray-finned fishes.

To further investigate the RE of opsin gene subfamilies per tissue, we retained 65 samples, which had at least 1,000 reads mapped to opsins (if most opsins are expressed, this would represent approximately 30 reads per opsin on average), and we restricted our analysis to six tissues with at least four samples per tissue: brain, eye, heart, liver, ovary and testis (Fig. 6). In all species investigated, and similar to previous findings (4, 67), *rh1* represented the vast majority of visual opsin expression in the eye, ranging from 74.1% in *Chaenogobius annularis* to 99.8% in *Electrophorus electricus* (mean = 93.4%) (Fig. 6). Importantly, the RE of *rh1* was not correlated with the number of *rh1* copies in the genome (Fig. S61). Among the cone opsins, expression was dominated by *rh2* and *lws*, with a mean cone opsin RE across species of 41.6% and 43.9%, respectively. The expression of *sws2* and *sws1* was usually much lower, with a mean cone opsin RE of 14.7% and 3.7%, respectively (Fig. 6) (4, 67, 68). These proportions align well with the 1:2:2 ratio of single to double cones found in many teleost fishes (69, 70). Contrary to *rh1*, the RE of cone opsins was correlated with their copy numbers (note that *D. rerio* had an exceptionally high RE of *sws1*, Fig. S62).

**Figure 6.**
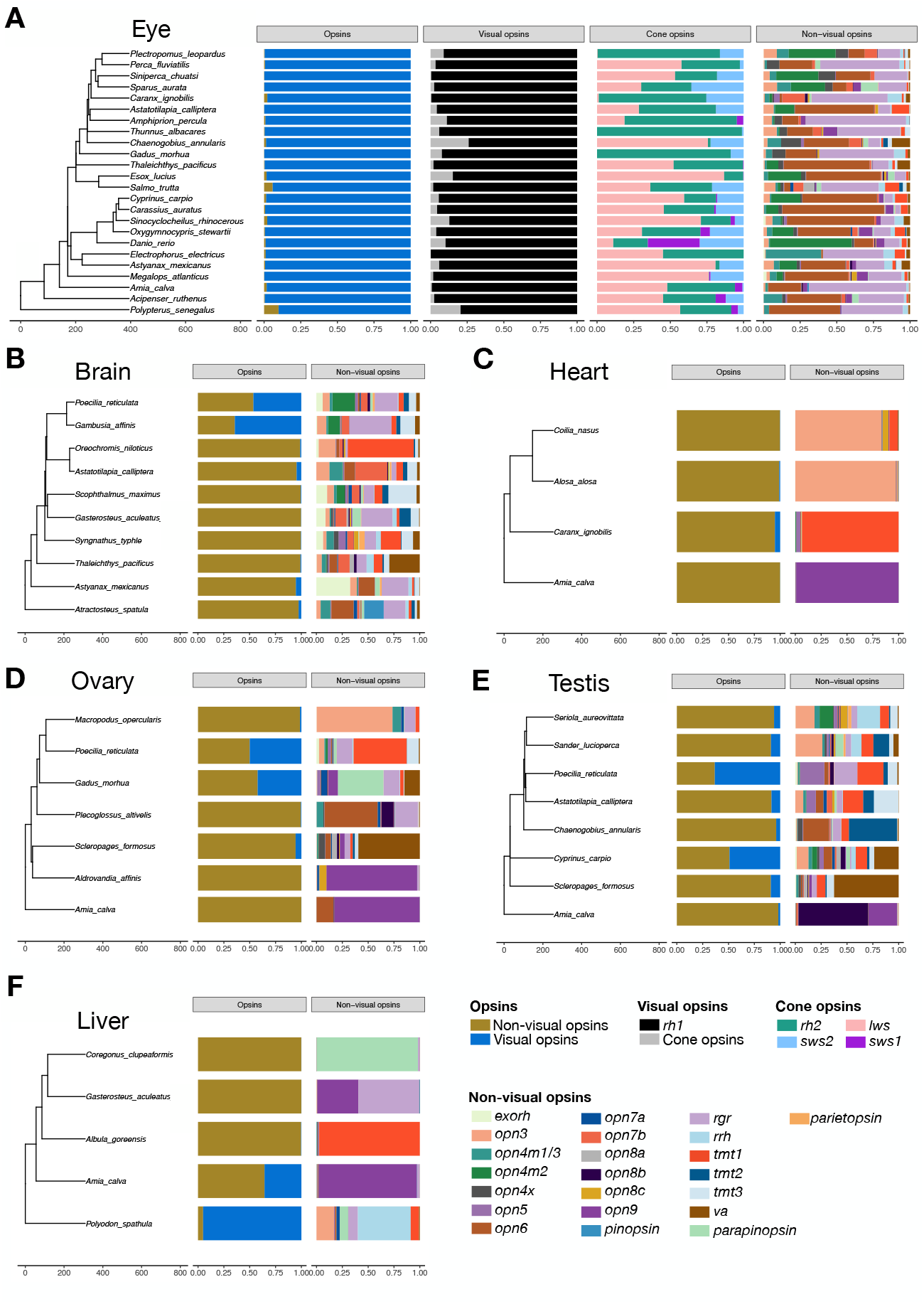
Relative gene expression levels of visual and non-visual opsins in the eye (A), brain (B), heart (C), ovary (D), testis (E) and liver (F). For the eye (A), the expression of rhodopsin and cone opsins relative to the whole visual opsin gene repertoire, and each cone opsin relative to the entire cone opsin repertoire are represented. The expression of each non-visual opsin subfamily relative to the whole non-visual opsin gene repertoire is represented for all tissues.

The expression of non-visual opsins in the eye was almost always dominated by the same three subfamilies, *opn6, rgr* and *opn4m2* (mean non-visual opsin RE of 30.3%, 27.5% and 13.9%, respectively). The high expression of *rgr* is not surprising, given that this gene is a retinal photoisomerase, which facilitates the regeneration of a chromophore used by visual opsins: the 11-*cis*-retinol (71, 72). Similarly, *opn6* has already been shown to be expressed in the photoreceptors of *D. rerio*, where it likely plays a role in their differentiation (73), and the melanopsin *opn4m2* is known to be expressed in the retina to regulate the circadian clock of fishes (74, 75). The other non-visual opsin gene subfamilies had a mean RE of 5% or less (Fig. 6, Fig. S63). There was no correlation between the RE and the gene copy number of non-visual opsins (Table S2). Overall, there was greater interspecific variability in the non-visual opsin RE in brain, ovary and testis than in the eye (Fig. 6). We also found that the diversity of expressed non-visual opsins was highest in the brain and testis (mean Shannon’s H of 2.18 and 1.96, respectively, Fig. S64), followed by the eye and the ovary (1.69 and 1.24, respectively, Fig. S70). In contrast, non-visual opsin expression diversity was much lower in the liver and the heart (0.577 and 0.308, respectively, Fig. S64), where typically one subfamily dominated the RE (>90%; Fig. 6): *opn3* in the heart of *Alosa alosa* and *Coilia nasus, tmt1* in the heart of *Caranx ignobilis* and in the liver of *Albula goreensis, opn9* in both organs of *Amia calva* and, *parapinopsin* in the liver of *Coregonus clupeaformis*. The two exceptions were the liver of *Gasterosteus aculeatus*, which had strong expression of both *rgr* and *opn9* (58.5% and 38.8%, respectively), and the liver of *Polyodon spathula*, for which *rrh* represented around half of the opsin expression, while the other half was a mix of several other subfamilies (Fig. 6).

## Discussion

The diversity of aquatic habitats and light conditions where ray-finned fishes live render this group an ideal system to scrutinize the evolution of photoreception. While previous studies mainly focused on visual opsins, and used relatively small numbers of species, we thoroughly analyzed the evolution of both visual and non-visual opsins based on 535 high-quality genome assemblies. We found evidence for co-evolution between visual and non-visual opsins and determined key ecological factors associated with the expansion or contraction of these repertoires. Specifically, we found a convergent reduction of the non-visual and visual opsin repertoire sizes in species that inhabit dim light or dark environments, such as the deep sea or caves. Similarly, our results support a sensory trade-off in electric species, commonly inhabiting turbid environments, with much fewer opsins than in other fishes. We also found that many opsin genes were rapidly evolving in polar species. Furthermore, we took advantage of our dataset to investigate genes co-evolving with the copy-number and selective pressure of opsins. Many of these genes are related to the eye and to photoreception (*e*.*g*., *pde6g, cryg, cry, capn15, oflm2, crebl2, zdhhc7, bicdl1, fzd5*, and *rp2*). We hypothesize that the remaining co-evolving genes we found are also involved in light perception or related functions, both at the image-or non-image-forming level, producing a suite of new candidates for future in-depth studies. Furthermore, we confirm that, in addition to being the primary organ for visual photoreception, the eye was also the organ with the highest non-visual opsin expression (17). Finally, the high expression levels of non-visual opsins in gametes compared to other tissues strongly support the hypothesis that non-visual opsin genes play essential roles in the reproduction of ray-finned fishes.

## Materials and Methods

### Genome data

We used genome_updater (https://github.com/pirovc/genome_updater) with the option -T “7898” to download 1,203 ray-finned fish genome assemblies available at the NCBI database as of 30 May 2023, and with the option -A 1 to retain the best assembly per species, according to four filters: (i) refseq category; (ii) assembly level; (iii) relation to type material; (iiii) release date. The completeness of each genome was then assessed using BUSCO v5.1.274 (76) with the Actinopterygii odb10 database, which includes 3,640 genes (Fig. S1). We retained 527 teleost assemblies with a BUSCO score ≥ 90% and 8 non-teleost assemblies with a BUSCO score ≥ 80%, for a total of 535 ray-finned fishes genome assemblies.

### Species tree

We extracted the protein sequences of the 2,000 most represented BUSCO genes, and for each gene, protein sequences were aligned using MUSCLE v5.1 (77). Alignments were trimmed using trimAl v1.4 (78) with the option “-automated1”. IQ-TREE2 was then used to build a maximum-likelihood tree for each gene, using the optimal model found by ModelFinder (79) and the robustness of the nodes was assessed with 1,000 ultrafast bootstraps (80). Low supported nodes (< 10% bootstrap support) were contracted using the R package ape v5.0 (81). We then used ASTRAL-III (82) to build an unrooted species tree using this set of 2,000 unrooted gene trees. The unrooted species tree was rooted and dated with IQ-TREE2 and the least square dating method (83) using three calibration dates extracted on TimeTree.org (84) : (i) 250 million years (My) between *Danio rerio* and *Megalops atlanticus*; (ii) 396 My between *Danio rerio* and *Polypterus senegalus*; (iii) 224 My between *Denticeps clupeoides* and *Gasterosteus aculeatus aculeatus*. The order of each species was extracted from the NCBI taxonomy database (85) using the R package “taxonomizr” (86).

### Opsin gene mining procedure

Known and curated *D. rerio* opsin coding sequences were extracted from previous studies (17, 35). Corresponding protein sequences were then used as queries in a blastp (87), with an e-value of 1e-5, against all annotated proteins of ten other ray-finned fish genomes : *Polypterus senegalus, Poecilia reticulata, Sebastes umbrosus, Lepisosteus oculatus, Anguilla anguilla, Takifugu rubripes, Hippocampus zosterae, Oreochromis niloticus, Gadus morhua, Gasterosteus aculeatus aculeatus*. All matching protein sequences were then used as queries in a second blastp against the UniProt database (88). All protein sequences that best matched against a known opsin gene present in the UniProt database were extracted and aligned using MUSCLE v5.1, as well as with zebrafish opsins and a non-opsin gene, *gpr52* (Gene ID: 101884270). The alignment was trimmed using trimAl and a maximum-likelihood tree was built with IQ-TREE2 using the optimal model found by ModelFinder. The tree was manually rooted using the outgroup gene *gpr52* in iTOL v5 (89). All protein sequences that clustered with zebrafish opsins were retained to build a “known-opsins database”. This database was then used as query in a tblastn against the 535 genome assemblies, with an e-value of 1e-05. We used a modified version of a previously published pipeline (90), detailed in the Extended methods, to extract opsin gene sequences from these tblastn results. A total of 17,318 complete opsins, 1,090 incomplete opsins and 2,207 pseudogenes were retrieved. By assessing our opsin gene mining results, we confirmed that the extracted sequences were of high-quality, and that the numbers of retrieved opsins were more exhaustive than previous studies or than RefSeq annotations (Extended methods). Furthermore, we show that the number of opsin genes retrieved is consistent between different assemblies of the same species, and that differences in assembly sequencing methods or release dates had no or very little impact on our results (Extended methods).

### Opsins classification and gene trees

A backbone alignment was built by aligning complete opsin protein sequences retrieved in *Lepisosteus oculatus*, using MUSCLE v5.1. MAFFT v7.467 (91) was then used to align all the other complete opsins to this backbone alignment, using the options “--add” and “--keeplength”. The alignment was trimmed to remove positions where more than 50% of sequences had a gap using trimAl. We used IQ-TREE2 to build a maximum-likelihood phylogeny from this alignment, using the optimal model found by ModelFinder and the robustness of the nodes was assessed with 1,000 ultrafast bootstraps. Genes were assigned to a subfamily based on their position on the phylogenetic tree and with the help of already classified *Danio rerio* sequences in a previous study (17). Our opsins maximum-likelihood phylogeny (Fig. S2) was similar to previously published phylogenies (17, 18). Incomplete genes and pseudogenes were then assigned to a subfamily based on their best blastx match against complete genes. For each subfamily, we aligned all retrieved protein sequences, with the addition of complete proteins of Chondrichthyes or Agnatha opsin sequence(s) of the same subfamily, or of a close subfamily, (retrieved in the NCBI non-redundant protein database) using MUSCLE v5. These alignments were trimmed using trimAl and maximum-likelihood trees were computed for each subfamily using IQ-TREE2, with the same strategy mentioned before. We also used trimAl to reverse translate these amino acid alignments into codon alignments, using the option “-backtrans”. All the subfamilies maximum-likelihood trees and alignments (DNA and proteins) can be found on FigShare. A detailed description of methods used to evaluate the dN/dS on *pinopsin* sequences can be found in Extended methods.

### Statistical analysis and graphical representations

Phylogenetic Generalized Least Squares (pGLS) analysis were conducted using the R package ‘caper’ (92). For pGLS analysis involving a categorical variable as predictor, we first removed categories represented by less than three species. Phylogenetic tree were manipulated using iTOL, taxonium (93), and the R packages ‘phytools’ (94), ‘ggtree’ (95), ‘phylobase’ (96), and ‘adephylo’ (97). Other data manipulation and graphical representations were performed using the R packages ‘dplyr’ (98), ‘tidyverse’ (99), and ‘ggplot2’ (100). Animal silhouettes were extracted from http://phylopic.org.

### Gene subfamilies dynamic and selection pressure

For each opsin subfamily maximum-likelihood (ML) gene tree, we used “ape” to remove the Chondrichthyes or Agnatha sequence(s) and to collapse nodes with a bootstrap support lower than 95%. These gene trees and the species tree were used as input in TreeRecs (46) to find the best root for each subfamily, i.e, the root that minimizes the number of duplication and loss events in the species tree. Duplications and losses events were mapped to the species tree using NOTUNG (101). Mean birth and death rates were computed for each gene subfamily using equations retrieved in a previous study (5). Due to the complete loss of some opsins subfamilies in certain lineages, for each subfamily, we excluded nodes for which the ancestor had no gene from the rates calculations, preventing their artificial decrease. Birth and death rates did not correlate with mean sequence length or the mean number of exons (Fig. S65-S66). The reconciled gene trees, as well as the ML gene trees were used as input in the standard MG94 fit program implemented in HyPhy (102) to compute maximum likelihood values of ω (= dN/dS) per branch. Branches with a saturation of synonymous or non-synonymous substitutions (dS or dN ≥1) or with too few synonymous mutations (dS < 0.01) to allow a reliable computation of ω were discarded. There was an extremely high correlation between ω-values computed on terminal branches with both trees (Pearson’s R =0.962, p-value < 2e-16, Fig. S6), but also between the mean ω-values obtained for each subfamily (Pearson’s R=0.996, p-value < 2e-16, Fig. S67). Thus, ω-values reported in this study are those obtained with the reconciled trees, but results are consistent considering ω-values obtained with the ML gene trees. Reconciled gene trees were also used as input in BUSTED (103) implemented in HyPhy. Trees were modified to assign daughter branches of duplications nodes to ‘test’ branches, while all other branches, resulting from speciation events, were assigned to ‘background’ branches. BUSTED allowed to compute a maximum-likelihood value of ω per branch type (here ‘test’ vs ‘background’) and to perform likelihood ratio test to infer if ‘test’ branches experienced significant positive selection relative to ‘background’ branches. No correlation was found between the strength of ω differences (ω_duplication_ - ω_speciation_) and the birth or death rates of subfamilies (Fig. S68). Finally, *tmt3*, opn5 and *opn7* alignments and gene trees were used as input in RELAX (104), implemented in HyPhy, with branches corresponding to electric fish opsins genes to “test” branches and all other terminal branches to “foreground”.

### Genome-wide orthologous genes identification and gene ontology analysis

4,617,897 protein sequences from the 178 annotated species of our dataset (filtered and extracted with AGAT, see Extended methods) were used as input in OrthoFinder (105), which retrieved 35,448 orthogroups (OGs), containing a total of 4,538,760 genes (98.3% of input proteins). 22 orthogroups corresponding to opsin genes (identified by BLAST against our opsin sequences dataset) were removed. For each remaining orthogroup, protein sequences were aligned using MUSCLE v5.1, and the alignment was trimmed and reverse translated using trimAl. Maximum likelihood trees were computed with IQ-TREE2, using the general amino acid replacement matrix (106) and a discrete gamma model with four rate categories (LG+G4). We then used the standard MG94 fit program implemented in HyPhy to compute maximum likelihood values of ω per branch. Again, to only keep reliable estimates of dN/dS, we discarded branches with dS < 0.01, dS ≥1 or dN ≥1. We then conducted two set of pGLS analysis: (i) between the number of opsins and the number of genes in 33,018 of these OGs which had a non-null variance, and only considering teleost species with no recent whole-genome duplication (155 species); (ii) between the number of opsins and the mean dN/dS per OG, for 14,740 OGs for which the dN/dS was available for at least 20 species, again only considering teleost with no recent whole-genome duplication. Each OG containing at least one zebrafish gene (14,451 OGs) was assigned to a unique name (107) corresponding to the zebrafish Entrez gene name. If an OG contained more than one zebrafish gene, then the gene name was chosen at random among the possible names. We then used the R package ‘gprofiler2’ (108) to conduct functional enrichment analysis. The Entrez gene names of significantly correlated OGs were used as query and, as background, we used the gene names of the 14,451 OGs for which at least one zebrafish gene was retrieved. Organism was set to “drerio” and we used three sources : GO biological process (109), Reactome (51) and KEGG (110). Finally, we retrieved a list of 53 genes involved in light-perception and 42 genes involved in the circadian clock regulation from a previous study (111). These genes corresponded to 31 and 23 OGs in our dataset respectively. Given that the number of non-visual opsins was associated to the number of genes in 48 OGs, including 4 light-perception OGs and 1 circadian clock OG, we performed 10,000 simulations where 48 OGs were drawn at random, followed by a count of the number of light-perception and circadian clock OGs. The probability (p-value) of having four light-perception OGs among the significantly associated OGs was computed as the number of simulations where four or more light-perception OGs were drawn divided by the total number of simulations (10,000). The same was performed to have the probability of picking one circadian clock OG. The same procedure was performed to assess the probability of having 1 light-perception OG among 71 OGs for which the dN/dS was negatively associated to the number of non-visual opsins.

### Ecological and morphological data

Mean depth of occurence of 399 species were extracted from previous studies (68, 112, 113). For species not present in these three studies, data were extracted from OBIS (http://www.iobis.org/) catch records using the method described in (40) [N=225], and the R package ‘robis’ (114). Finally, for species not present in previous studies and for which no catch records were found in OBIS, depth data were retrieved from fishbase (115) [N=30]. Following the methods described in a recent study (68), riverine/pond/sptring living species for which depth data were not retrieved in the various sources described above, we assigned a depth of 2 meters [N=63]. We then classified species in five depth categories (112): (*i*) shallow above 30 meters ; (*ii*) mesophotic between 30 and 150 meters; (*iii*) rariphotic between 150 and 300m; (*iv*) mesopelagic between 300 and 1000 meters; (*v*) bathypelagic below 1000 meters. The habitat salinity (fresh, salt, brackish water or a combinations thereof), trophic level, electrogenic ability, the climate zone (‘EnvTemp’), the migratory behavior (“AnaCat”) as well as the standard length and eye diameter for each species were extracted from fishbase. Diel activity patterns were extracted from a previous study (116). Diet categories (herbivore, omnivore, carnivore) were infered from the trophic level according to fishbase recommandations (https://www.fishbase.se/manual/English/fishbasethe_ecology_table.htm). Associations between opsin number or ω and ecological factors were relatively similar when considering teleosts without recent whole-genome duplication or when considering all ray-finned fishes (Fig. S69).

### Transcirptome data

RNA sequencing data for a set of 75 phylogenetically representative ray-finned fish species, from 39 different orders, were manually retrieved on the SRA website. For each species, we systematically searched for RNA data extracted from 14 different tissues: eye, liver, gut, fin, heart, skin, gill, muscle, brain, kidney, bones, spleen, testis and ovary. We only retained RNA-seq data from adult specimens and removed those from specimens treated with chemical products. A total of 799 SRA runs (accessions can be retrieved on Supplementary Data 1) were downloaded using ‘fastqer-dump’ implemented in the SRA toolkit. For each species and each tissue reads were concatenated if different sequencing runs were downloaded. Thus, a total of 601 different species/tissues combinations were retrieved (one species/tissue combination will be named sample thereafter). Reads were then trimmed and filtered using fastp (117), and aligned to their corresponding species genome assembly using STAR (118). We further removed 17 samples for which less than 70% of reads mapped to the genome and removed two samples that had less than five million mapped reads, for a final dataset of 582 samples (Fig. S70). For each species, a custom GFF3 containing retrieved opsin genes was generated by remapping opsins to their respective genome assembly, using a modified version of EXONERATE (https://github.com/hotdogee/exonerate-gff3), and using the options ‘--gff3 yes’ and ‘--showtargetgff yes’. The numbers of reads mapped to each opsins were counted with HTSeq (119). In addition, for 45 species which had an annotation file available on NCBI, the number of reads on each annotated genes were also counted with HTSeq (retaining the longest transcript per gene), by first replacing automatically annotated opsin genes by our curated opsin annotations. For each gene, we computed the reads per kilobase (RPK) by dividing the number of reads by the gene length (in kbp). A per million scaling factor was computed for each sample, as the sum of all RPK values, divided by 1,000,0000. Finally, RPK values were divided by this scaling factor to obtain transcripts per million (TPM) per gene in each sample. By using the 45 annotated species, we ensured that these public RNA samples clustered by tissue type on a PCA of TPM values for 5,782 shared OGs (Fig. S60). In addition, we observed that in most samples, there were low but significant negative correlations between genes TPM and ω-values (Fig. S71-S72). The relative expression (RE) of opsins per sample was computed as the sum of opsins TPM divided by the sum of all genes TPM. Using a linear regression (p-value < 2e-16, Fig. S73), we inferred the following relation between RE of opsins and the proportion of mapped reads that were mapped on opsins (prop_reads_):

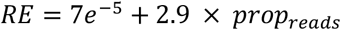

This equation was thus used to compute the relative expression of opsins for non-annotated species based on the proportion of mapped reads that were mapped on opsins.

Finally, to investigate the RE of each opsin subfamily (relative to all opsins), we directly used TPM values. For opsin subfamilies with two or more genes, TPM values of these genes were summed to obtain the TPM at the subfamily level. Shannon diversity indexes were computed using the formula described in (120).

## Acknowledgments

We would like to thank the members of the Salzburger laboratory for valuable suggestions and comments on this study. All calculations were performed at sciCORE (http://scicore.unibas.ch/), the center of scientific computing at University of Basel (with support by the SIB/Swiss Institute of Bioinformatics). This work was funded by the Swiss National Science Foundation (SNSF; grants 189970 and 208002) to W.S.

## Author Contributions

M.P. and W.S. designed this study, with input from L.G.F. and F.C. M.P. performed all data analyses. L.G.F. performed manual verification of opsin copy-numbers and sequences. M.P., L.G.F and W.S. wrote the manuscript with input from F.C.

