## Supplementary File for "Evolution of the non-visual and visual opsin gene repertoire in ray-finned fishes"

**This PDF file includes:**

Extended methods
Tables S1 to S2
Figures S1 to S73
SI References

**Other supporting materials for this manuscript include the following:**

Supplementary data 1

### Extended methods

**Opsin gene mining procedure.** Tblastn matches were extended 30,000 bp upstream and downstream and resulting non-overlapping regions were extracted. EXONERATE (1) was then used to predict opsin genes in these regions and resulting predictions were classified in four categories: (i) 'complete' if a proper coding sequence was found with a length ≥ 80% of the closest opsin length in the known-opsins database; (ii) 'pseudogene' if at least one LoF mutation was retrieved on the coding sequence; (iii) 'edge if no proper CDS and no LoF mutation was retrieved, and if the sequence was near a contig or scaffold border; (iv) 'truncated' if no proper CDS and no LoF mutation was retrieved and if the sequence was not near a contig nor scaffold border, or if the sequence contained ambiguous nucleotides. Genes falling in the edge and truncated categories are referred as 'incomplete' thereafter. All complete and incomplete predicted sequences were then translated into protein sequences and used as queries in a blastp against the UniProt database. Sequences that best-matched to an opsin gene were retained and considered as opsin sequences. The same procedure was used for pseudogenes but by using DNA sequences in a blastx against the UniProt database.

**Assessment of opsin gene mining results.** The number of opsins retrieved using our pipeline was compared to the number of opsins found on genome annotations available on NCBI. Among the 535 genome assemblies, a total of 178 had annotations. The corresponding GFF3 files were downloaded and the longest transcript per gene was retained and extracted using AGAT (2) commands "agat_sp_keep_longest_isoform.pl" and "agat_sp_extract_sequences.pl", respectively. All sequences were used as queries in a blastx against the UniProt database, and only sequences that best matched against an opsin gene were retained. Proteins were classified as complete if they had a proper coding sequence and a length ≥ 80% of the closest opsin length in our known-opsins database. Otherwise, the sequence was classified as incomplete, or as pseudogene if they were annotated as so in the GFF3 file (annotated by NCBI as "LOW QUALITY PROTEIN"). For each opsin gene retrieved in both the NCBI annotation file and by our pipeline, both DNA sequences were aligned using MUSCLE v5.1, and their percentage identity was computed using the R package "bio3d" (3). The number of opsins retrieved by our pipeline and present in NCBI annotations were highly correlated (Pearson's R=0.93, p-value < 2e-16, Fig. S3), and opsin DNA sequences of were highly similar. On a total of 6,372 opsin genes retrieved by both our pipeline and by NCBI annotations, 5,213 had a prefect DNA identity (82% of opsins) and 6,100 had more than 95% identity (96% of opsins) (Fig. S3).

We then compared the number of cone opsins (*sws1*, *sws2*, *lws*, *rh2*) we retrieved with the number of cone opsin genes reported in a previous study (4). A total of 85 species were compared. Among them, cone opsin repertoires were extracted from the exact same genome assembly in 47 cases, while in 38 cases, different genome assemblies were used. The number of cone opsin genes was equal in the two studies for 39 species, while we retrieved more cone opsins in 44 species (Fig. S3). Only two species had a repertoire of genes higher in the previous study: *Poecilia formosa* with two more *lws* genes (for a total of four *lws* against only two retrieved by our pipeline) and *Chaenocephalus aceratus* with one more *sws1* gene (no *sws1* gene retrieved for this species by our pipeline). However, another study by the same authors (5) reported the presence of only two *lws* genes in *P. formosa* genome, and only two *lws* genes are present in its annotation file (XM_007545451.2 and XM_007565629.2). *For C. aceratus*, no *sws1* gene is present on its annotation file (carefully searched by the mean of blastp, blastx and gene name), but it could be the case that this gene is present in the previous genome assembly version (GCA_900302675.1) but not in the one used in the current study (GCA_023974075.1). There was a high correlation between the number of cone opsins retrieved in both studies (Pearson's R= 0.9, p-value < 2e-16, Fig. S3).

We also manually verified the visual opsin gene sequences for 24 species which had publicly available eye transcriptomes. First, raw transcriptomic reads were aligned to the opsin CDS retrieved by the pipeline using STAR v.2.7.9a (6) and an initial screen based on raw read coverage was performed. Read coverage across the CDS was visually inspected in Geneious Prime v.2022.2.2 (Biomatters Ltd) and 18 sequences with irregular coverage were identified as candidates for manual validation following the method described in (7, 8). For manual gene extraction, the most similar published opsin CDS was obtained for each candidate using a blastn search against the NCBI nr database and selecting only results which were derived from manual methods (*e.g.,* sequences predicted by genome annotation software were discarded). Filtered raw reads were then mapped to candidate-specific references with default medium sensitivity settings (30% identity threshold) in Geneious Prime. Reads derived from the same sequence were identified by following single nucleotide polymorphisms (SNPs) across each gene with regular visual inspection for ambiguity and were extracted as paired mates to reduce sequence gaps. The extracted reads were assembled into contigs by mapping them back to the reference and extracting the consensus. Any partial CDS extractions were completed through cyclic mapping using low sensitivity (high accuracy, 100% identity threshold) settings to repeatedly extend the CDS. The manual extraction method yielded CDS which were identical to those mined using the automated pipeline.

Finally, we compared the number of non-visual opsin genes we retrieved with a previous study (9). On a total eight species compared, the numbers of non-visual opsins were equal in three cases, while we retrieved more non-visual opsins in five cases. Nevertheless, there was a good correlation between the numbers of non-visual opsin genes in the two studies (Pearson correlation: R=0.75, p-value=0.03, Fig. S3).

**Impact of the genome assembly quality and version on the number of opsins.** We evaluated if the number of opsin genes retrieved was dependent on the assembly quality (chromosome or scaffold level), on the sequencing technology or on the year of release. The sequencing technology was set to “long reads” if the genome was sequenced with MinION, Nanopore or PacBio, and to “short reads” if it was only sequenced with Illumina HiSeq, Illumina NovaSeq or BGISEQ-500. We observed that there was a similar number of visual opsins in chromosome and in scaffold level assemblies (pGLS p-value = 0.48, mean of 7.18 and 7.02 genes, respectively, Fig. S4), and a slightly higher number of non-visual opsins in chromosome-level assemblies than in scaffold-level assemblies (pGLS p-value = 0.02, mean of 25.8 and 22.3 genes, respectively). The number of visual and non-visual opsins were similar in long-reads and short-reads assemblies (pGLS p-value = 0.054 and 0.39, respectively, Fig. S4). The release date of the genome assembly also had no impact of the number of visual nor non-visual opsins (pGLS p-value = 0.5192 and 0.2113, respectively, Fig. S4)

We then assessed the variation in the number of visual and non-visual opsins across different genome assemblies of the same species. We used genome_updater to find species in our genome dataset which had another genome assembly in NCBI (called alternative assemblies thereafter). Among the 535 species of our dataset, 90 species had one other assembly, and 47 had two or more other assemblies, for a total of 222 alternative assemblies. We then used BUSCO v5.1.274 with the Actinopterygii odb10 database to retain 142 alternative assemblies with a BUSCO score ≥ 90% (Fig. S5), representing 84 species (Supplementary data 1). Opsin genes in these alternative genome assemblies were extracted following the procedure described above, and extracted genes were classified based on their best blastp match against our database composed of 17,318 complete opsin genes. For each alternative assembly, we then computed the absolute difference in the number of visual opsins and in the number of non-visual opsins retrieved in comparison to the main assembly used in this study. We found a very low mean absolute difference of the number of visual opsins 0.4 (min = 0 ; max = 2, Fig. S5), while it was slightly higher, 1.6, for the number of non-visual opsins (min = 0 ; max = 7, Fig. S5). Furthermore, there was a very strong correlation between the number of visual opsins retrieved in the alternative assemblies and the assemblies used in this study (Pearson's R = 0.96 , p-value < 2e-16, Fig. S5), and the slope of the linear regression was almost equal to 1 (Number of visual opsins in the alternative assembly = 0.2 + 0.97 * Number of visual opsins in the main assembly, Fig. S5). The same was true for the number of non-visual opsins (Pearson's R = 0.97 , p-value < 2e-16 , Number of non-visual opsins in the alternative assembly = -0.43 + 0.98 * Number of non-visual opsins in the main assembly, Fig. S5). Thus, we expect the results of this study to be robust and only slightly impacted by the genome assemblies versions used (provided that these assemblies have a BUSCO score greater than or equal to 90%).

***Pinopsin* gene evolution.** All complete *pinopsin* genes and pseudogenes retrieved in ray-finned fishes were aligned using MACSE (10), which can handle premature stop codons and frameshifts. Because the *pinopsin* pseudogene retrieved in the three species of the genus *Ilyophis* were identical, only one was retained. The alignment was visualized using AliView (11). We extracted the regions around the *pinopsin* gene (100,000bp upstream and downstream) in species which had a NCBI annotation (*Erpetoichthys calabaricus*, *Polypterus senegalus*, *Polyodon spathula*, *Atractosteus spatula*, *Lepisosteus oculatus*, *Megalops cyprinoides*, *Megalops atlanticus*) and all genes present in these regions were extracted. Genes were defined as orthologous if they were in the same orthogroup found by OrthoFinder (as described above). Regions, genes and links between orthologous genes were visualized using the R package "gggenomes" (12).

To assess if the complete *pinopsin* gene found in *M. cyprinoides* was under negative selection, positive selection or under relaxed selection, we used the *pinopsin* gene tree and codon alignment as input in PAML (13) to compute maximum likelihood estimates of ω (= dN/dS). Three branch models were tested: (i) a free-ratio model assuming a different ω per branch; (ii) a one-ratio model where all branches are assumed to evolve under the same ω; (iii) a two-ratio model where the *M. cyprinoides pinopsin* evolves under $\omega_1$, while all other branches evolve under $\omega_2$. Models were compared with the mean of likelihood ratio tests and p-values were computed based on a $\chi 2$-distribution. We also used RELAX implemented in HyPhy, with *M. cyprinoides* as the test branch, and all other branches as foreground.

We found that the *M. cyprinoides pinopsin* evolved under negative selection, with a ω (= dN/dS) similar to the *pinopsin* of non-teleost species (ω = 0.057) (Fig. S38). We also excluded the possibility of a lateral transfer or of a contamination, as the phylogeny of *pinopsins* is congruent with the species phylogeny (Fig. S26), and *pinopsin* was located next to the same genes in *M. cyprinoides* and *L. oculatus* (Fig. S38). We also retrieved *pinopsin* pseudogenes in *Megalops atlanticus* and in the Anguilliformes species of the genus *Ilyophis*. While the *pinopsin* in *Ilyophis* spp. was highly degenerated, with only 2 remaining exons (out of 5) and two premature stop codons, the *pinopsin* gene of *M. atlanticus* seems to have been lost recently, with only one frameshift retrieved (Fig. S38). Thus, the evolutionary history of *pinopsin* seems much more intricate than previously thought, with at least three independent losses: (i) before the MRCA of Clupeocephalans; (ii) before the MRCA of Osteoglossiformes; (iii) before the MRCA of Albuliformes, Notacanthiformes and Anguilliformes.

**A**

| term | source | name | pvalue | gene number |
|---|---|---|---|---|
| R-DRE-112316 | REAC | Neuronal System | 0.0016 | 5 |
| R-DRE-112315 | REAC | Transmission across Chemical Synapses | 0.0054 | 4 |
| R-DRE-112314 | REAC | Neurotransmitter receptors and postsynaptic signal transmission | 0.0397 | 3 |

**B**

| term | source | name | pvalue | gene number |
|---|---|---|---|---|
| R-DRE-174362 | REAC | Transport and synthesis of PAPS | 0.0166 | 2 |
| R-DRE-4086398 | REAC | Ca2+ pathway | 0.0193 | 3 |
| KEGG:00500 | KEGG | Starch and sucrose metabolism | 0.0271 | 2 |
| R-DRE-3858494 | REAC | Beta-catenin independent WNT signaling | 0.0397 | 4 |

**Supplementary Table 1.** Functional enrichment of (A) OGs for which the number of genes was positively correlated with the number of non-visual opsin and (B) OGs for which the mean ω was negatively correlated with the number of non-visual opsins. 'P-value' corresponds to the corrected p-value computed with the g:SCS method, and 'gene number' corresponds to the number of gene present in the gene list.

### A - eye

| subfamily | Pearson's R | p-value |
|---|---|---|
| exorh | 0.274 | 0.19 |
| opn3 | -0.25 | 0.24 |
| opn4m1_3 | 0.198 | 0.35 |
| opn4m2 | 0.505 | 0.012 |
| opn4x | 0.25 | 0.24 |
| opn5 | -0.022 | 0.92 |
| opn6 | 0.215 | 0.31 |
| opn7a | 0.079 | 0.71 |
| opn7b | 0.212 | 0.32 |
| opn8a | 0.23 | 0.28 |
| opn8b | 0.18 | 0.4 |
| opn8c | 0.509 | 0.011 |
| opn9 | 0.158 | 0.46 |
| parapinopsin | 0.236 | 0.27 |
| parietopsin | 0.065 | 0.76 |
| rgr | -0.05 | 0.82 |
| rrh | 0.358 | 0.086 |
| tmt1 | 0.331 | 0.11 |
| tmt2 | 0.247 | 0.25 |
| tmt3 | 0.099 | 0.64 |
| va | 0.041 | 0.85 |
| pinopsin | 0.614 | 0.0014 |

### B - brain

| subfamily | Pearson's R | p-value |
|---|---|---|
| exorh | 0.234 | 0.51 |
| opn3 | -0.38 | 0.28 |
| opn4m1_3 | -0.141 | 0.7 |
| opn4m2 | 0.332 | 0.35 |
| opn4x | 0.223 | 0.54 |
| opn5 | 0.291 | 0.42 |
| opn6 | 0.735 | 0.015 |
| opn7a | 0.018 | 0.96 |
| opn7b | -0.548 | 0.1 |
| opn8a | 0.62 | 0.056 |
| opn8b | -0.091 | 0.8 |
| opn8c | 0.133 | 0.72 |
| opn9 | 0.615 | 0.058 |
| parapinopsin | 0.22 | 0.54 |
| parietopsin | 0.412 | 0.24 |
| rgr | -0.08 | 0.83 |
| rrh | 0.075 | 0.84 |
| tmt1 | 0.382 | 0.28 |
| tmt2 | -0.004 | 0.99 |
| tmt3 | 0.17 | 0.64 |
| va | 0.017 | 0.96 |
| pinopsin | 1 | <2e-16 |

**Supplementary Table 2**. Relationship between the relative expression of non-visual opsins relative to their gene copy number (relative to the entire non-visual opsin gene repertoire), for (A) eye and (B) brain. For each subfamily, the Pearson correlation coefficient and p-value are indicated. The only significant associations are retrieved for *opn4m2* and *opn8c* in the eye, opn*6* in the brain and *pinopsin* in both tissues.

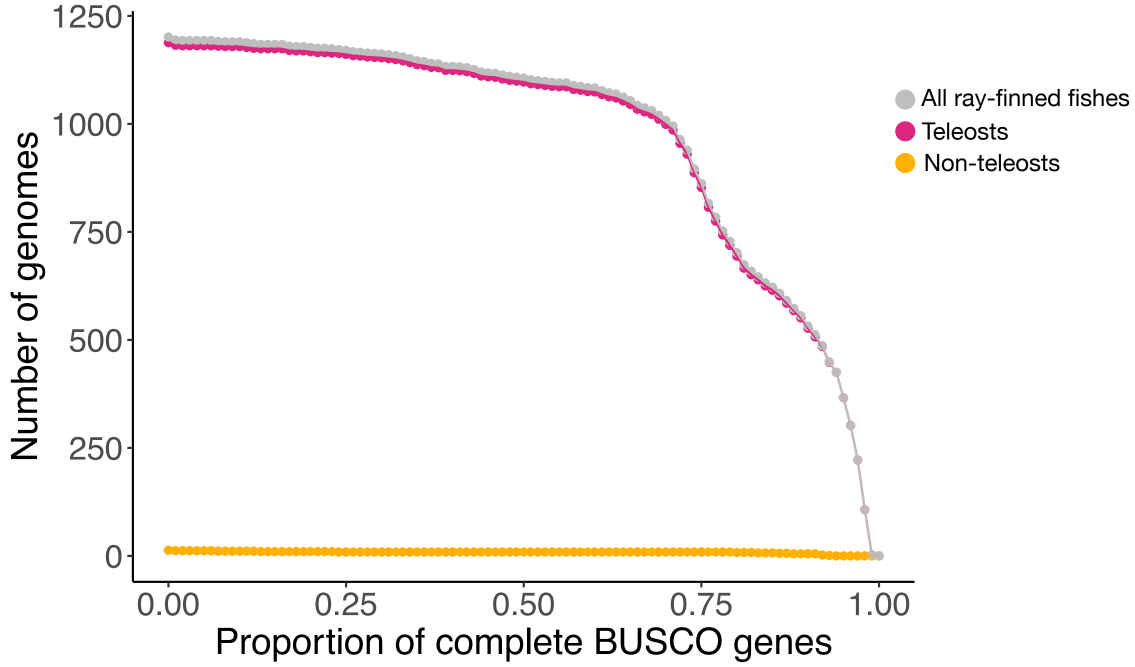


**Supplementary Figure 1.** BUSCO completeness assessment. Number of genomes in relation to the completeness threshold (proportion of complete BUSCO genes). We retained 527 teleost assemblies with a BUSCO score ≥ 90% and 8 non-teleost assemblies with a BUSCO score ≥ 80%. BUSCO raw results can be found in Supplementary Data 1.

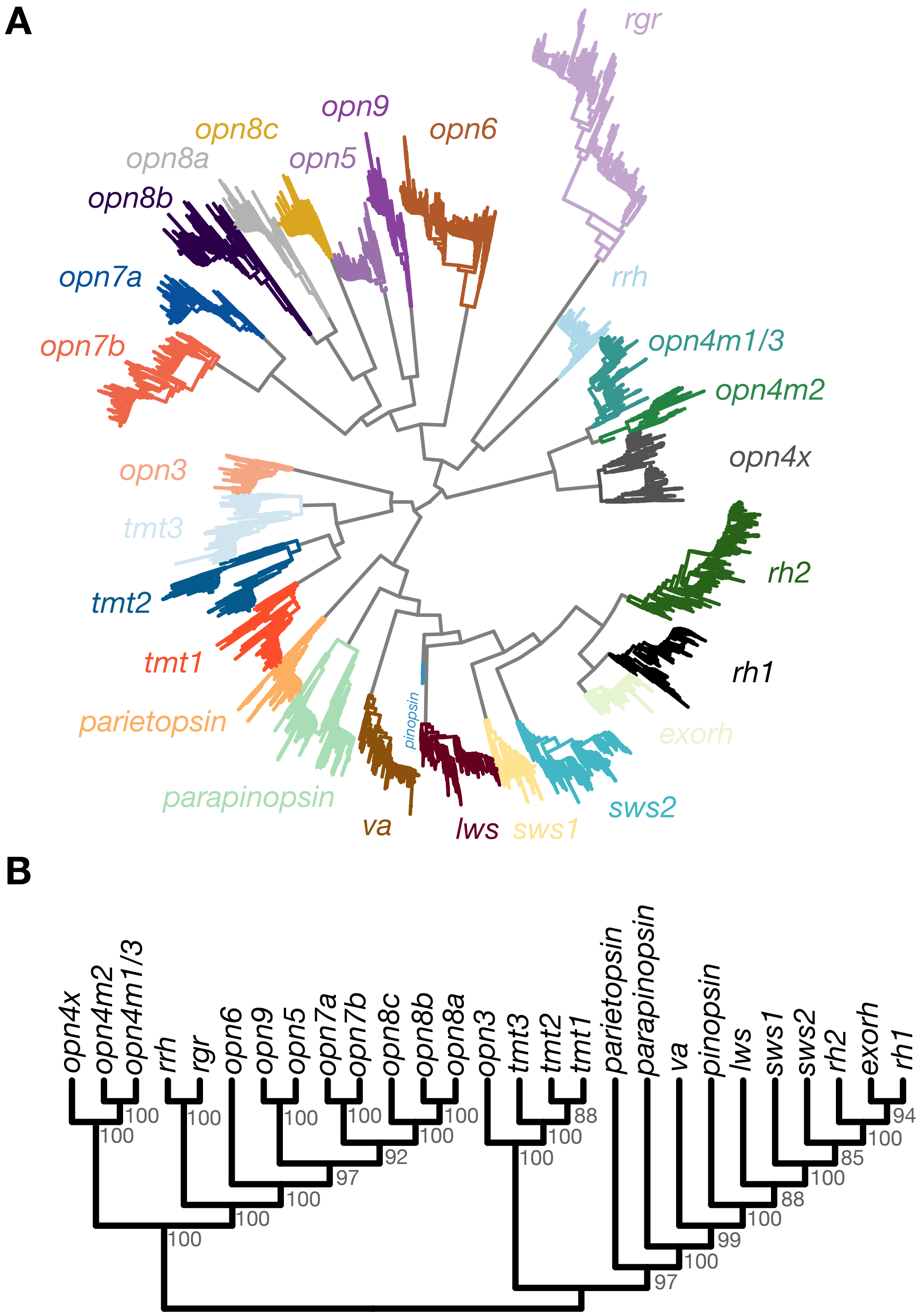


**Supplementary Figure 2.** Opsin gene phylogeny. (A) Maximum likelihood tree of 17,318 ray-finned fish opsin genes (535 species). The tree was computed with IQ-TREE and visualized using ggtree. (B) Clade tree of opsin genes generated from (A). Bootstrap values computed with ultra-fast bootstraps are indicated at each node.

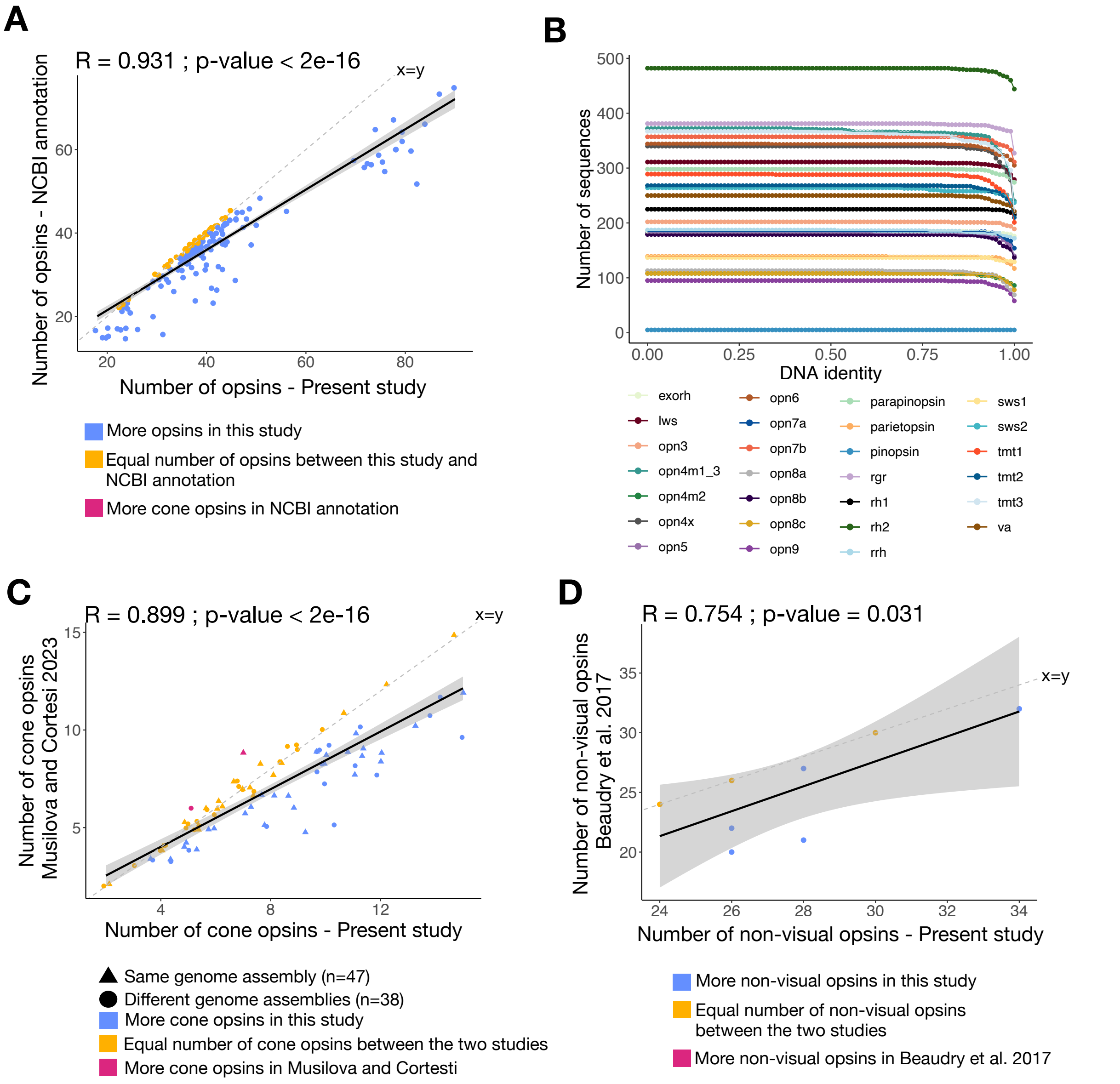


**Supplementary Figure 3**. Opsin mining procedure evaluation. (A) Comparison of the number of opsin genes retrieved in our study with the number of opsin genes annotated on NCBI for the same genome assemblies (N=178). (B) Number of opsin genes in relation to the DNA identity proportion between the NCBI annotated version and our annotation. Opsins are colored according to the subfamily. (C) Comparison of the number of cone opsins retrieved in our study with the number of cone opsins reported in Musilova and Cortesi 2023 (4). The shape of points represents if the same or different genome assemblies were used in the two studies. (D) Comparison of the number of non-visual opsins retrieved in our study with the number of non-visual opsins reported in Beaudry et al. 2017 (9). Pearson correlation coefficients and p-values are indicated for each comparison. The solid black lines represent the linear regressions, and the grey areas represent confidence intervals. Dashed lines indicate slope = 1 (x=y).

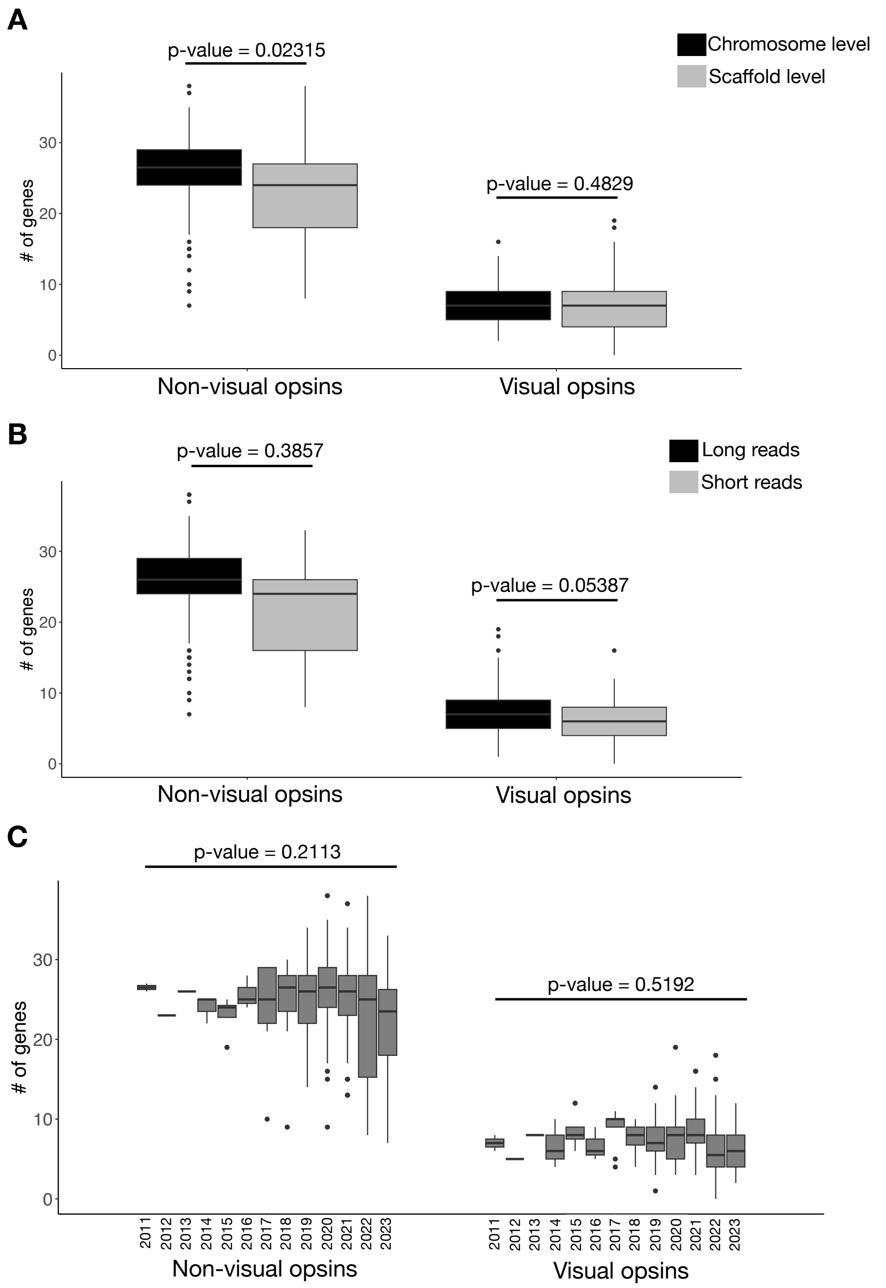


**Supplementary Figure 4.** Impact of assembly quality and release date on the number of opsin genes. Evaluation of the impact of (A) the assembly level, (B) the sequencing technology, and (C) the assembly release date on the number of non-visual and visual opsin genes. For each comparison, the pGLS p-value is reported. Only the assembly level had an impact on the number of opsin genes retrieved, with slightly fewer non-visual opsins retrieved from scaffold level assemblies than chromosome level assemblies.

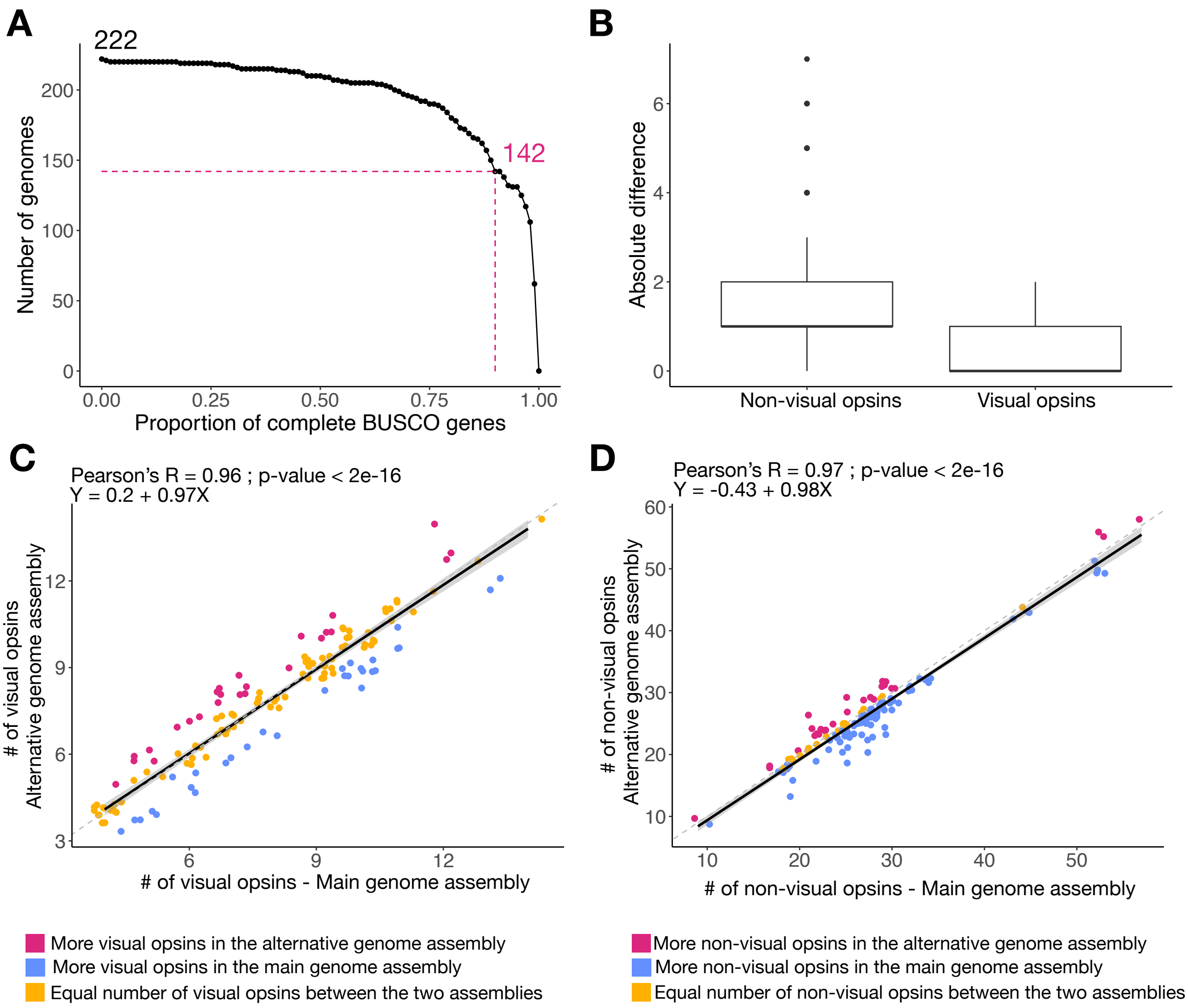


**Supplementary Figure 5.** Variation in the number of opsin genes between genome assemblies of the same species. (A) Number of alternative genome assemblies in relation to the completeness threshold (proportion of complete BUSCO genes). We retained 142 alternative assemblies with a BUSCO score ≥ 90% (Supplementary File 1). (B) Absolute difference between the number of non-visual or visual opsin genes from the main genome assembly used in this study and the number retrieved from the alternative genome assemblies for the same species. (C) Comparison of the number of visual opsin genes retrieved from the main assembly and from the alternative assemblies. (D) Comparison of the number of non-visual opsin genes retrieved from the main assembly and from the alternative assemblies. Pearson correlation coefficients, p-values and linear regression equations are indicated for each comparison. The solid black lines represent the linear regressions, and the grey areas represent confidence intervals. Dashed lines indicate slope = 1 (x=y).

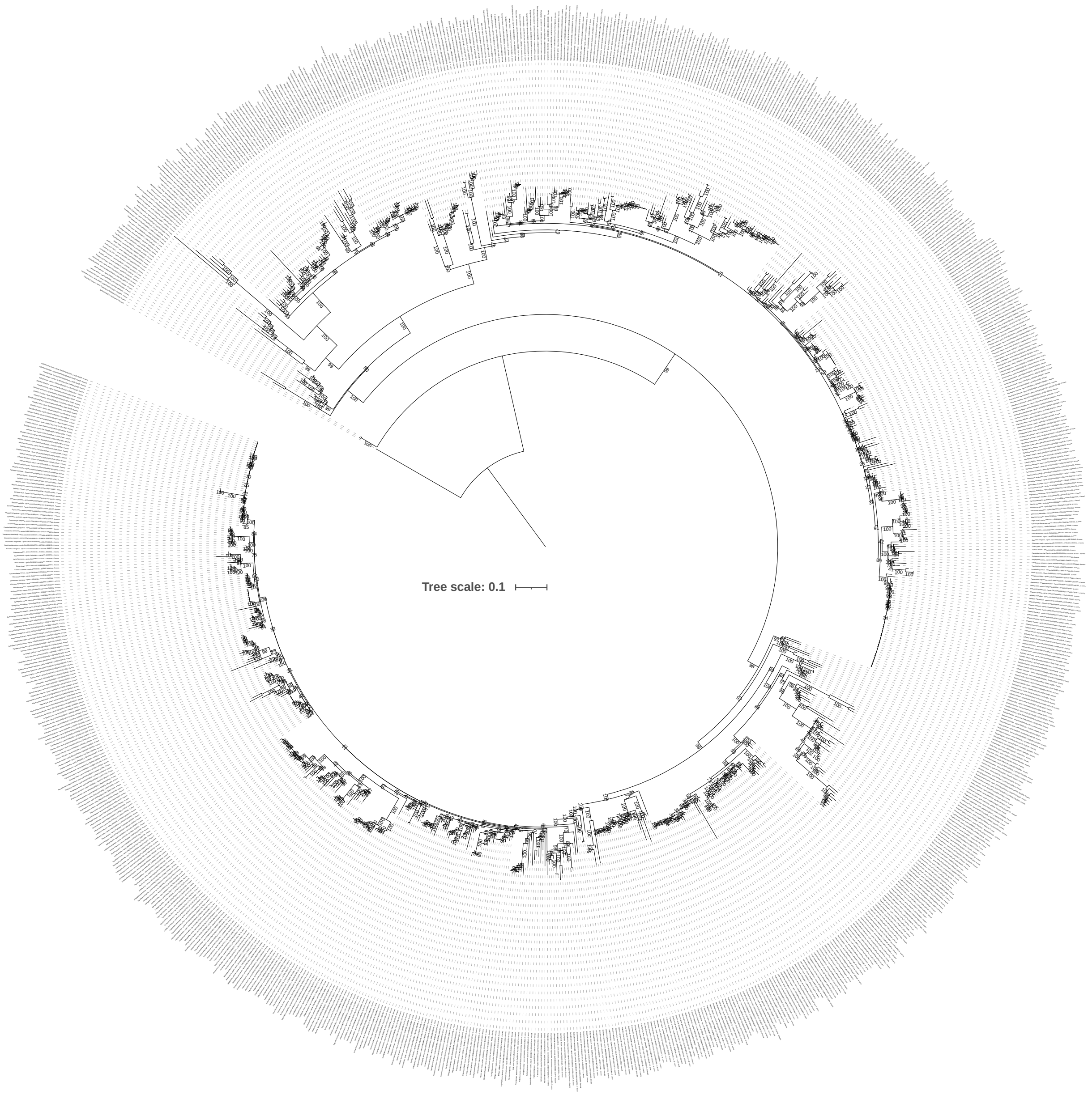


**Supplementary Figure 6.** Maximum likelihood phylogeny of *opn4x* genes. The tree was computed using the optimal model, as determined by ModelFinder (JTT+F+R9). The robustness of the nodes was evaluated with 1,000 ultrafast bootstraps and these bootstrap values are indicated at each node. The tree was rooted using the *Callorhinchus milii opn4x* gene (NP_001279400.1).

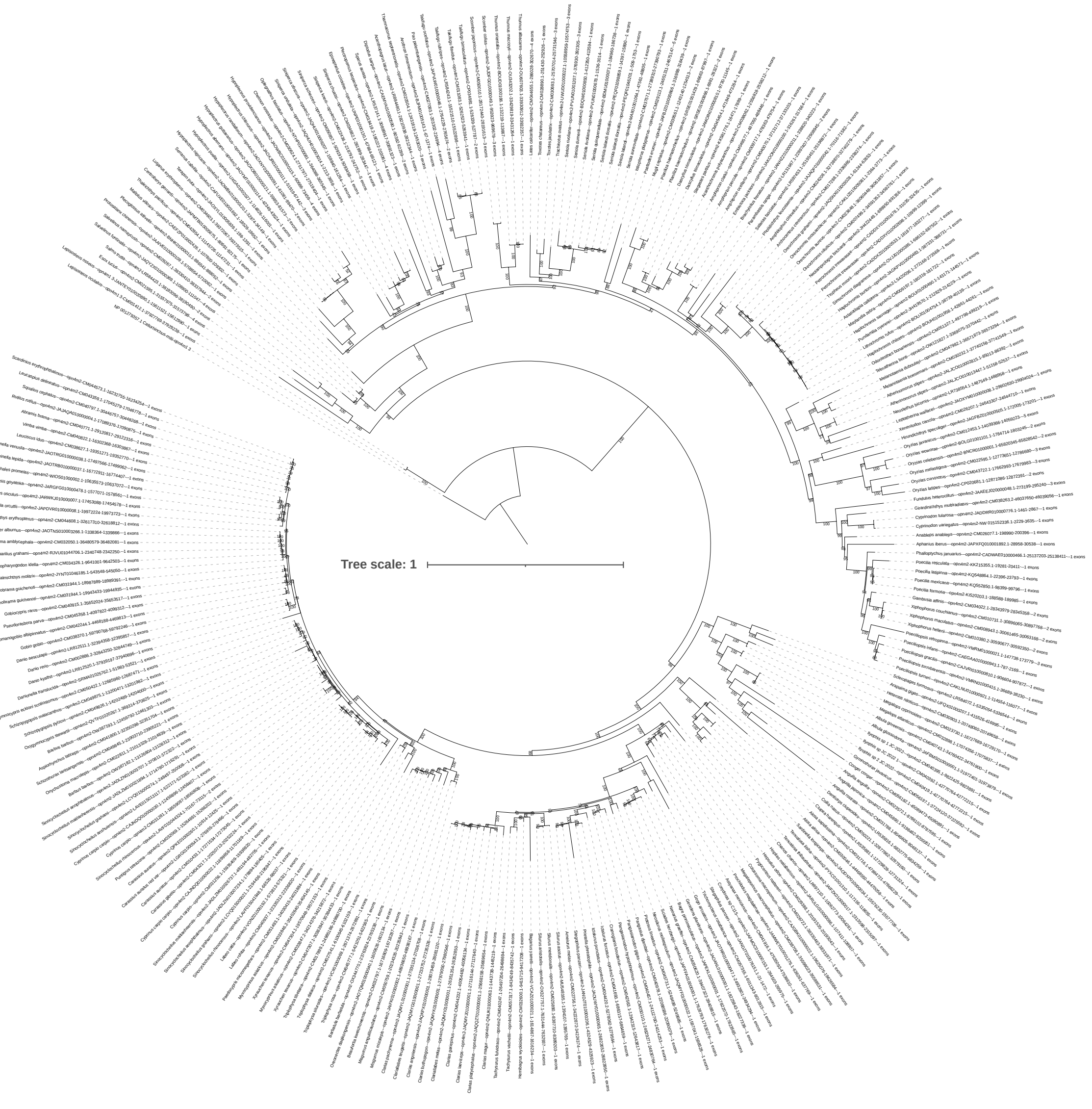


**Supplementary Figure 7.** Maximum likelihood phylogeny of *opn4m2* genes. The tree was computed using the optimal model, as determined by ModelFinder (JTT+F+R6). The robustness of the nodes was evaluated with 1,000 ultrafast bootstraps and these bootstrap values are indicated at each node. The tree was rooted using the *Callorhinchus milii opn4m1/3* gene (NP_001279357.1).

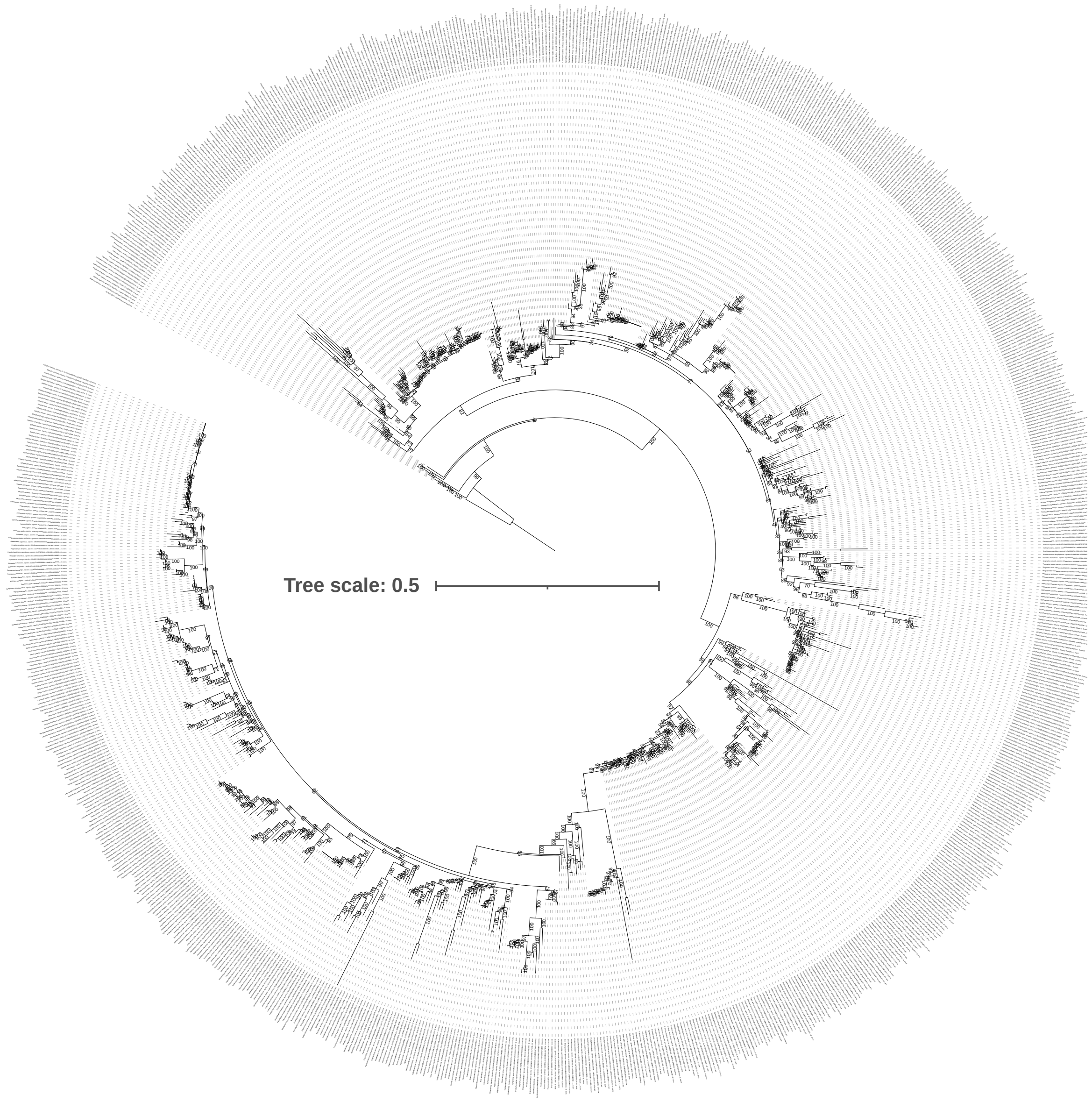


**Supplementary Figure 8.** Maximum likelihood phylogeny of *opn4m1/3* genes. The tree was computed using the optimal model, as determined by ModelFinder (JTT+R8). The robustness of the nodes was evaluated with 1,000 ultrafast bootstraps and these bootstrap values are indicated at each node. The tree was rooted using the *Callorhinchus milii opn4m1/3* gene (NP_001279357.1).

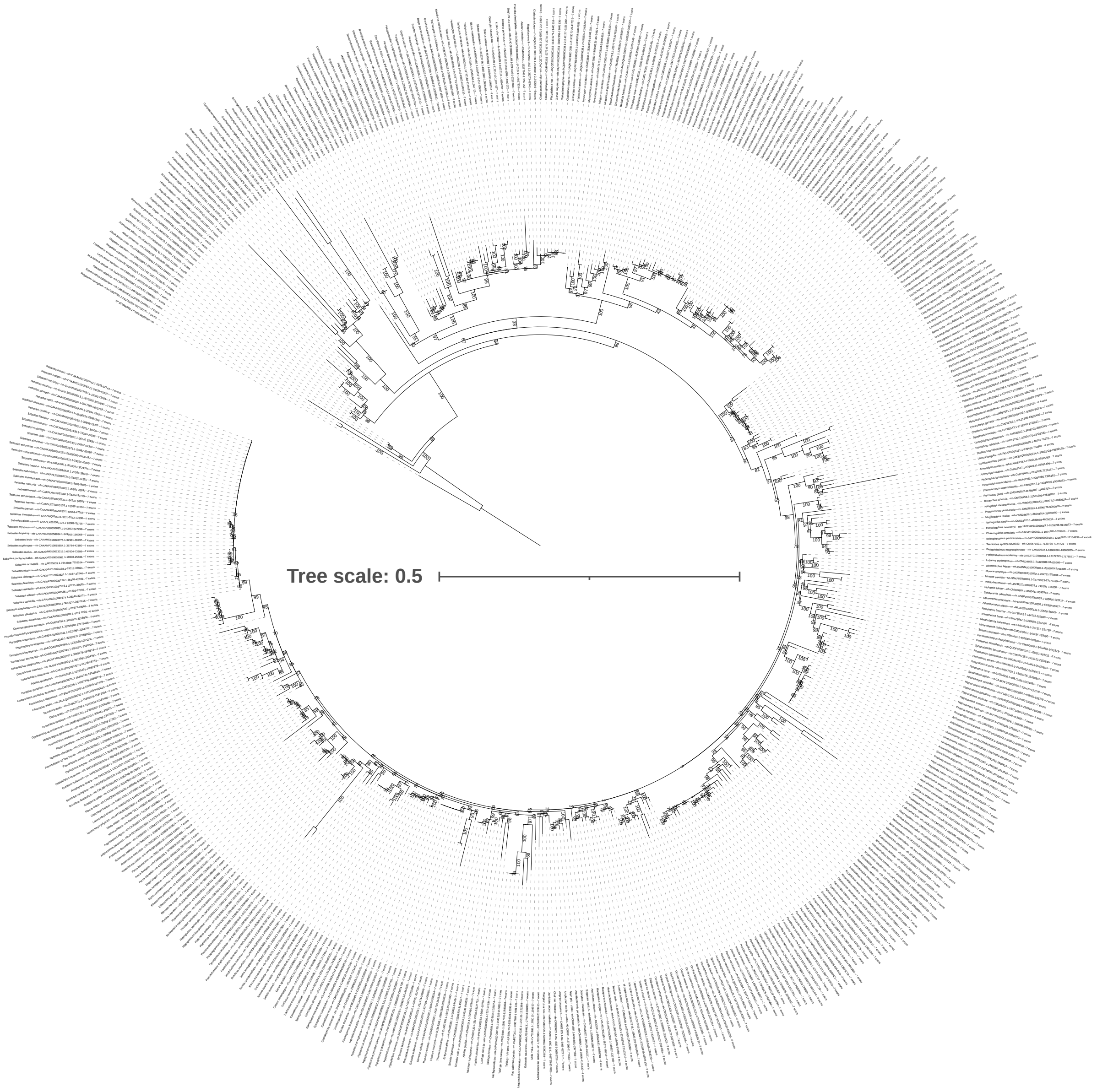


**Supplementary Figure 9.** Maximum likelihood phylogeny of *rrh* genes. The tree was computed using the optimal model, as determined by ModelFinder (JTT+R5). The robustness of the nodes was evaluated with 1,000 ultrafast bootstraps and these bootstrap values are indicated at each node. The tree was rooted using the *Pristis pectinata rrh* gene (XP_051866389.1).

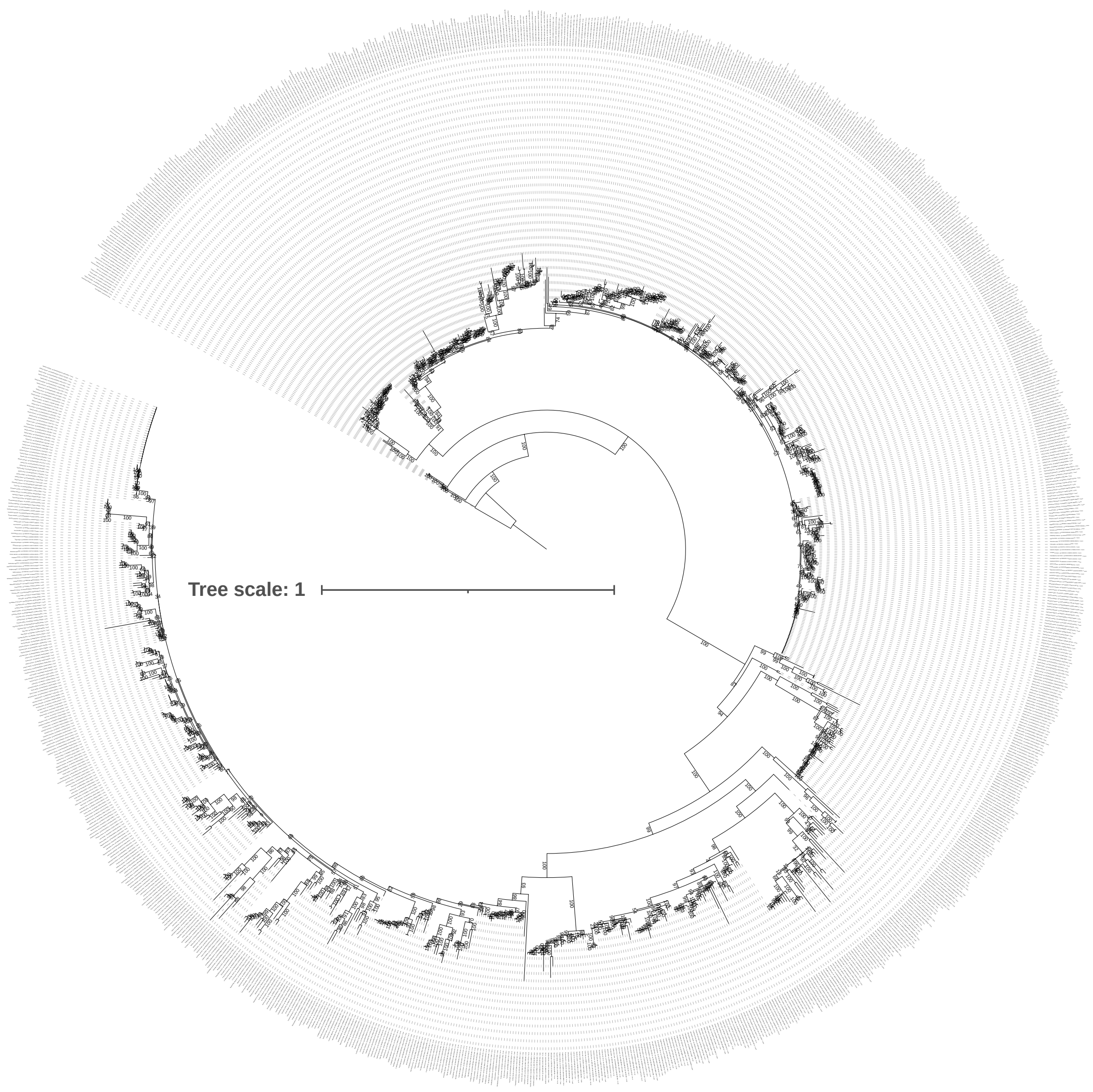


**Supplementary Figure 10.** Maximum likelihood phylogeny of *rgr* genes. The tree was computed using the optimal model, as determined by ModelFinder (JTT+F+R6). The robustness of the nodes was evaluated with 1,000 ultrafast bootstraps and these bootstrap values are indicated at each node. The tree was rooted using the *Callorhinchus milii rgr* gene (XM_007898380.1).

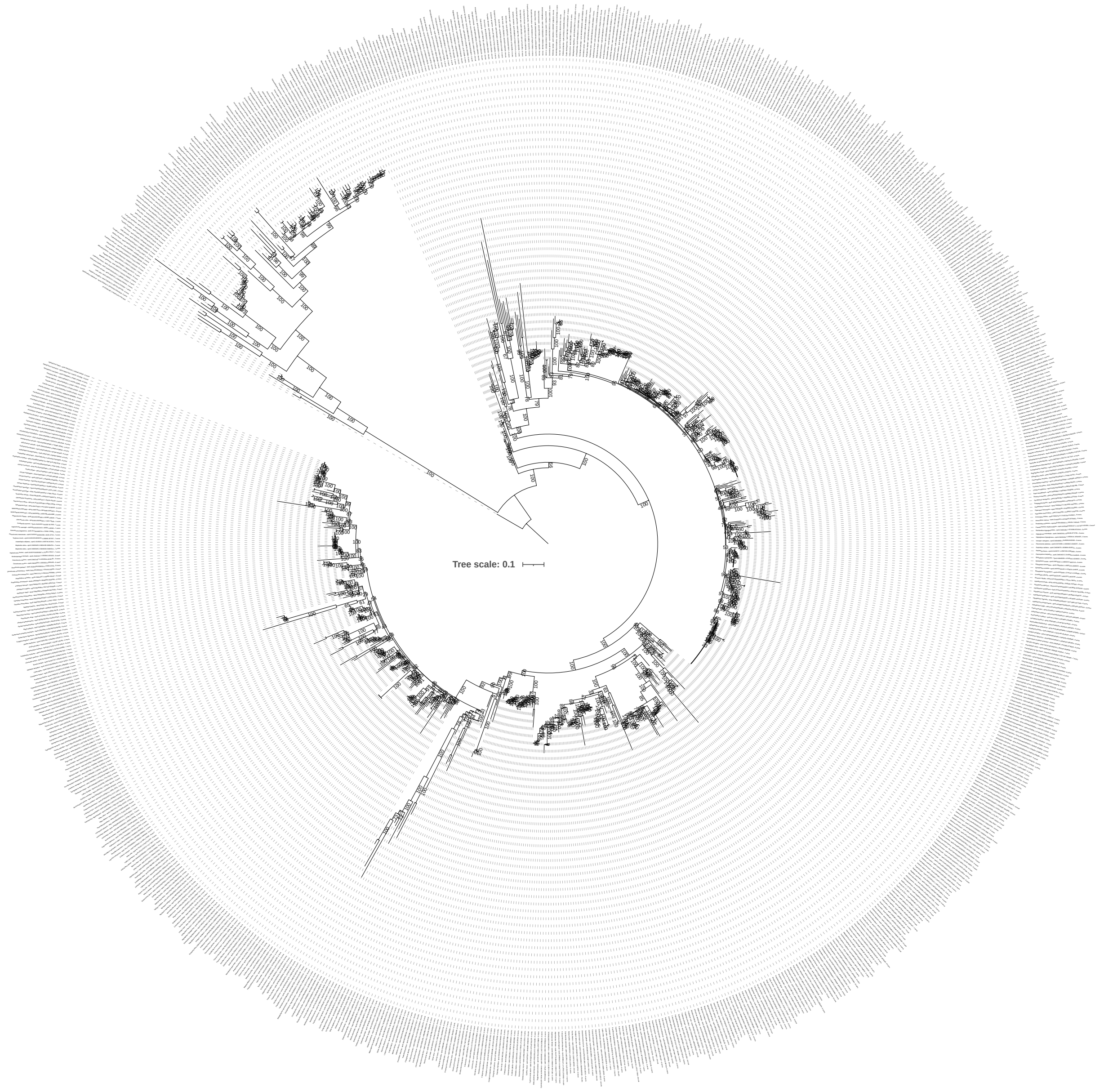


**Supplementary Figure 11.** Maximum likelihood phylogeny of *opn6* genes. The tree was computed using the optimal model, as determined by ModelFinder (JTT+R8). The robustness of the nodes was evaluated with 1,000 ultrafast bootstraps and these bootstrap values are indicated at each node. The tree was rooted using the *Callorhinchus milii opn6* gene (XP_007887390.1).

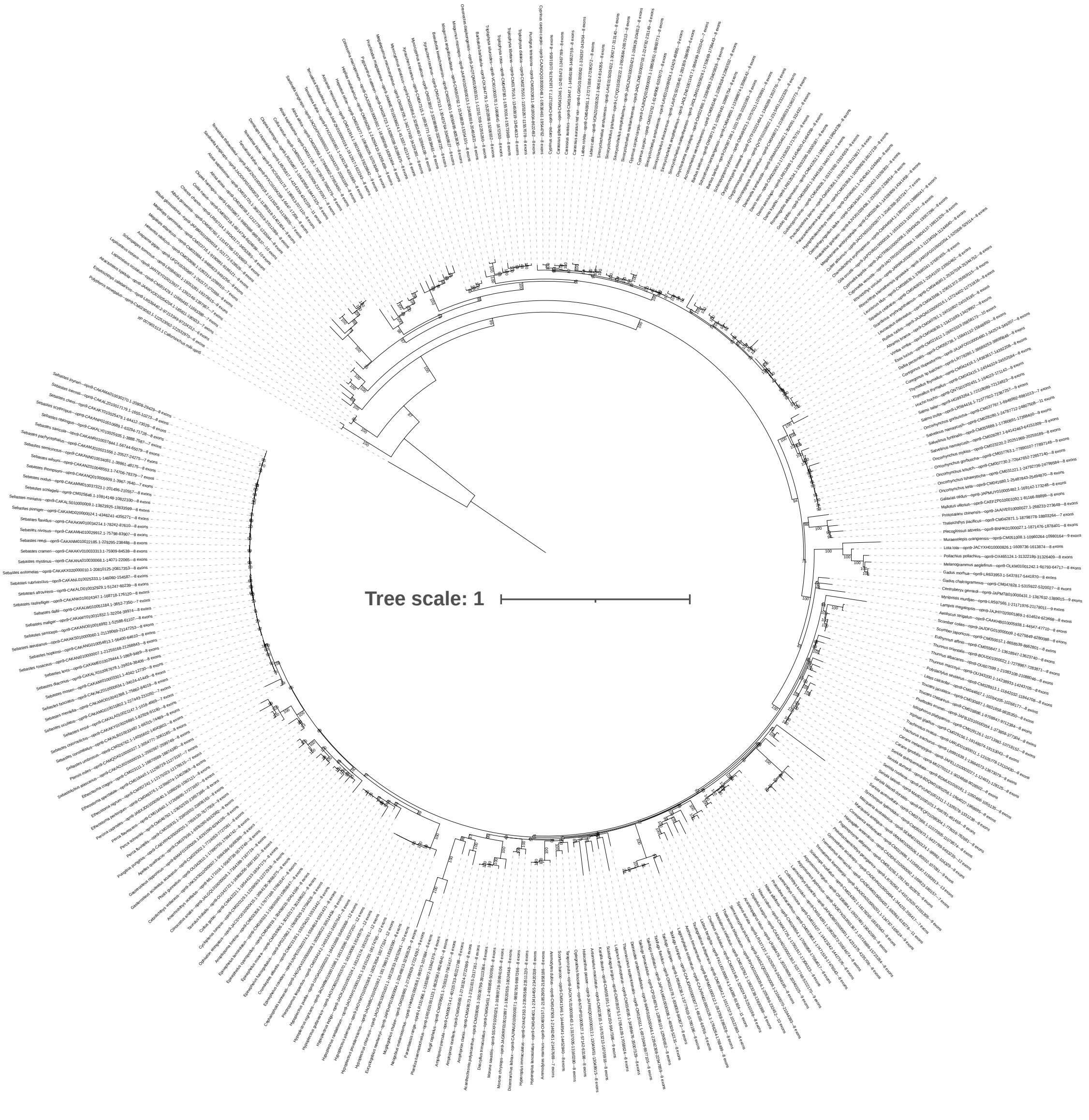


**Supplementary Figure 12.** Maximum likelihood phylogeny of *opn9* genes. The tree was computed using the optimal model, as determined by ModelFinder (JTT+F+R6). The robustness of the nodes was evaluated with 1,000 ultrafast bootstraps and these bootstrap values are indicated at each node. The tree was rooted using the *Callorhinchus milii opn5* gene (XP_007901513.1).

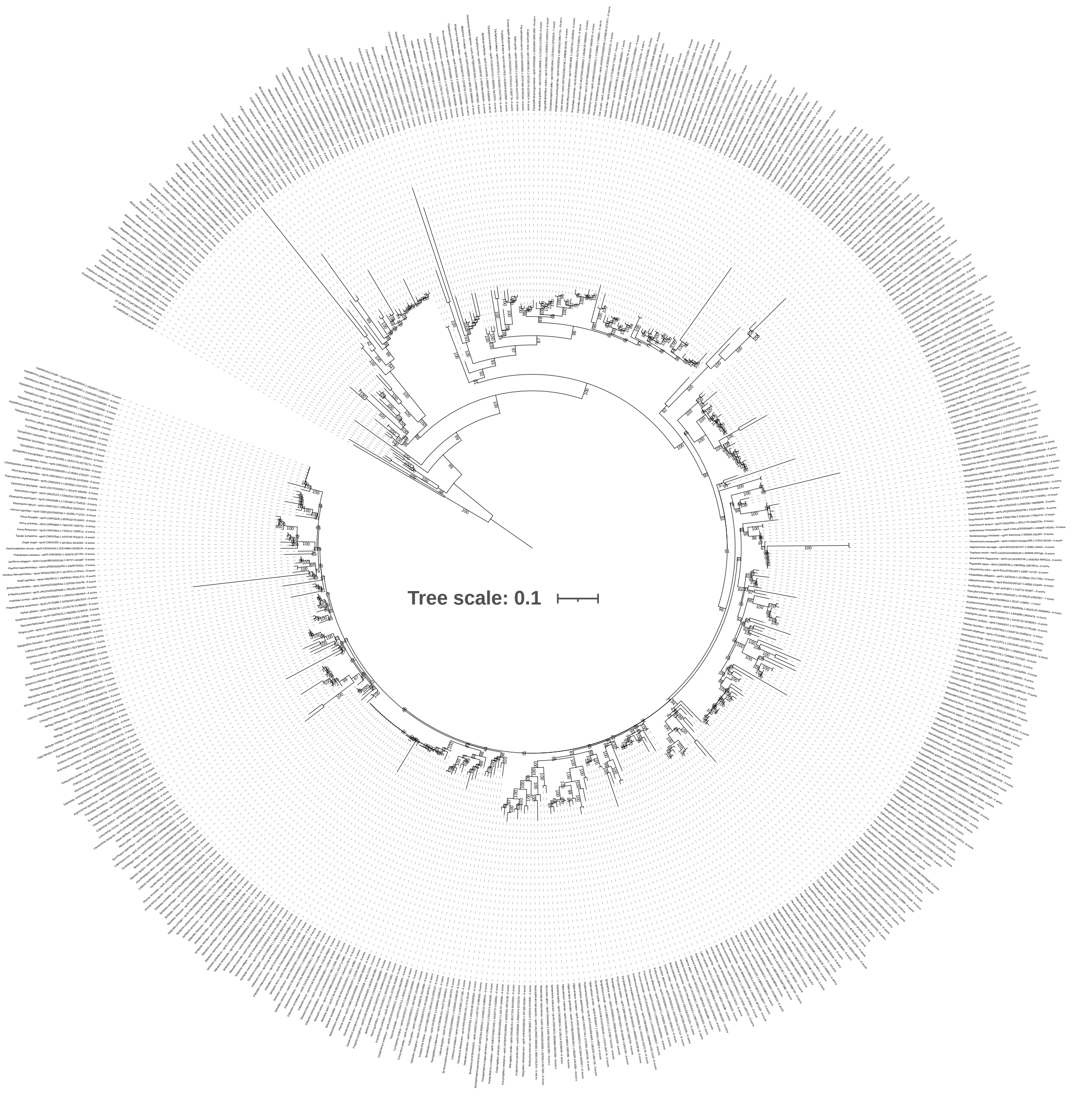


**Supplementary Figure 13.** Maximum likelihood phylogeny of *opn5* genes. The tree was computed using the optimal model, as determined by ModelFinder (JTT+R6). The robustness of the nodes was evaluated with 1,000 ultrafast bootstraps and these bootstrap values are indicated at each node. The tree was rooted using the *Callorhinchus milii* and *Rhincodon typus opn5* genes (XP_007901513.1 and XP_048453349.1 respectively).

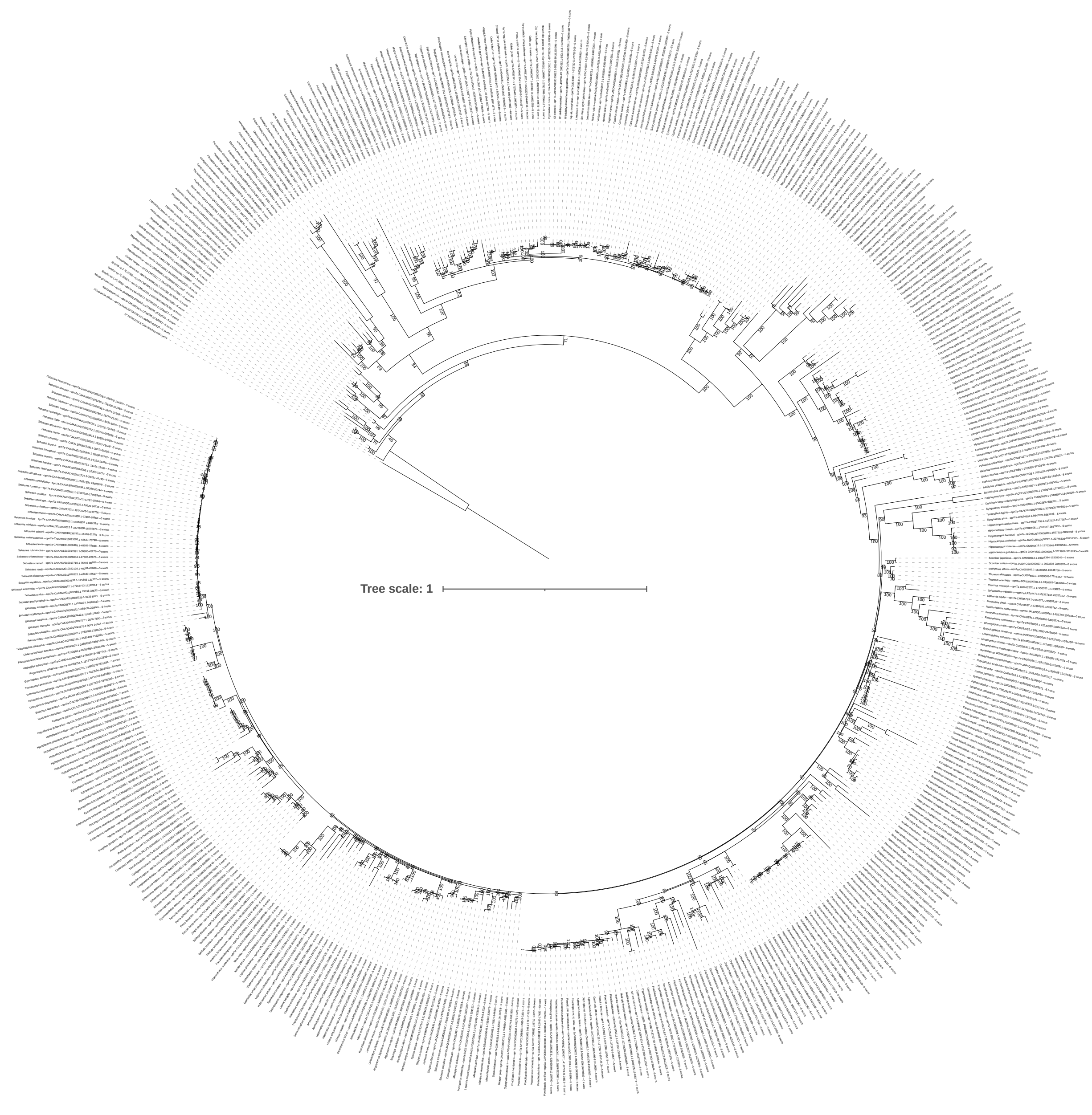


**Supplementary Figure 14.** Maximum likelihood phylogeny of *opn7a* genes. The tree was computed using the optimal model, as determined by ModelFinder (JTT+F+R6). The robustness of the nodes was evaluated with 1,000 ultrafast bootstraps and these bootstrap values are indicated at each node. The tree was rooted using the *Callorhinchus milii opn7a* gene (XP_007883754.2).

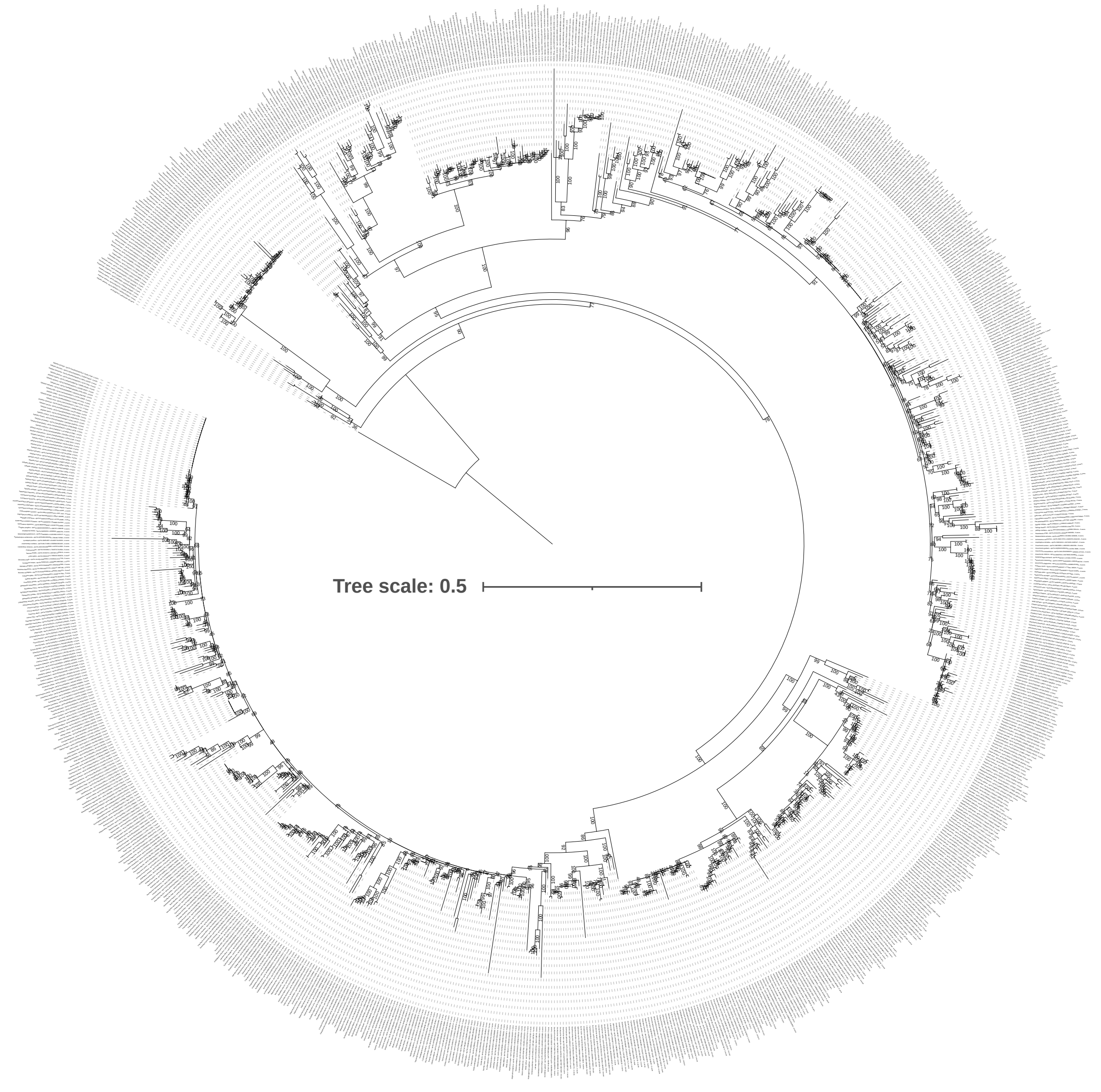


**Supplementary Figure 15.** Maximum likelihood phylogeny of *opn7b* genes. The tree was computed using the optimal model, as determined by ModelFinder (JTT+R7). The robustness of the nodes was evaluated with 1,000 ultrafast bootstraps and these bootstrap values are indicated at each node. The tree was rooted using the *Callorhinchus milii opn7a* gene (XP_007883754.2).

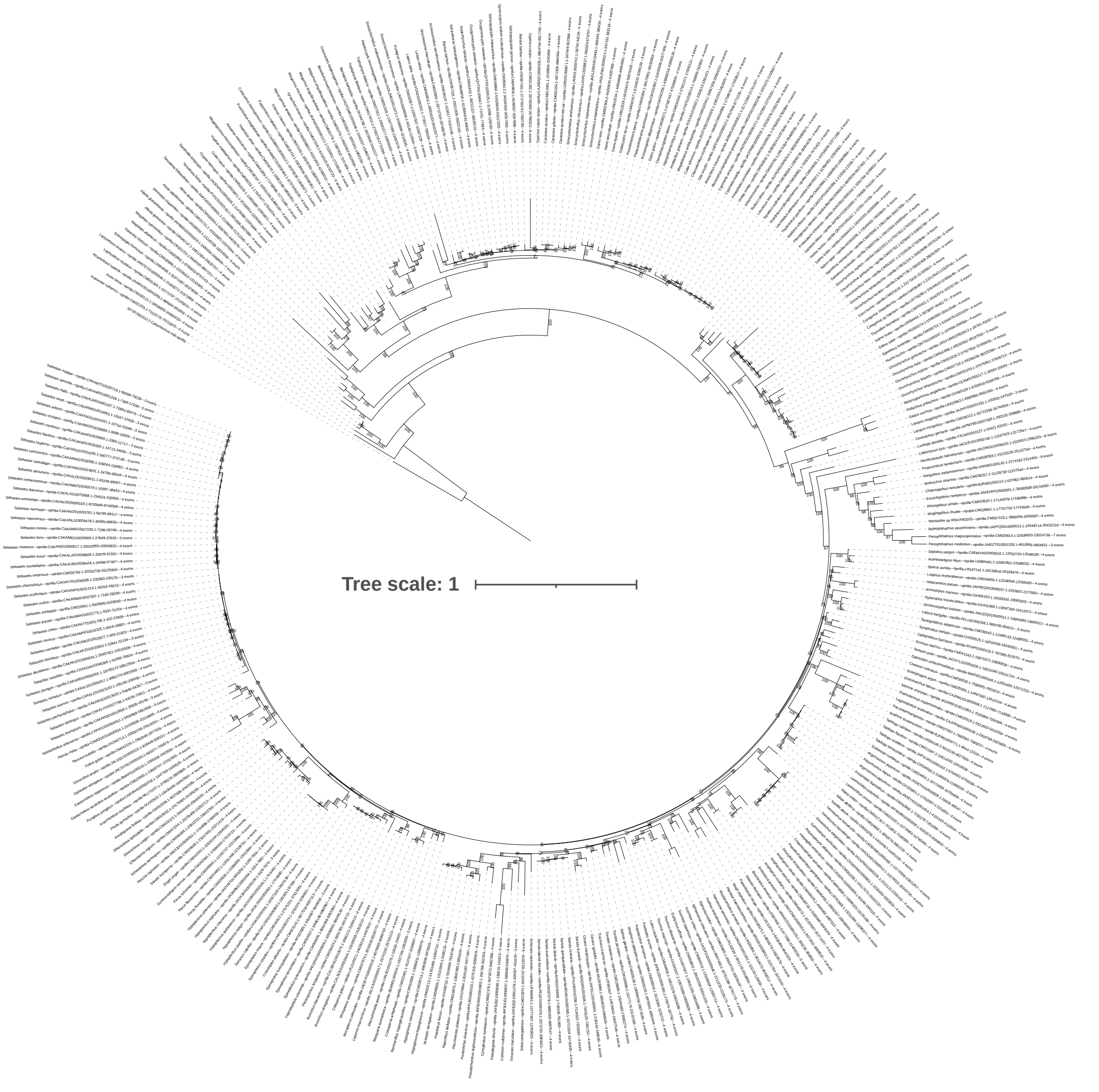


**Supplementary Figure 16.** Maximum likelihood phylogeny of *opn8a* genes. The tree was computed using the optimal model, as determined by ModelFinder (JTT+R6). The robustness of the nodes was evaluated with 1,000 ultrafast bootstraps and these bootstrap values are indicated at each node. The tree was rooted using the *Callorhinchus milii opn8b* gene (XP_007901512.2).

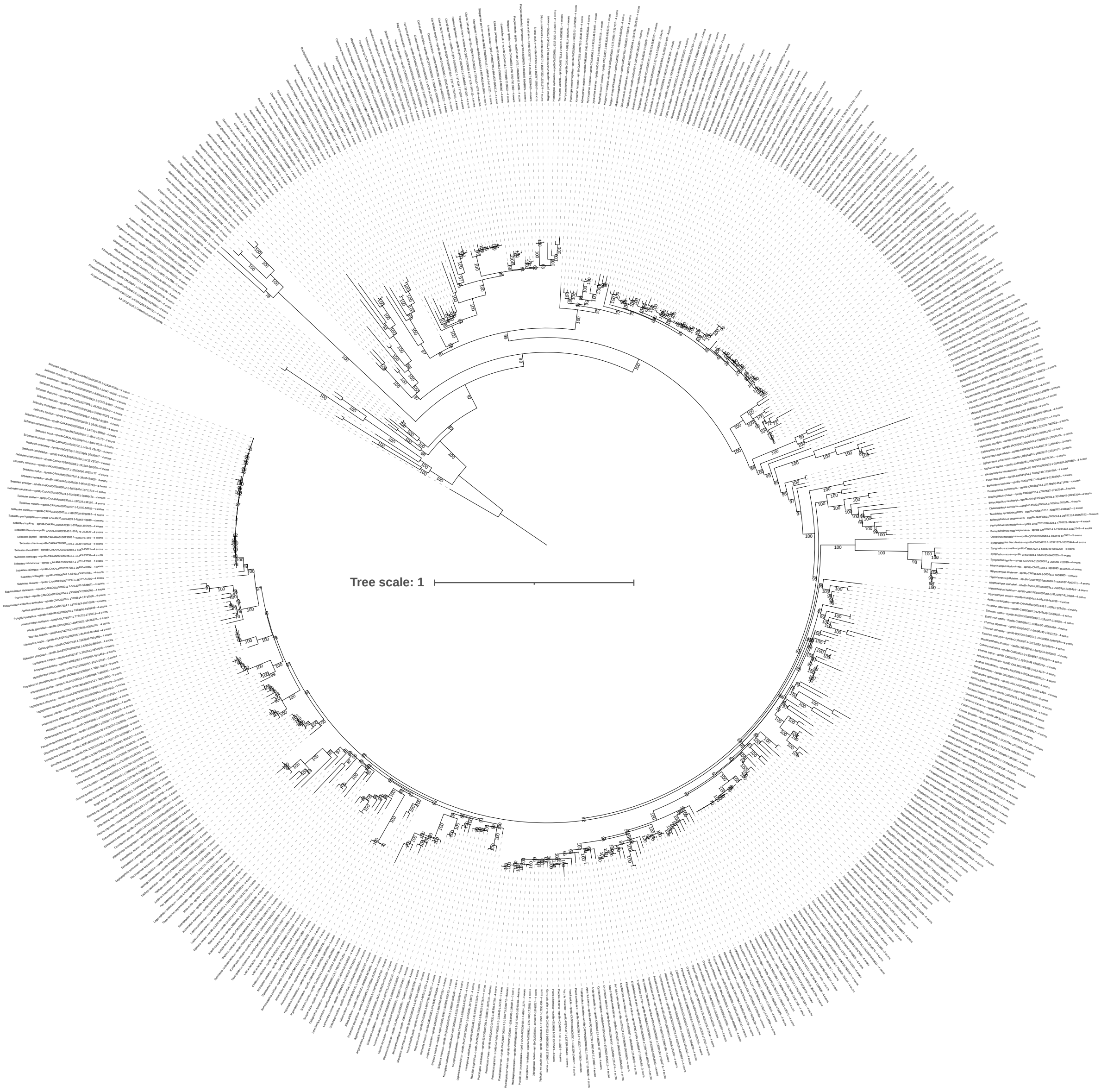


**Supplementary Figure 17.** Maximum likelihood phylogeny of *opn8b* genes. The tree was computed using the optimal model, as determined by ModelFinder (JTT+R8). The robustness of the nodes was evaluated with 1,000 ultrafast bootstraps and these bootstrap values are indicated at each node. The tree was rooted using the *Callorhinchus milii opn8b* gene (XP_007901512.2).

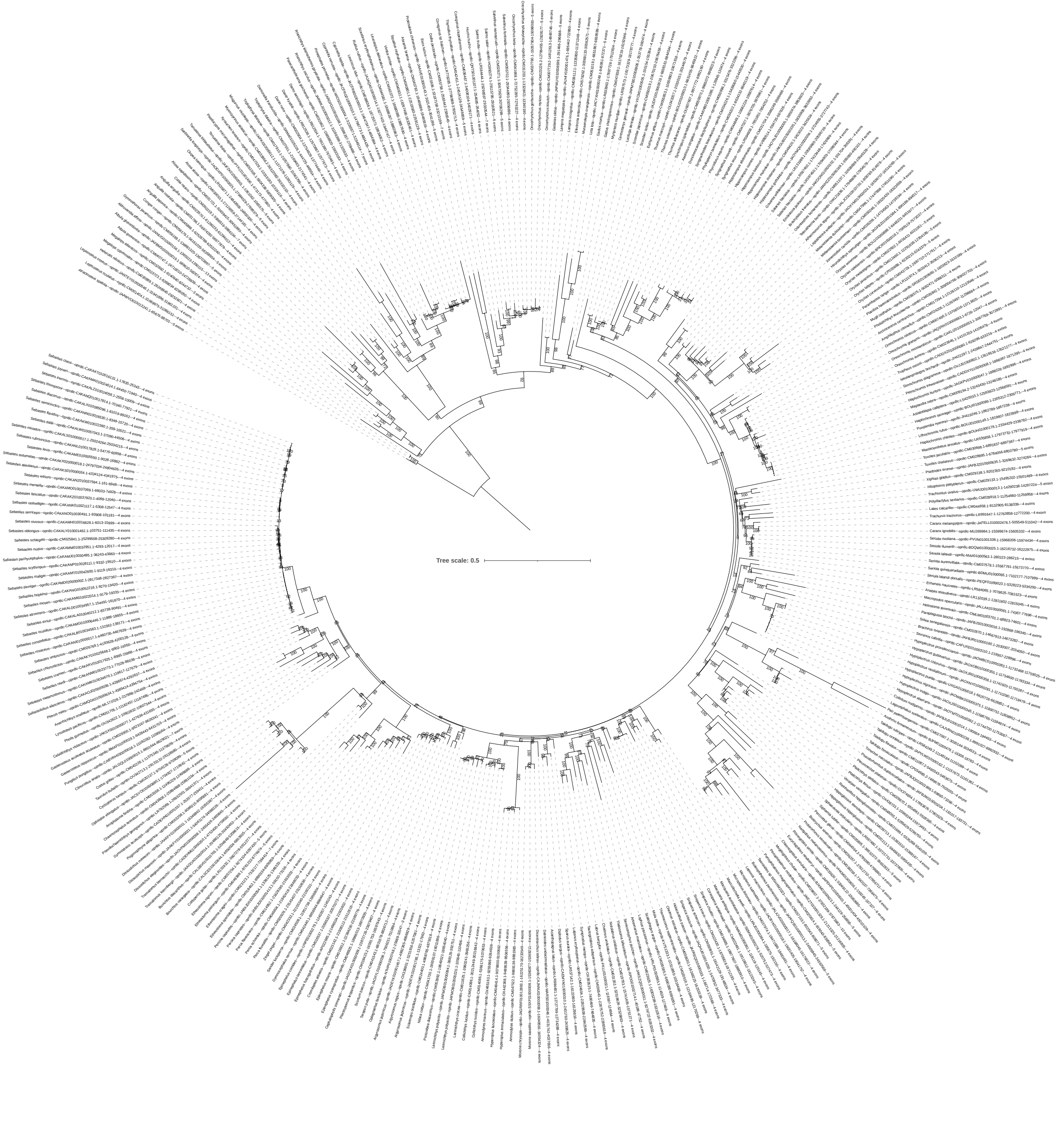


**Supplementary Figure 18.** Maximum likelihood phylogeny of *opn8c* genes. The tree was computed using the optimal model, as determined by ModelFinder (JTT+R7). The robustness of the nodes was evaluated with 1,000 ultrafast bootstraps and these bootstrap values are indicated at each node. The tree was rooted using non-teleost species opn8c genes.

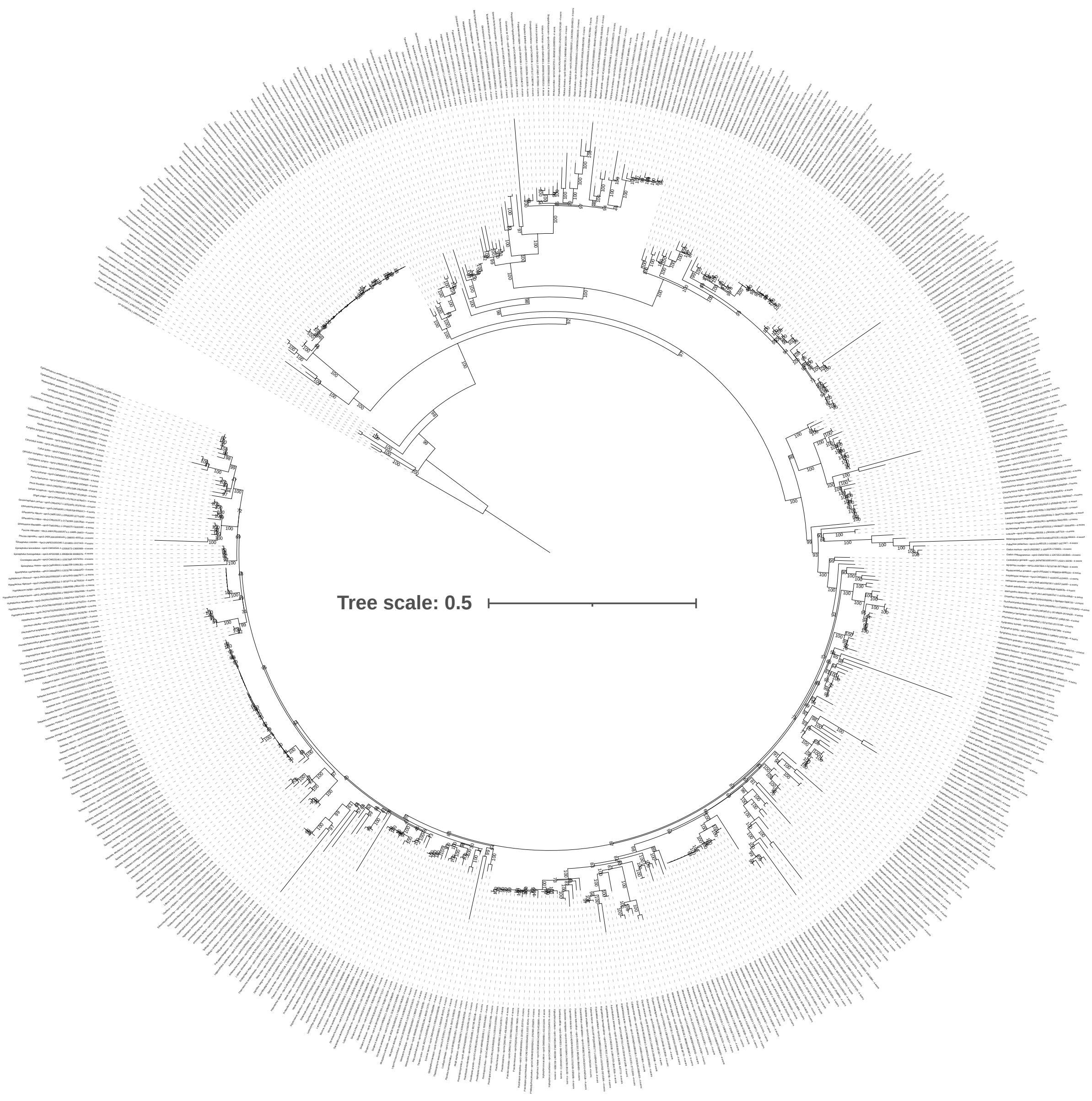


**Supplementary Figure 19.** Maximum likelihood phylogeny of *opn3* genes. The tree was computed using the optimal model, as determined by ModelFinder (JTT+R6). The robustness of the nodes was evaluated with 1,000 ultrafast bootstraps and these bootstrap values are indicated at each node. The tree was rooted using the *Callorhinchus milii opn3* gene (XP_007892106.1).

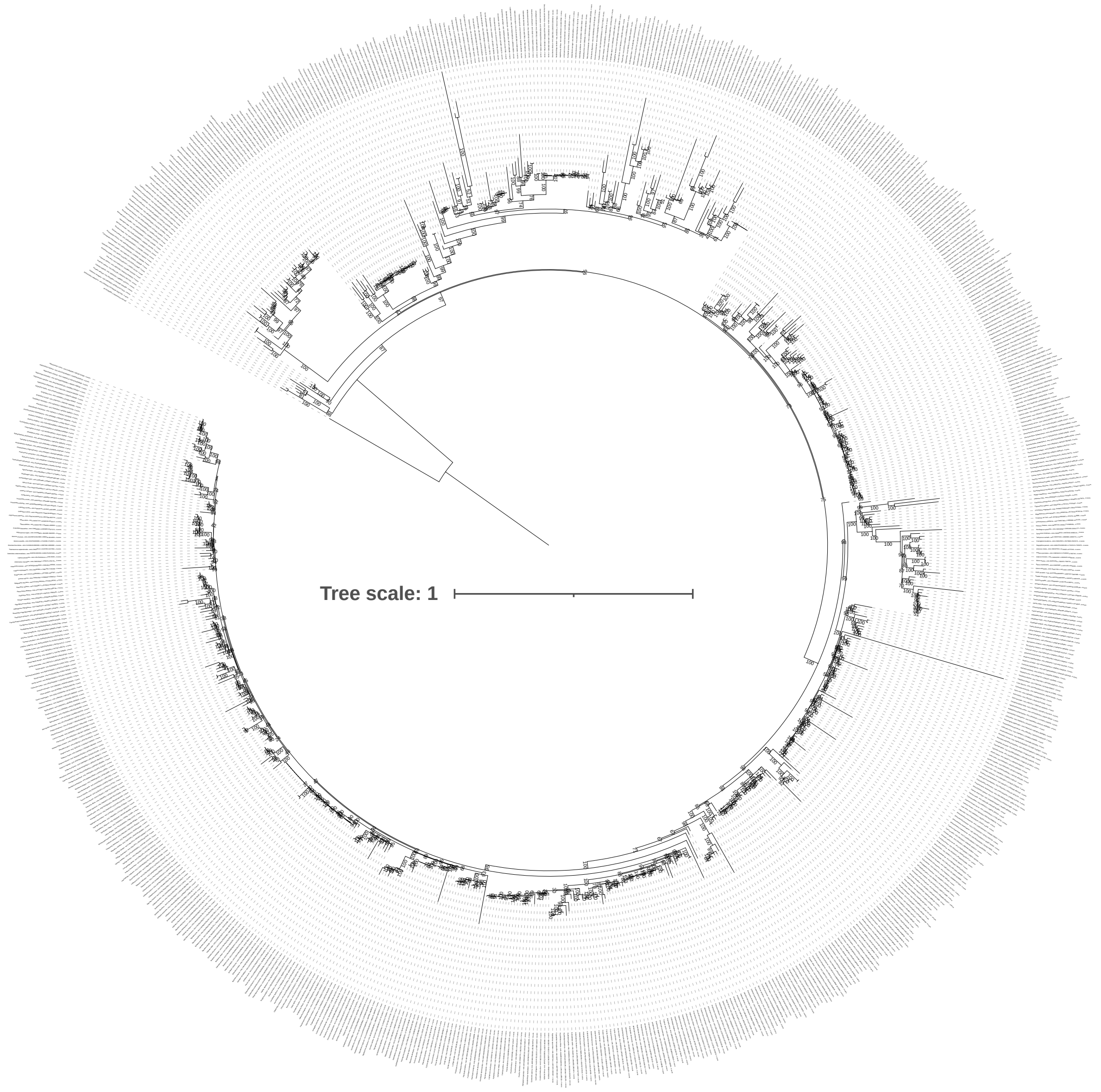


**Supplementary Figure 20.** Maximum likelihood phylogeny of *tmt1* genes. The tree was computed using the optimal model, as determined by ModelFinder (JTT+R9). The robustness of the nodes was evaluated with 1,000 ultrafast bootstraps and these bootstrap values are indicated at each node. The tree was rooted using the *Callorhinchus milii tmt2* gene (XP_007892895.2).

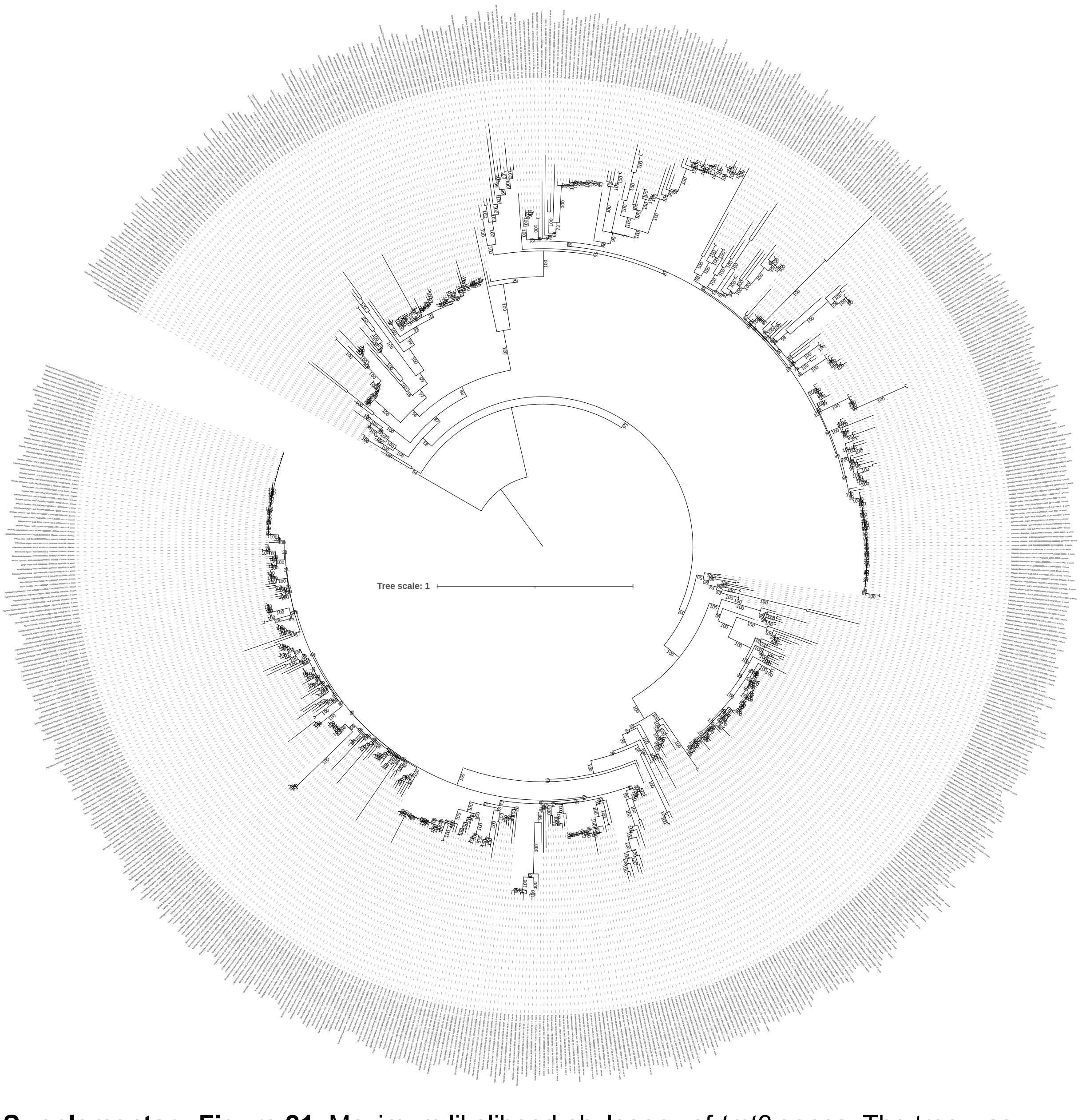


**Supplementary Figure 21.** Maximum likelihood phylogeny of *tmt2* genes. The tree was computed using the optimal model, as determined by ModelFinder (JTT+F+R8). The robustness of the nodes was evaluated with 1,000 ultrafast bootstraps and these bootstrap values are indicated at each node. The tree was rooted using the *Callorhinchus milii tmt2* gene (XP_007892895.2).

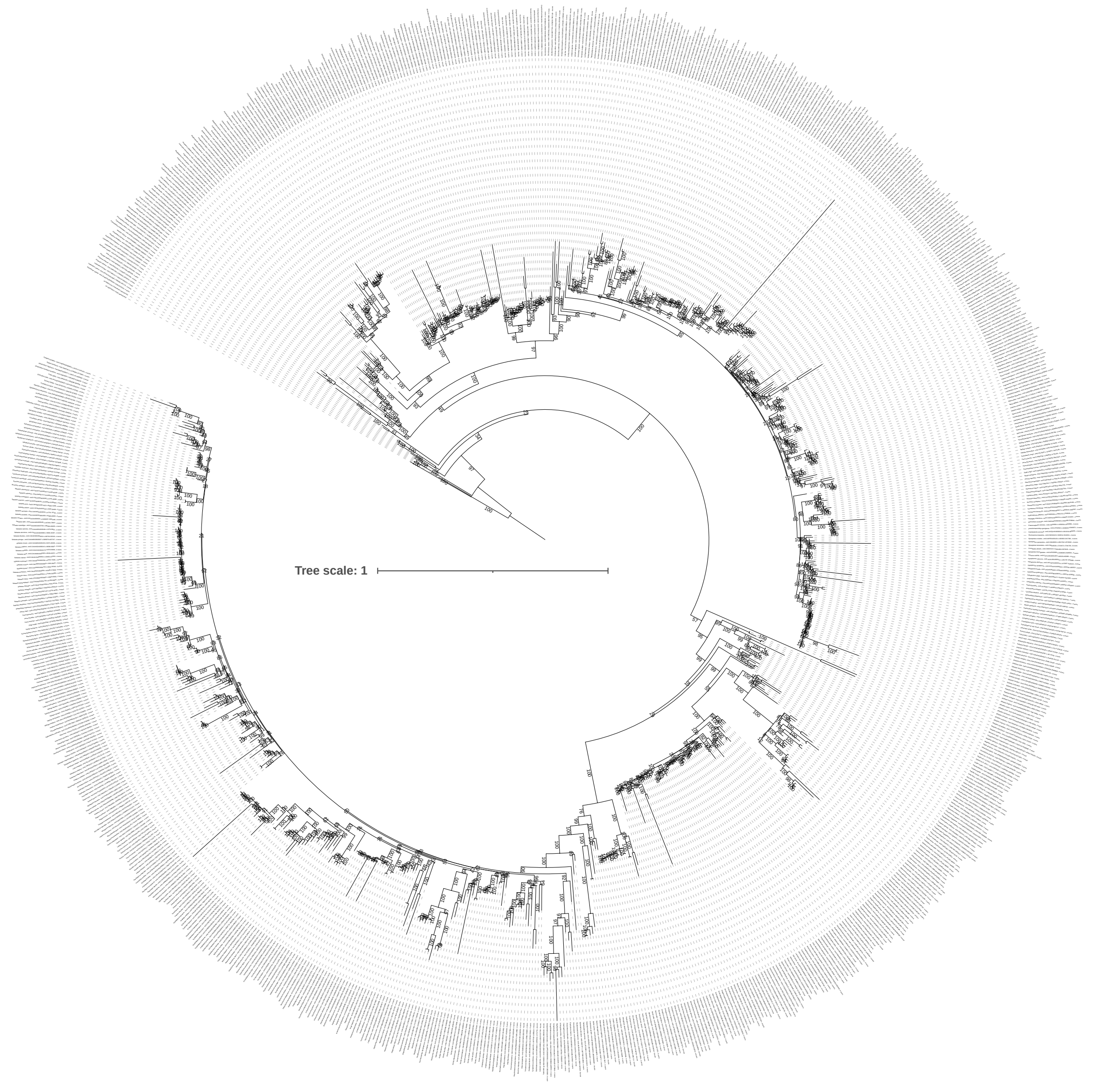


**Supplementary Figure 22.** Maximum likelihood phylogeny of *tmt3* genes. The tree was computed using the optimal model, as determined by ModelFinder (JTT+R10). The robustness of the nodes was evaluated with 1,000 ultrafast bootstraps and these bootstrap values are indicated at each node. The tree was rooted using the *Scyliorhinus canicula and Pristis pectinata tmt3* genes (XP_038672178.1 and XP_051874554.1 respectively).

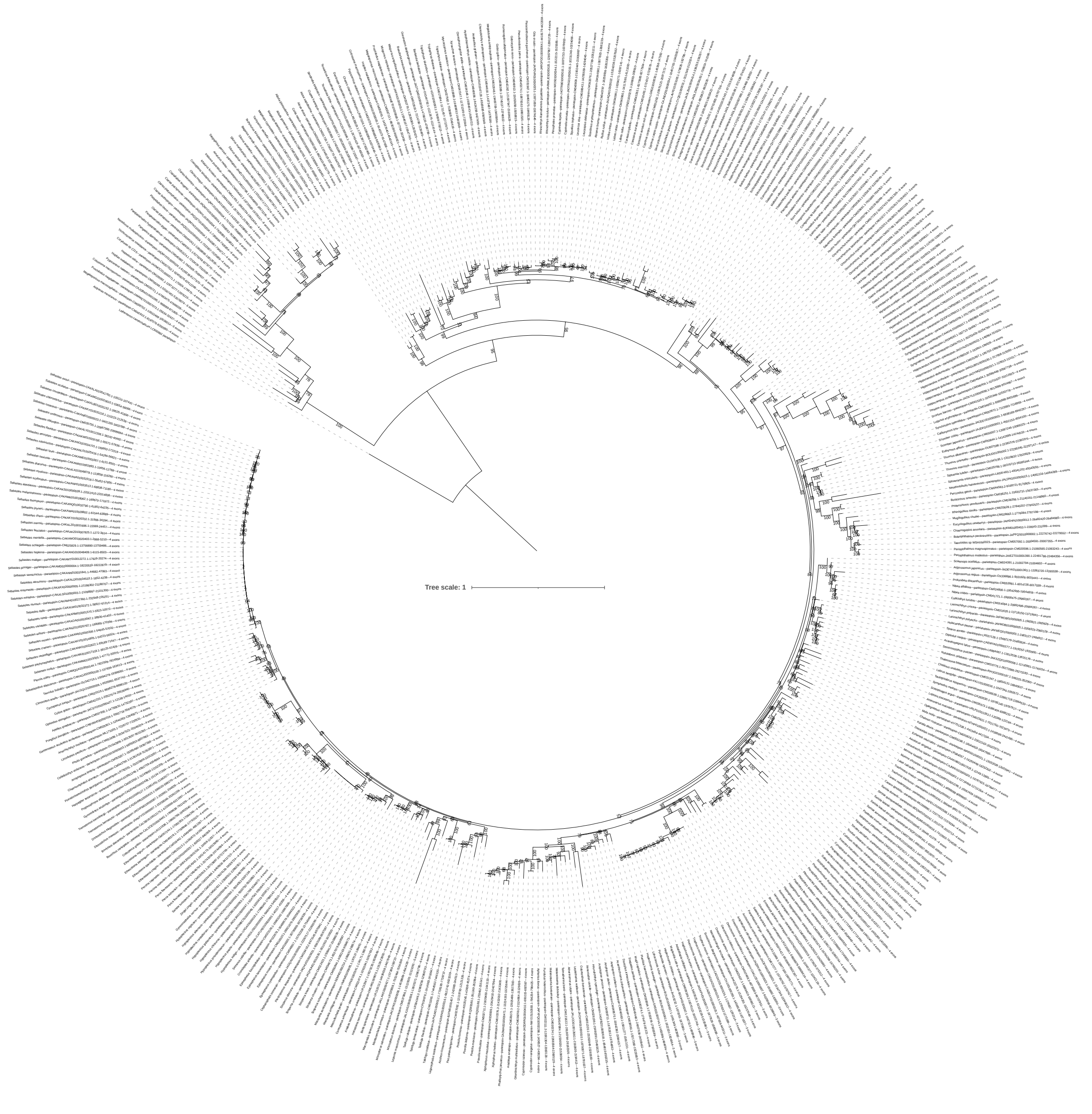


**Supplementary Figure 23.** Maximum likelihood phylogeny of *parietopsin* genes. The tree was computed using the optimal model, as determined by ModelFinder (JTT+R6). The robustness of the nodes was evaluated with 1,000 ultrafast bootstraps and these bootstrap values are indicated at each node. The tree was rooted using the *Lethenteron camtschaticum parietopsin* gene (LC600861.1).

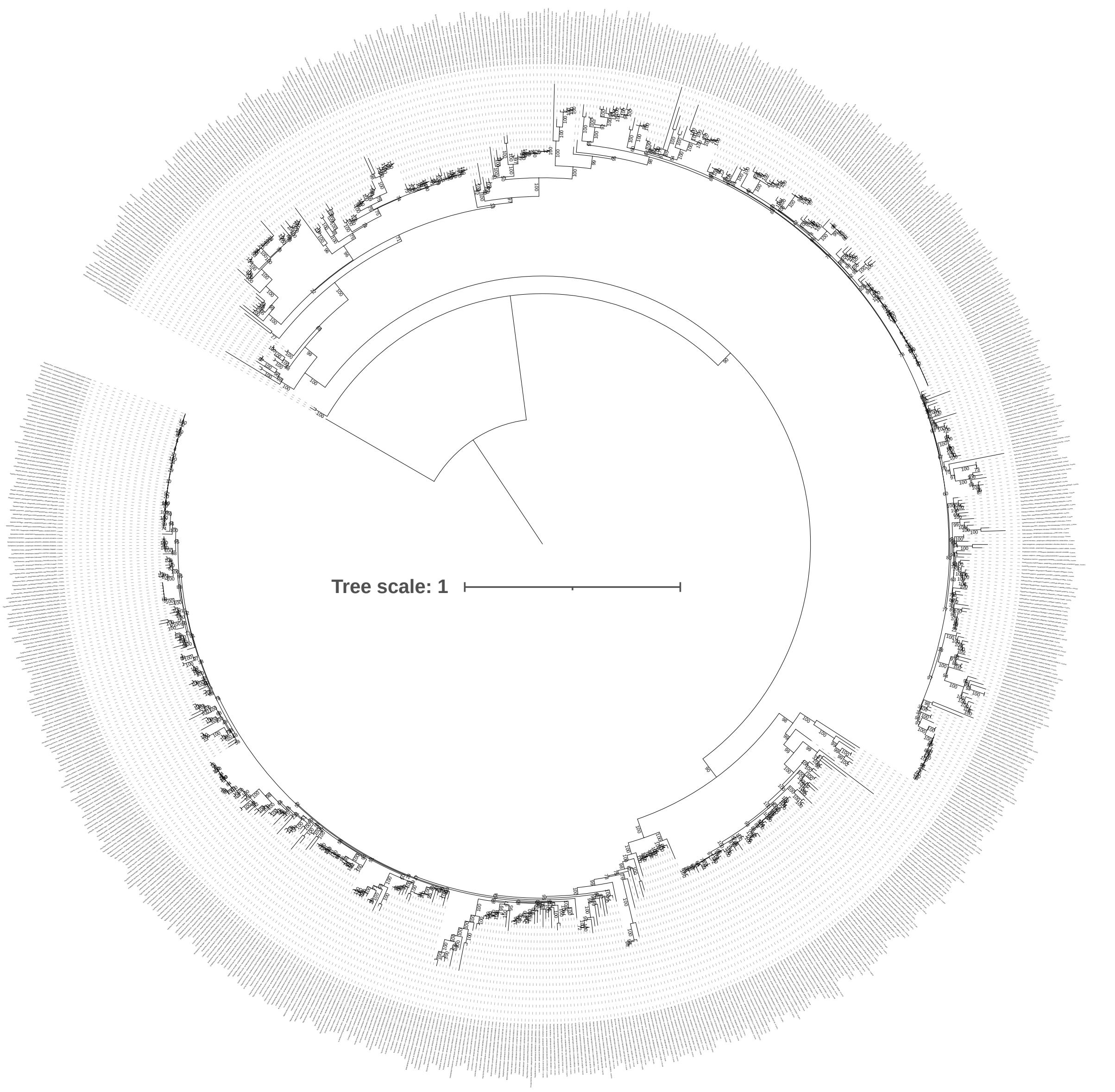


**Supplementary Figure 24.** Maximum likelihood phylogeny of *parapinopsin* genes. The tree was computed using the optimal model, as determined by ModelFinder (JTT+F+R7). The robustness of the nodes was evaluated with 1,000 ultrafast bootstraps and these bootstrap values are indicated at each node. The tree was rooted using the *Callorhinchus milii parapinopsin* gene (XP_042202474.1).

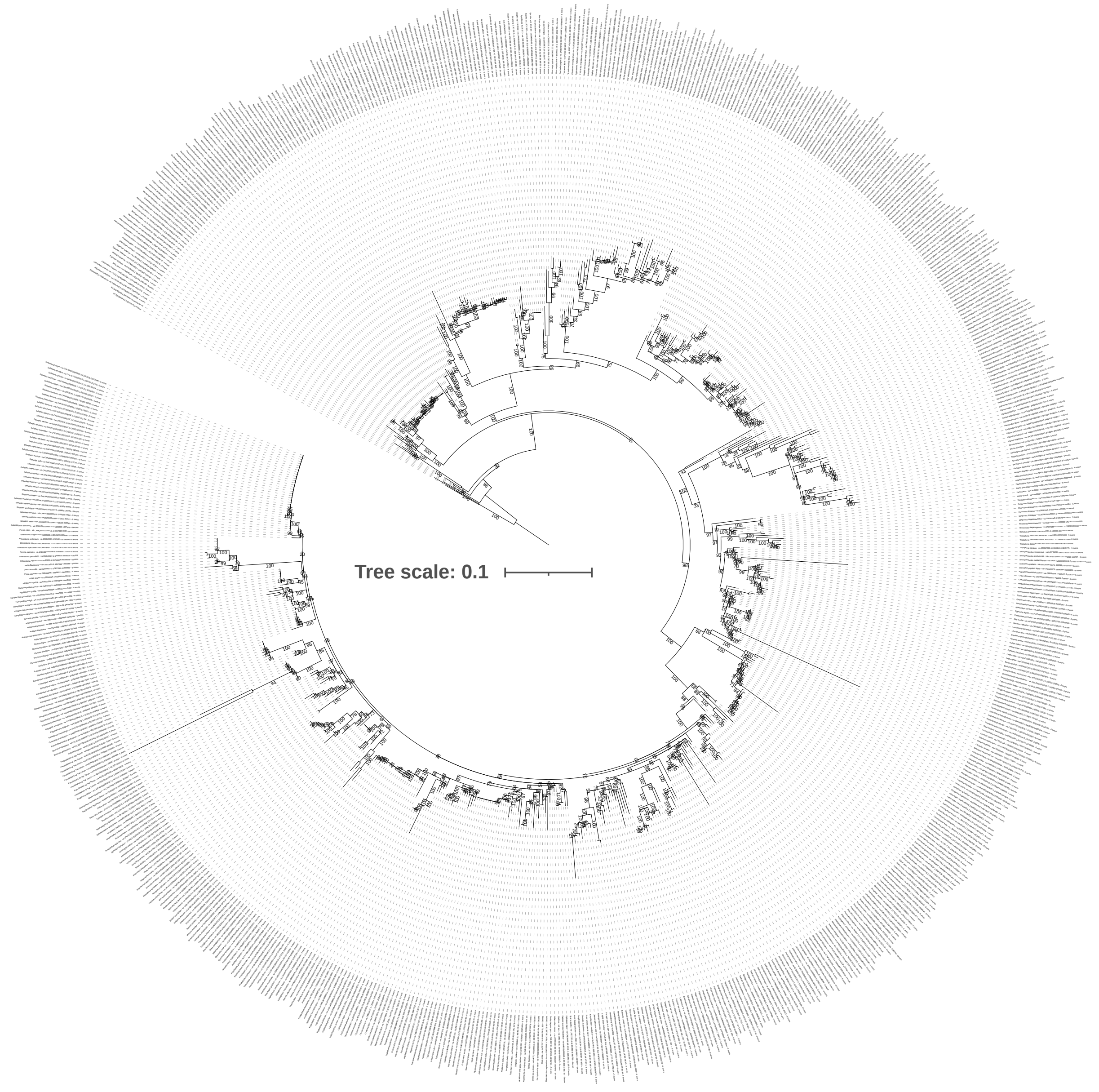


**Supplementary Figure 25.** Maximum likelihood phylogeny of *va* genes. The tree was computed using the optimal model, as determined by ModelFinder (JTT+F+R5). The robustness of the nodes was evaluated with 1,000 ultrafast bootstraps and these bootstrap values are indicated at each node. The tree was rooted using the *Callorhinchus milii and Carcharodon carcharias va* genes (XP_007894489.1 and XP_041065233.1 respectively).

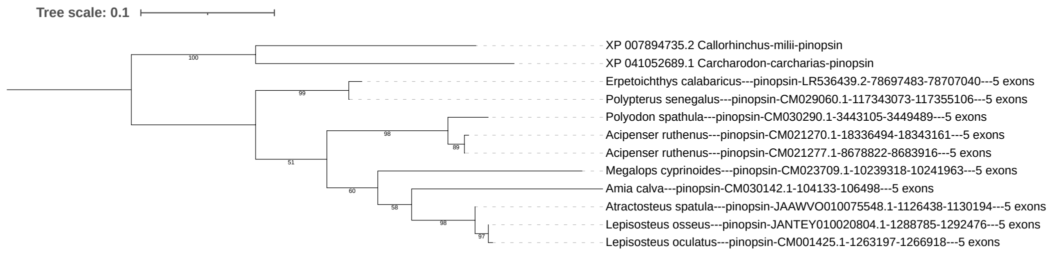


**Supplementary Figure 26.** Maximum likelihood phylogeny of *pinopsin* genes. The tree was computed using the optimal model, as determined by ModelFinder (JTT+F+R5). The robustness of the nodes was evaluated with 1,000 ultrafast bootstraps and these bootstrap values are indicated at each node. The tree was rooted using the *Callorhinchus milii and Carcharodon carcharias pinopsin* genes (XP_007894735.2 and XP_041052689.1 respectively).

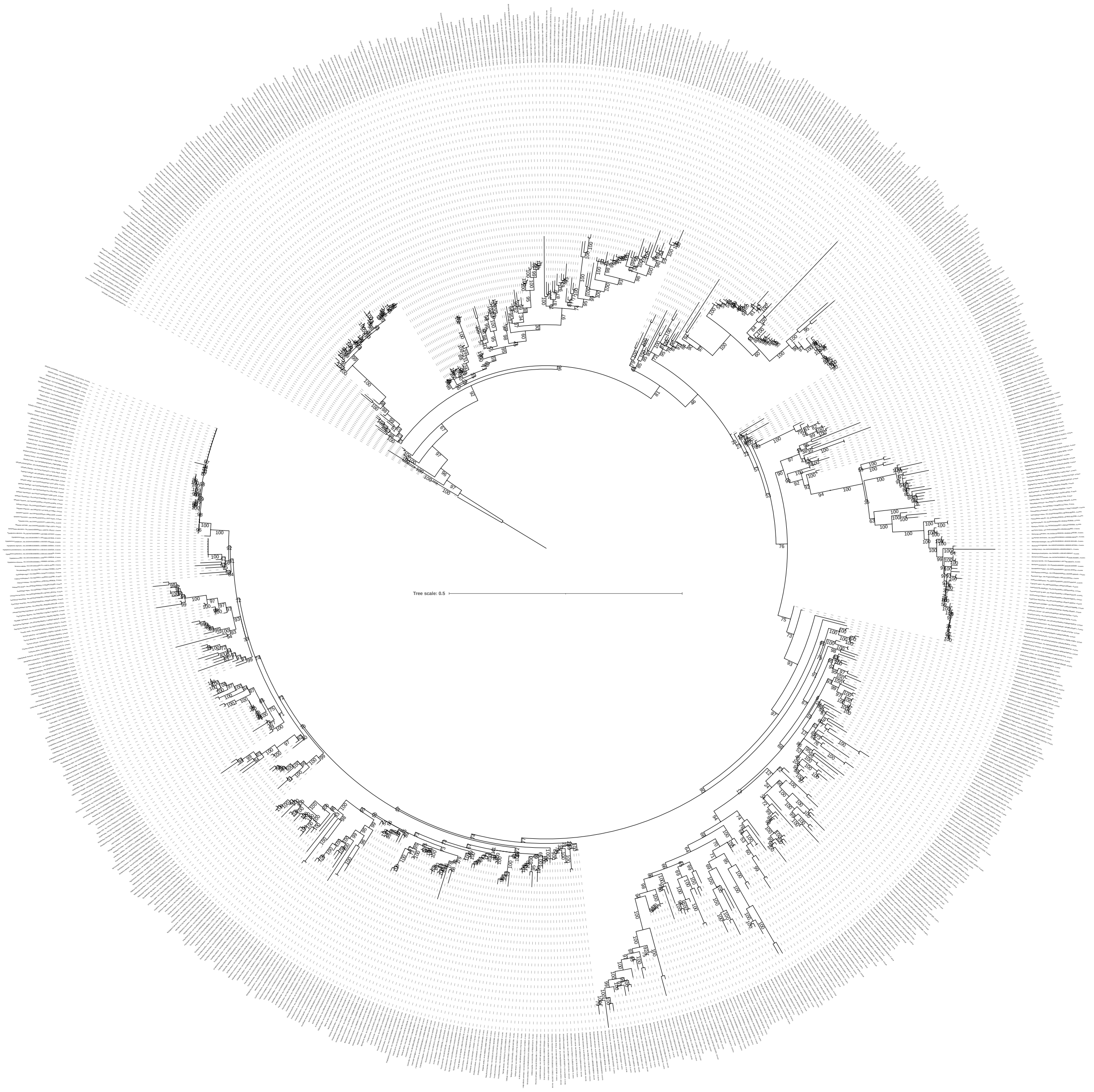


**Supplementary Figure 27.** Maximum likelihood phylogeny of *lws* genes. The tree was computed using the optimal model, as determined by ModelFinder (LG+R7). The robustness of the nodes was evaluated with 1,000 ultrafast bootstraps and these bootstrap values are indicated at each node. The tree was rooted using the *Lethenteron camtschaticum lws* gene (AB116381).

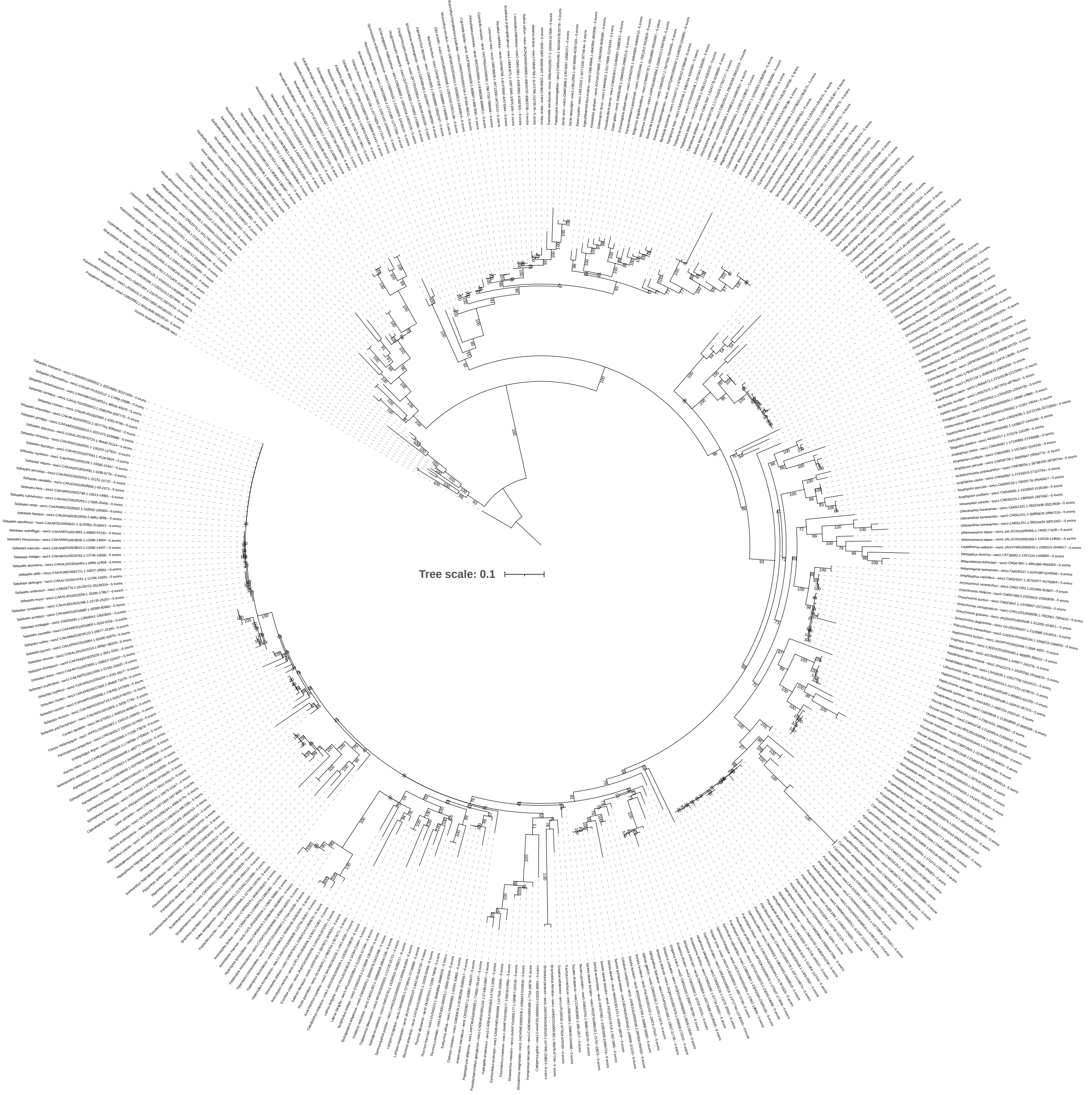

**Supplementary Figure 28.** Maximum likelihood phylogeny of *sws1* genes. The tree was computed using the optimal model, as determined by ModelFinder (LG+F+R5). The robustness of the nodes was evaluated with 1,000 ultrafast bootstraps and these bootstrap values are indicated at each node. The tree was rooted using the *Geotria australis sws1* gene (AY366495).

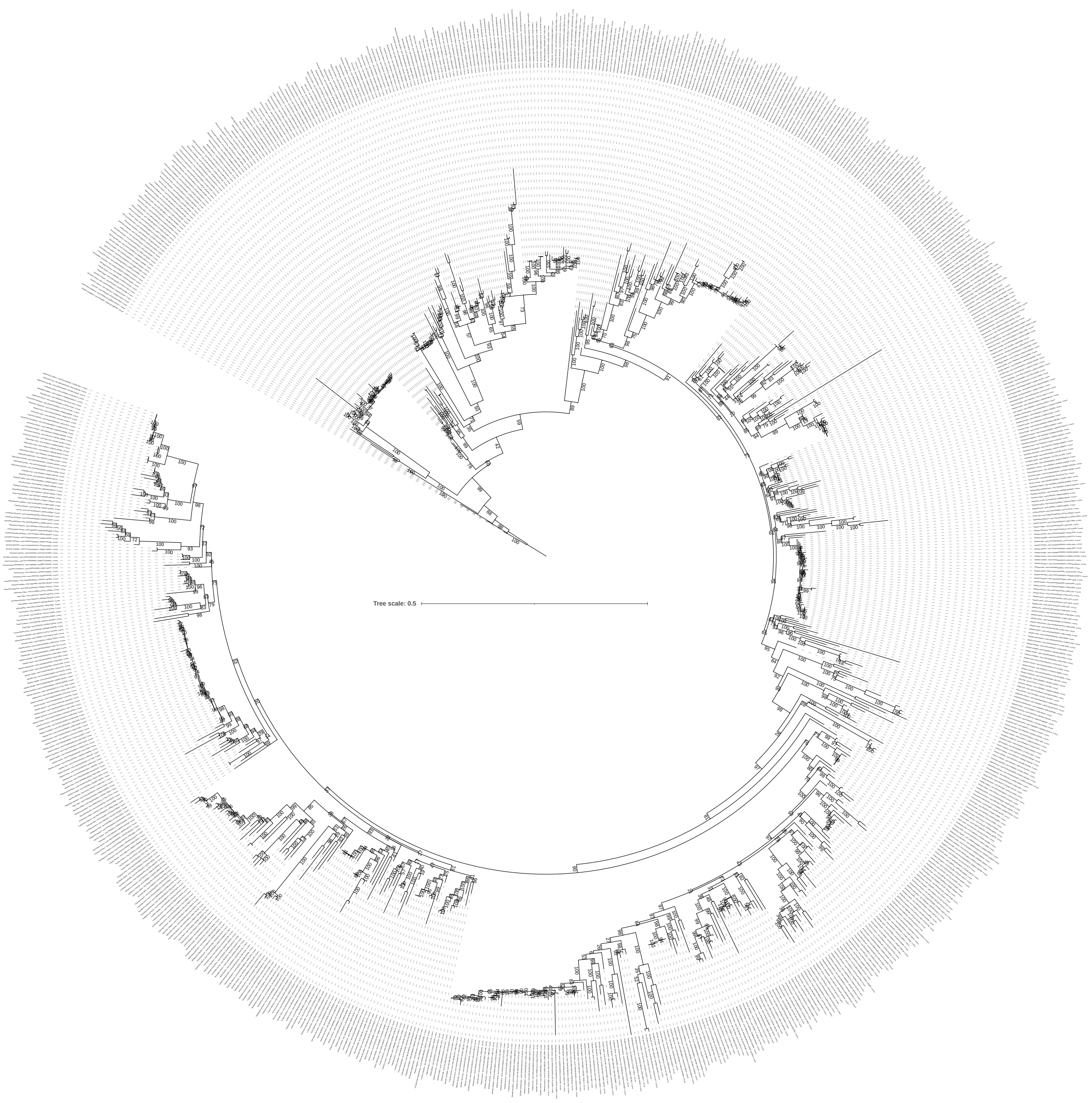


**Supplementary Figure 29.** Maximum likelihood phylogeny of *sws2* genes. The tree was computed using the optimal model, as determined by ModelFinder (mtInv+F+R8). The robustness of the nodes was evaluated with 1,000 ultrafast bootstraps and these bootstrap values are indicated at each node. The tree was rooted using the *Geotria australis sws1* gene (AY366495).

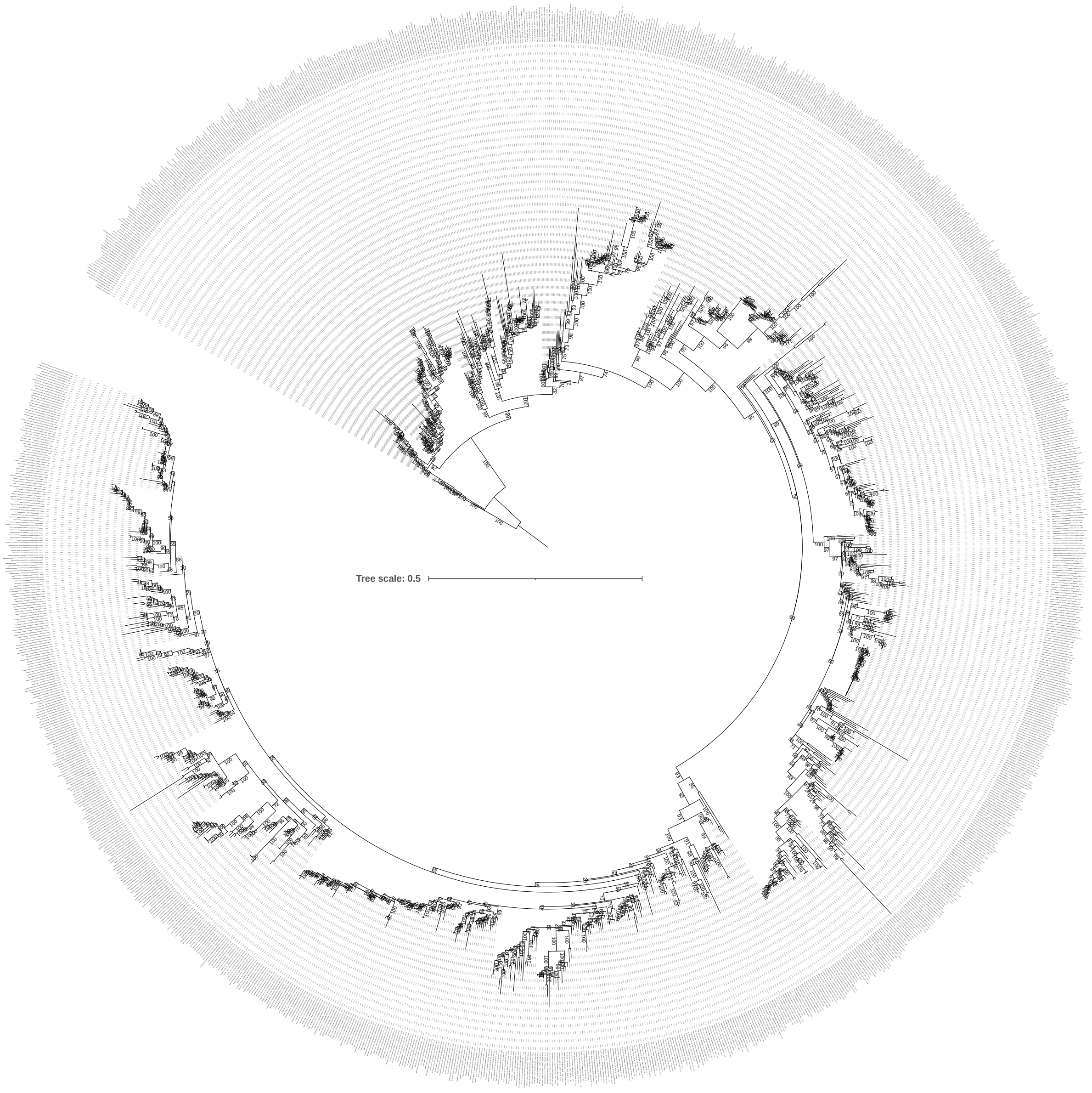


**Supplementary Figure 30.** Maximum likelihood phylogeny of *rh2* genes.The tree was computed using the optimal model, as determined by ModelFinder (LG+F+R9). The robustness of the nodes was evaluated with 1,000 ultrafast bootstraps and these bootstrap values are indicated at each node. The tree was rooted using the *Pristis pectinata and Hypanus sabinus rh2* genes (XP_051890468.1 and XP_059806431.1 respectively).

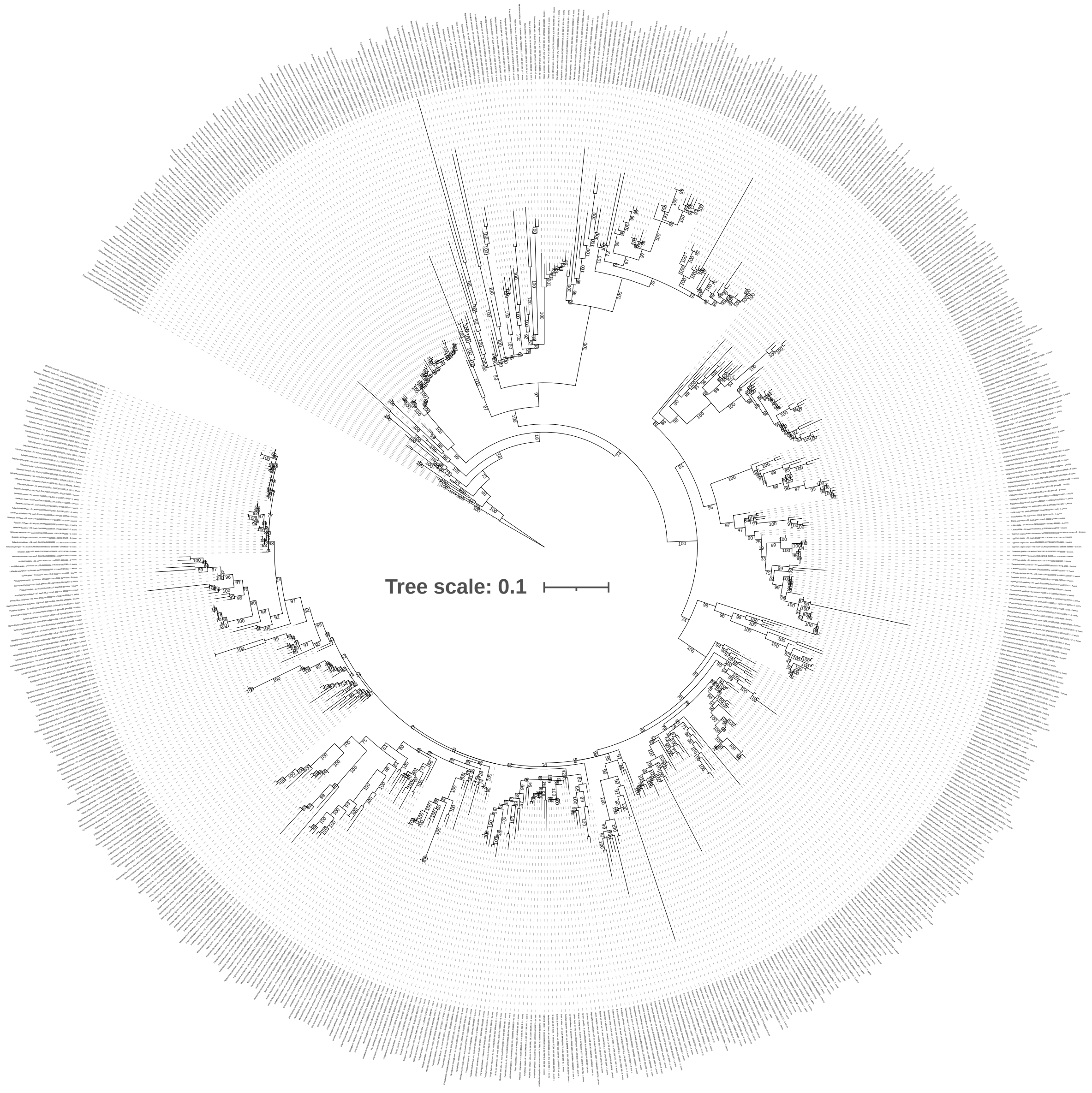


**Supplementary Figure 31.** Maximum likelihood phylogeny of *rh1* genes. The tree was computed using the optimal model, as determined by ModelFinder (mtZOA+R7). The robustness of the nodes was evaluated with 1,000 ultrafast bootstraps and these bootstrap values are indicated at each node. The tree was rooted using the *Rhincodon typus rh1* gene (A0A7T8R2L6).

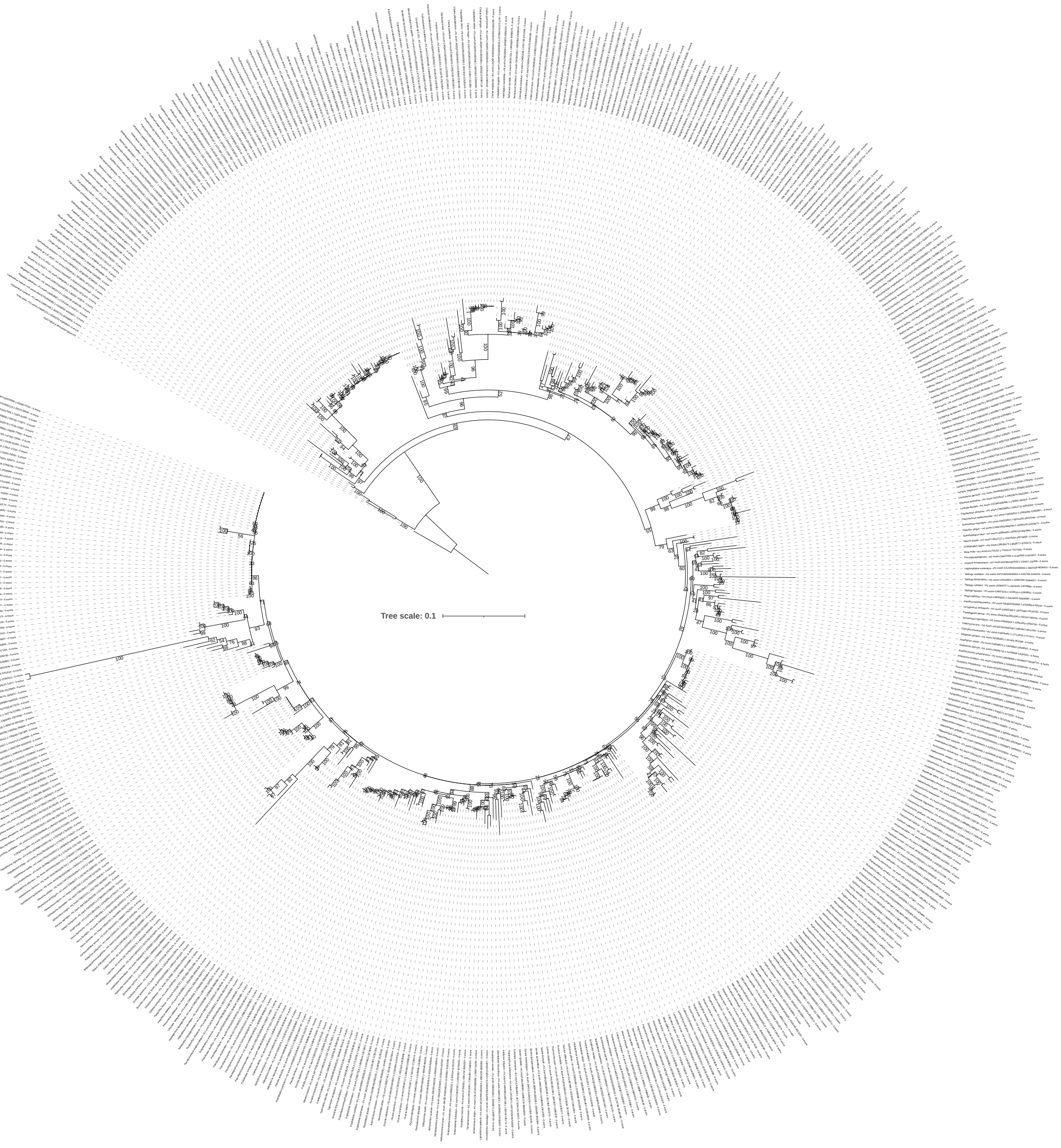


**Supplementary Figure 32.** Maximum likelihood phylogeny of *exorh* genes. The tree was computed using the optimal model, as determined by ModelFinder (JTT+R5). The robustness of the nodes was evaluated with 1,000 ultrafast bootstraps and these bootstrap values are indicated at each node. The tree was rooted using the *Rhincodon typus rh1* gene (A0A7T8R2L6).

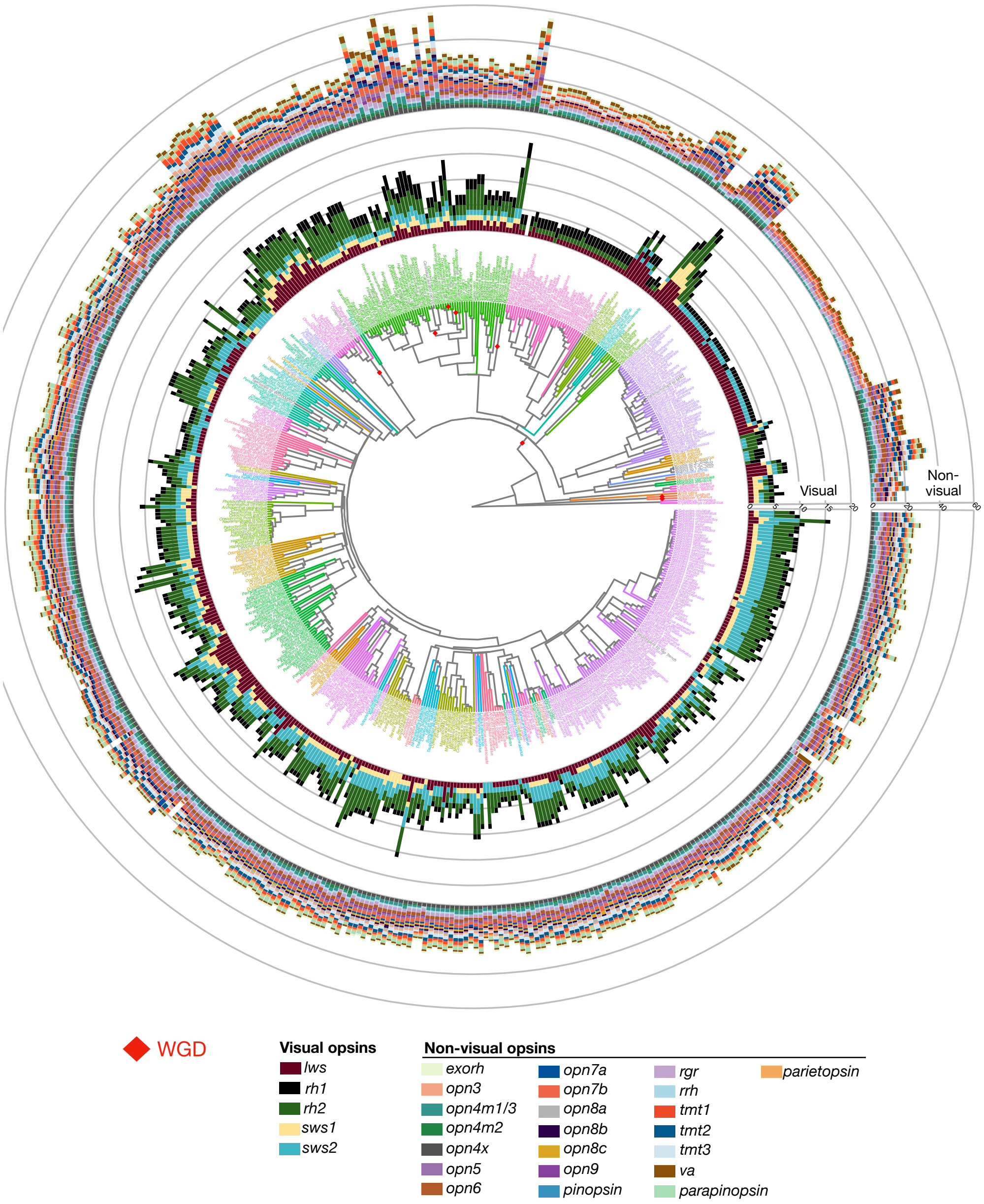


**Supplementary Figure 33.** Phylogeny of 535 ray-finned fish species computed with ASTRAL, using 2,000 BUSCO genes, and dated using the least-square dating method. Branches are colored according to the order assigned on the NCBI taxonomy database. The numbers of complete opsins in each subfamily are shown as barplots, with the inner barplot layer representing visual opsins and the outer barplot layer representing non-visual opsins. Red diamonds represent whole-genome duplication events.

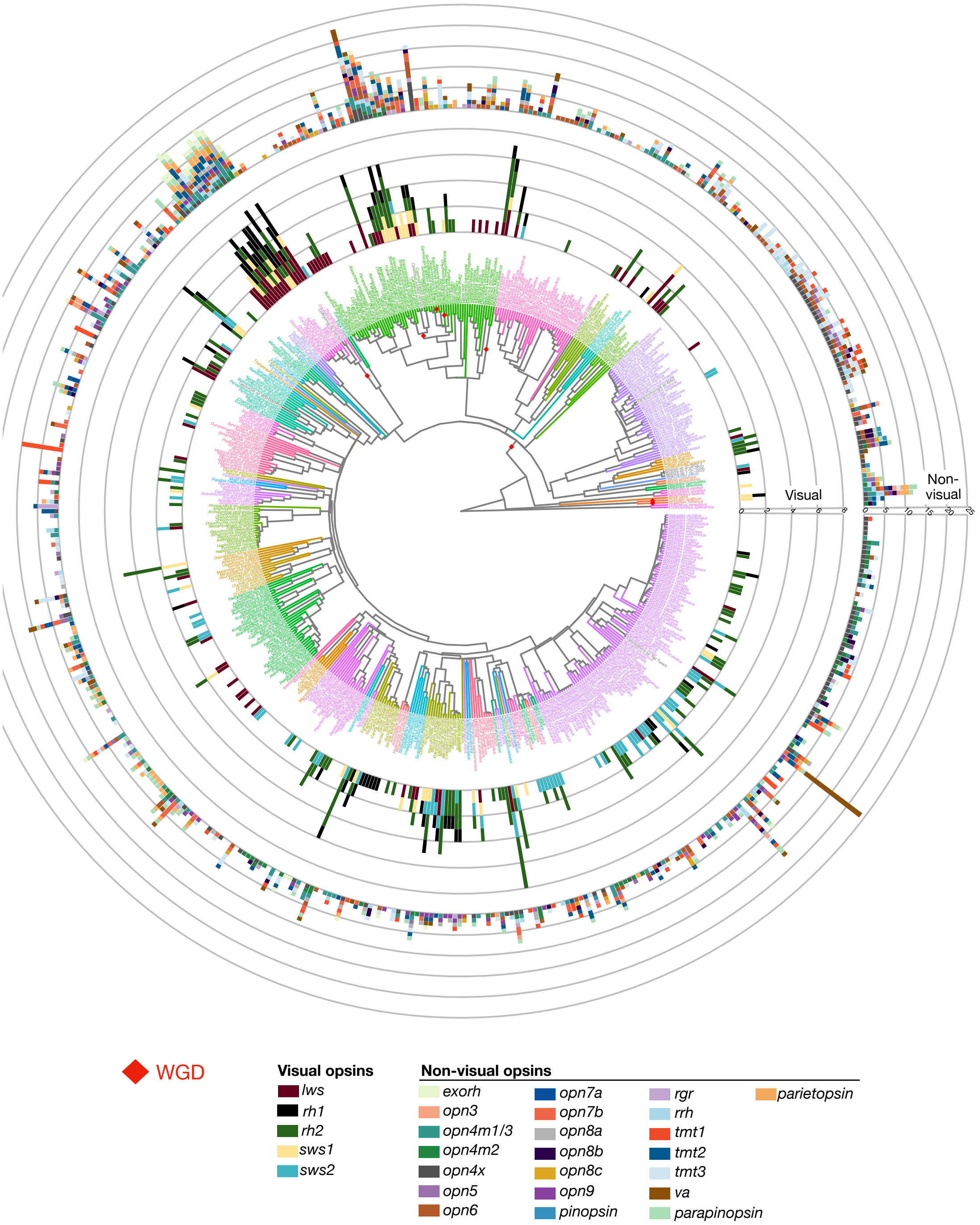


**Supplementary Figure 34.** Phylogeny of 535 ray-finned fish species computed with ASTRAL, using 2,000 BUSCO genes, and dated using the least-square dating method. Branches are colored according to the order assigned on the NCBI taxonomy database. The numbers of opsin pseudogenes in each subfamily are shown as barplots, with the inner barplot layer representing visual opsins and the outer barplot layer representing non-visual opsins. Red diamonds represent whole-genome duplication events.

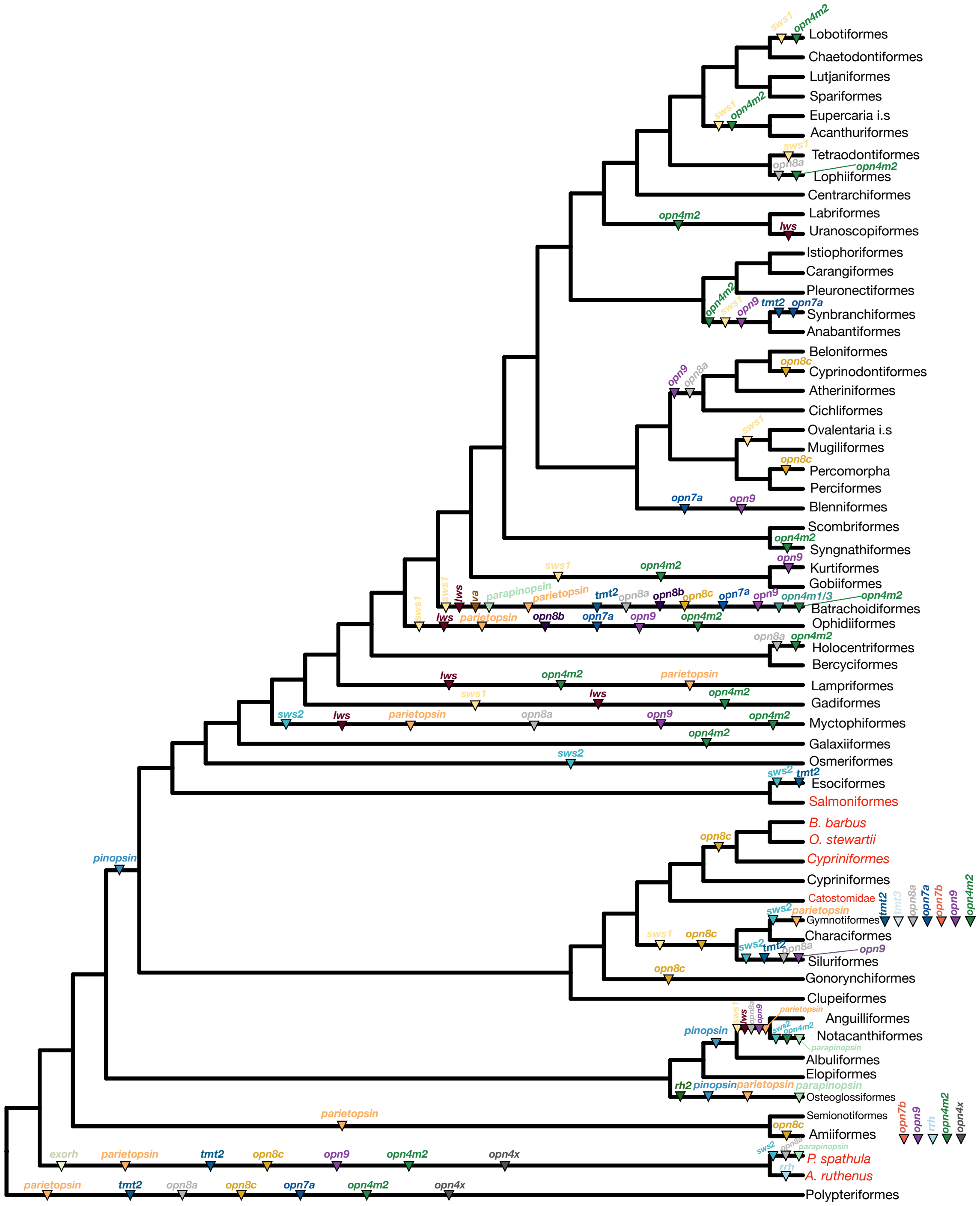


**Supplementary Figure 35.** Phylogeny of ray-finned fishes inferred using ASTRAL and dated using least square dating method. Order- or species-specific whole-genome duplications are indicated with red tip labels. Five branches correspond to Cypriniformes: one black branch for diploid species, one orange branch for tetraploid species of the genus Carassius, Cyprinus and Sinocyclocheilus as well as three orange branches for Catostomidae, O. stewartia and B. barbus that underwent linages-specific WGD. Triangles represent the complete loss of one opsin subfamily and are colored accordingly. The number of opsins per species and the mean number of opsins per order can be found in Supplementary data 1.

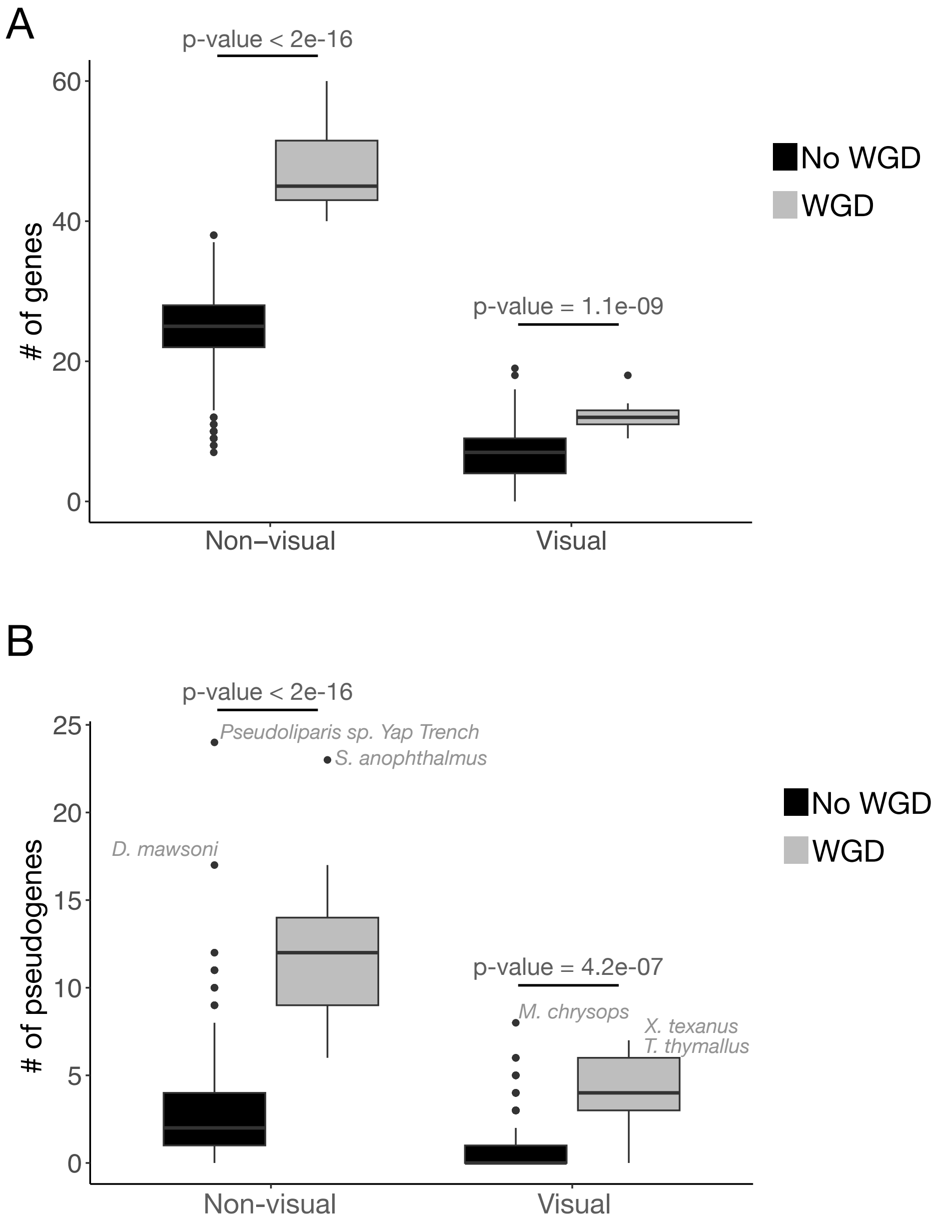


**Supplementary Figure 36.** Comparison of the number of (A) complete genes or (B) pseudogenes between diploid (black, N=500) and tetraploid (gray, N=27) teleosts. pGLS p-values for each comparison are indicated. The three species with the highest number of visual opsin or non-visual opsin pseudogenes are indicated.

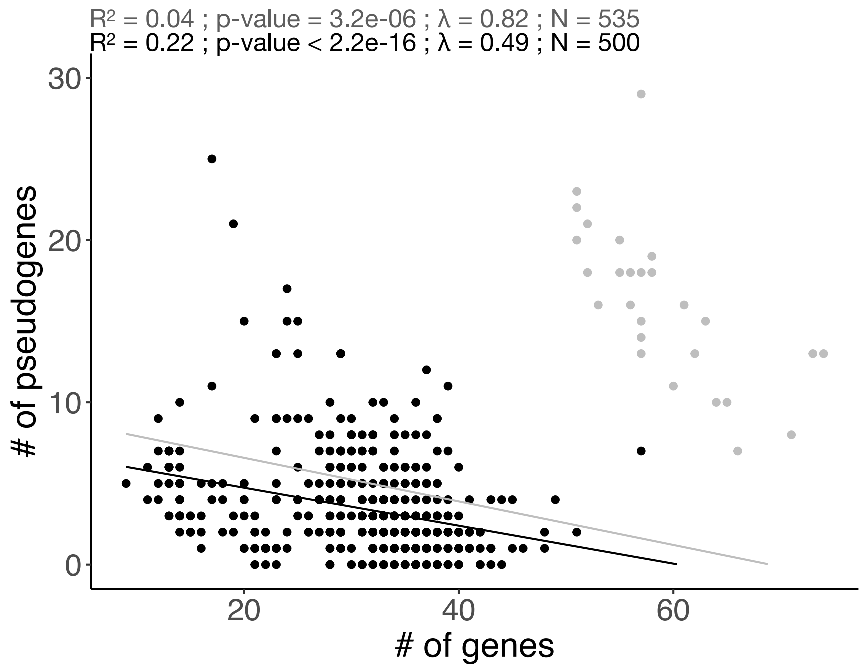


**Supplementary Figure 37.** pGLS between the number of pseudogenes and the number of genes per species. Gray points represent teleost species with recent whole-genome duplications. pGLS $R^2$, p-value, λ and regression line considering all ray-finned fishes (gray) or only diploid teleosts (black) are indicated. N represents the number of species.

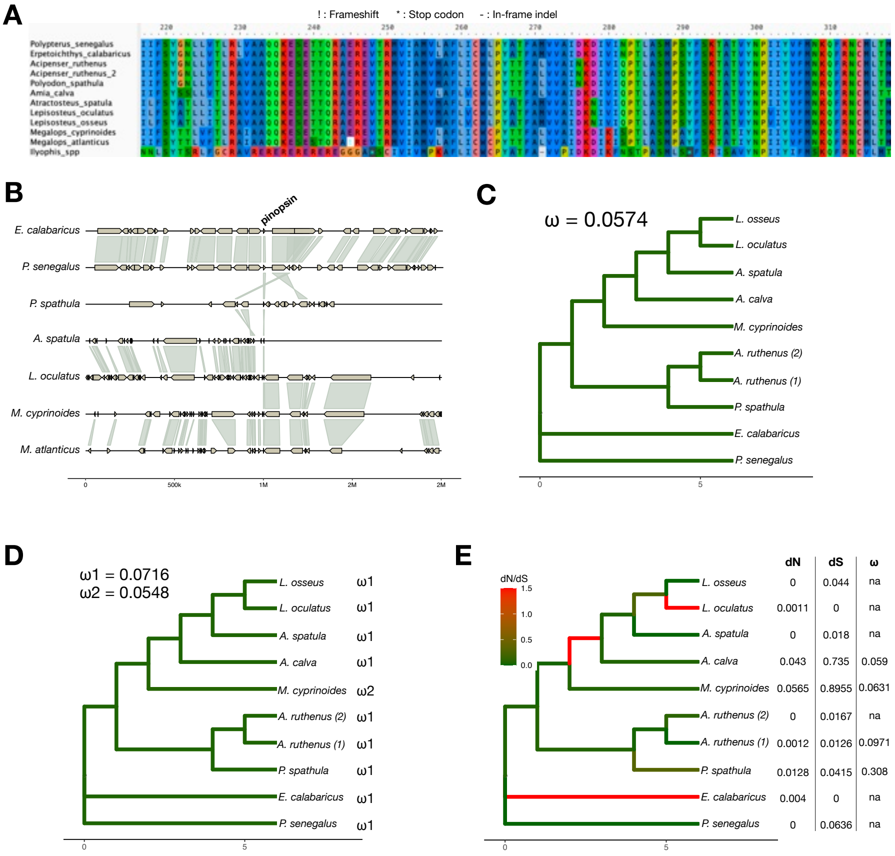


**F**

| Model | np | lnL |
|---|---|---|
| One Ratio | 19 | -4009.558 |
| Two Ratio | 20 | -4009.14 |
| Free Ratio | 35 | -3995.661 |

**G**

| | LRT | p-value |
|---|---|---|
| Free Ratio vs One Ratio | 27.79476 | 0.03345086 |
| Free Ratio vs Two Ratio | 26.95812 | 0.02907946 |
| Two Ratio vs One Ratio | 0.836644 | 0.3603584 |

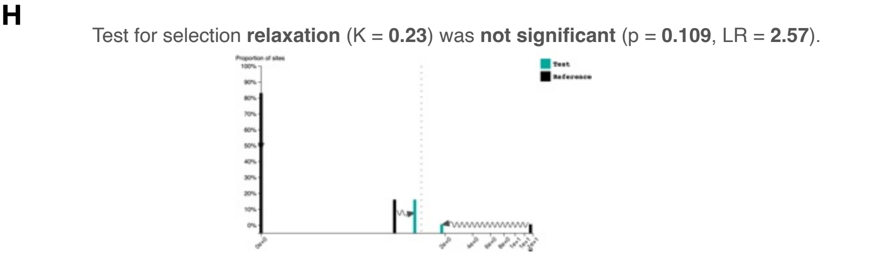


**Supplementary Figure 38.** *pinopsin* evolution. (A) Alignment of *pinopsin* extracted from non-teleost species, with *M. cyprinoides* complete *pinopsin* as well as with *M. atlanticus* and *Ilyophis* spp. *pinopsin* pseudogenes. (B) Synteny of the region surrounding *pinopsin* in non-teleost species and *Megalops* species. Genes are represented by gray arrows and orthologous genes are linked by green shades. The gene corresponding to *pinopsin* is indicated. (C) dN/dS per branch under a one-ratio model. (D) dN/dS per branch under a two-ratio model. (E) dN, dS and dN/dS per branch under a free-ratio model. The dN/dS of some terminal branches could not be computed as dN or dS was equal to 0. (F) Number of parameters (np) and likelihood score (lnL) per model. (G) Likelihood ratio test (LRT) between models and p-values computed based on the χ2 distribution. (H) Result of RELAX when assigning *M. cyprinoides* as test branch and all other (non-teleost) branches as foreground.

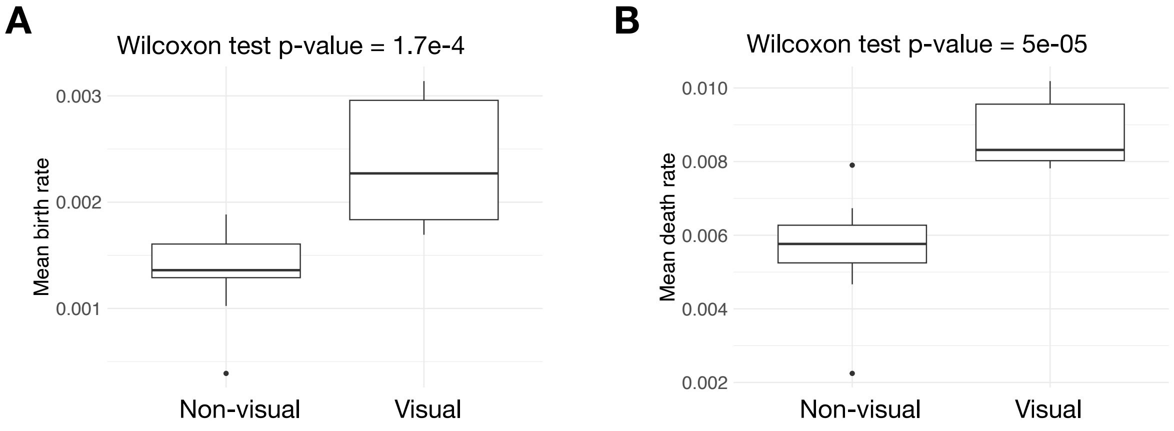


**Supplementary Figure 39.** Distribution of (A) mean birth and (B) mean death rates on non-visual and visual opsin subfamilies. Wilcoxon test p-values between non-visual and visual rates are indicated above each plot.

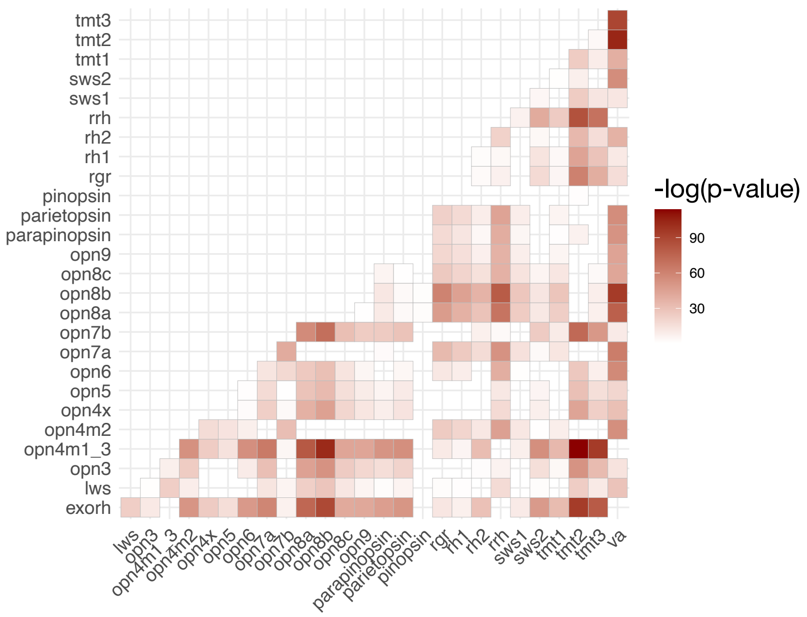


**Supplementary Figure 40.** Pairwise Wilcoxon test p-values between the distributions of dN/dS-values per subfamily. P-values are corrected for multiple testing using the Bonferroni method. Non-significant Bonferroni p-values (> 0.05) are indicated as blank squares while all significant Bonferroni p-values are colored (< 0.05).

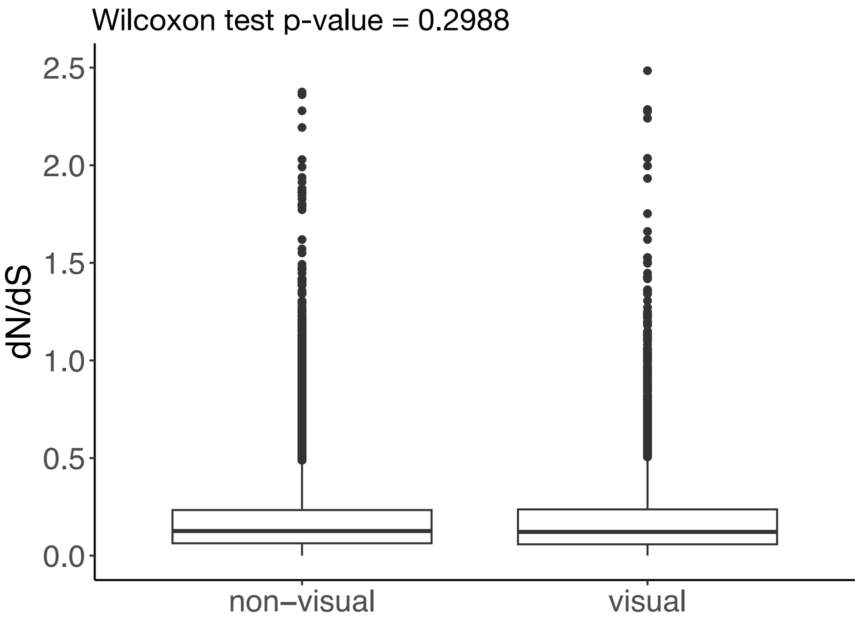


**Supplementary Figure 41.** Distribution of dN/dS computed on non-visual opsins and visual opsins. The Wilcoxon test p-value is indicated.

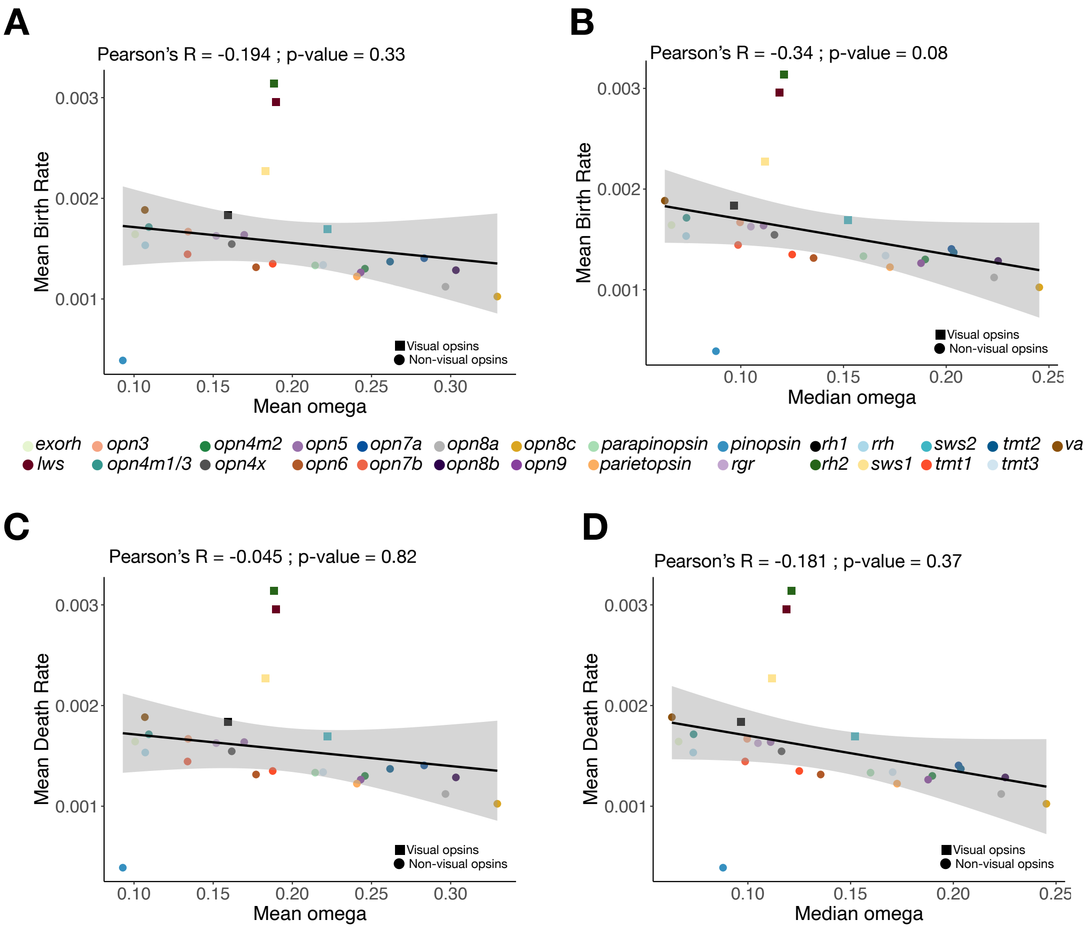


**Supplementary Figure 42.** Relationship between the mean birth rate per subfamily and (A) the mean dN/dS or (B) the median dN/dS. Relationship between the mean death rate per subfamily and (C) the mean dN/dS or (D) the median dN/dS. The solid black lines represent the linear regression lines, and the grey areas represent confidence intervals. Pearson correlation coefficients and p-values are indicated for each comparison.

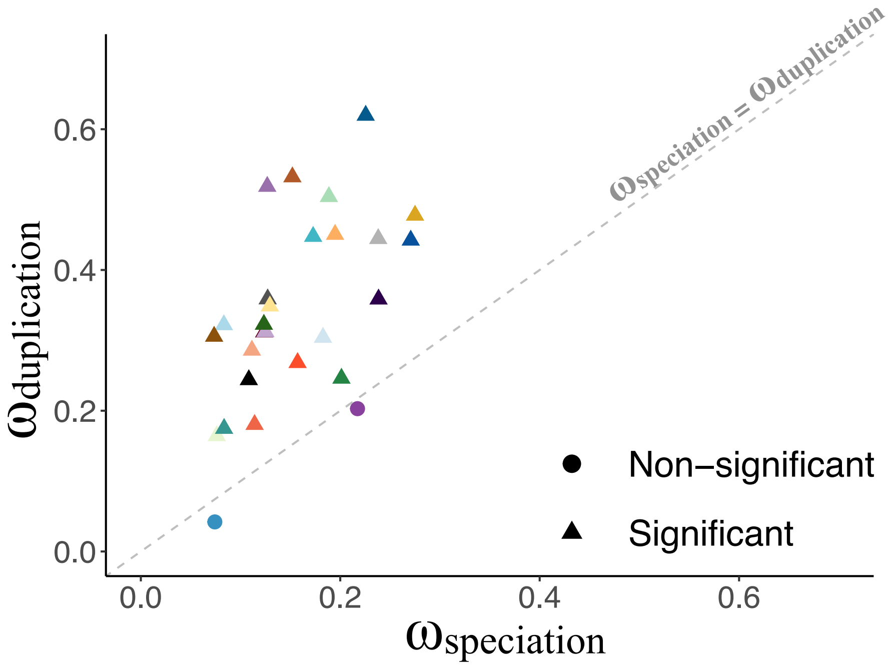


**Supplementary Figure 43.** Mean ω on branches resulting from duplication ($\omega_{duplication}$) against mean ω on branches resulting from speciation ($\omega_{speciation}$), for each subfamily, computed with BUSTED (14). Triangles represent subfamilies for which $\omega_{duplication}$ was significantly higher than $\omega_{speciation}$ and circles represent subfamilies for which $\omega_{duplication}$ and $\omega_{speciation}$ were not significantly different. The gray dashed line represents a slope of 1 ($\omega_{duplication} = \omega_{speciation}$).

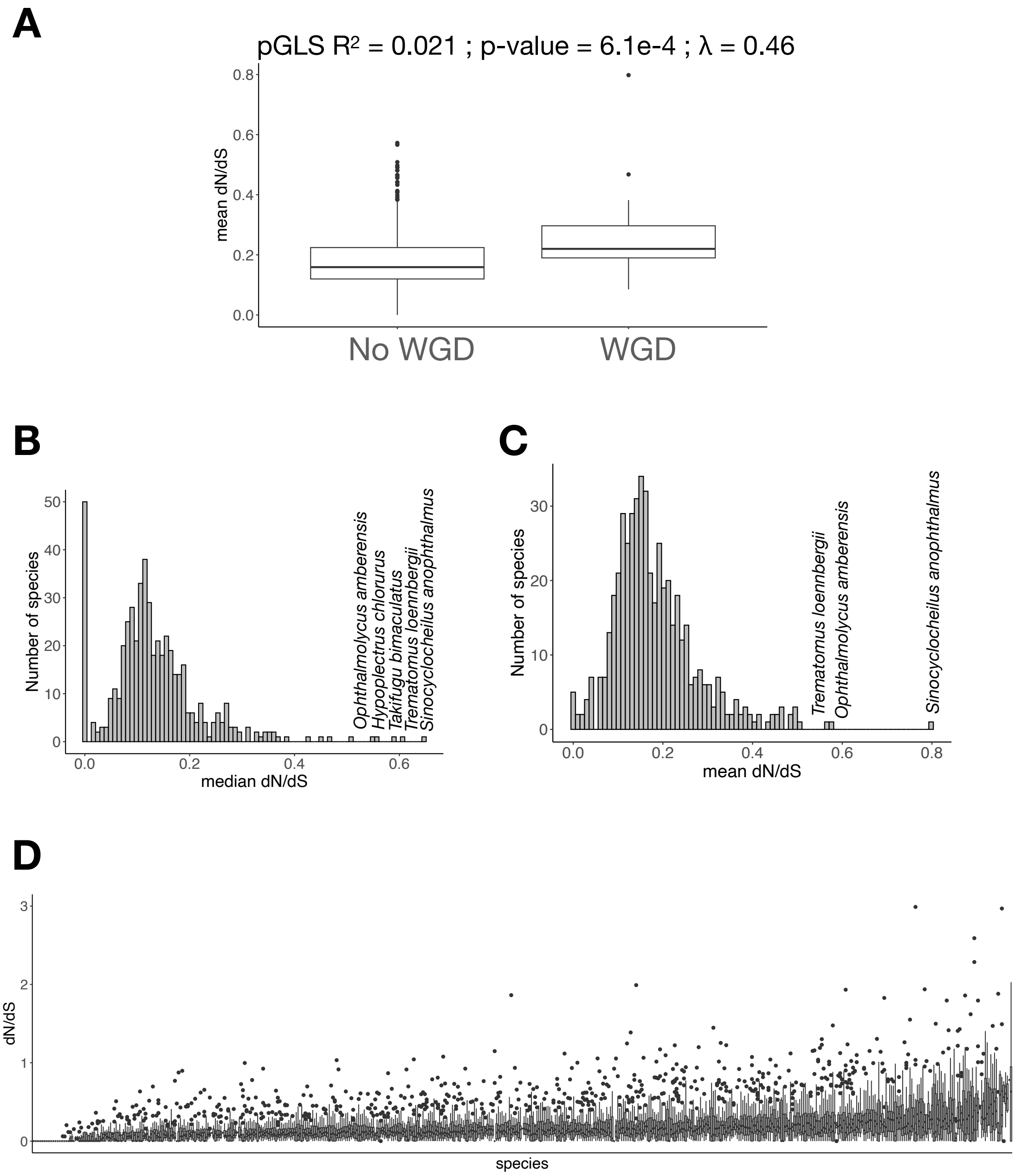


**Supplementary Figure 44.** dN/dS in opsin genes. (A) Comparison of the mean dN/dS of opsin genes for diploid and tetraploid teleosts. pGLS $R^2$, p-value and $\lambda$ are indicated. Distribution of (B) the median or (C) the mean opsin dN/dS per species. Species with elevated mean or median dN/dS are indicated. (D) Distribution of the dN/dS per opsin gene for each species.

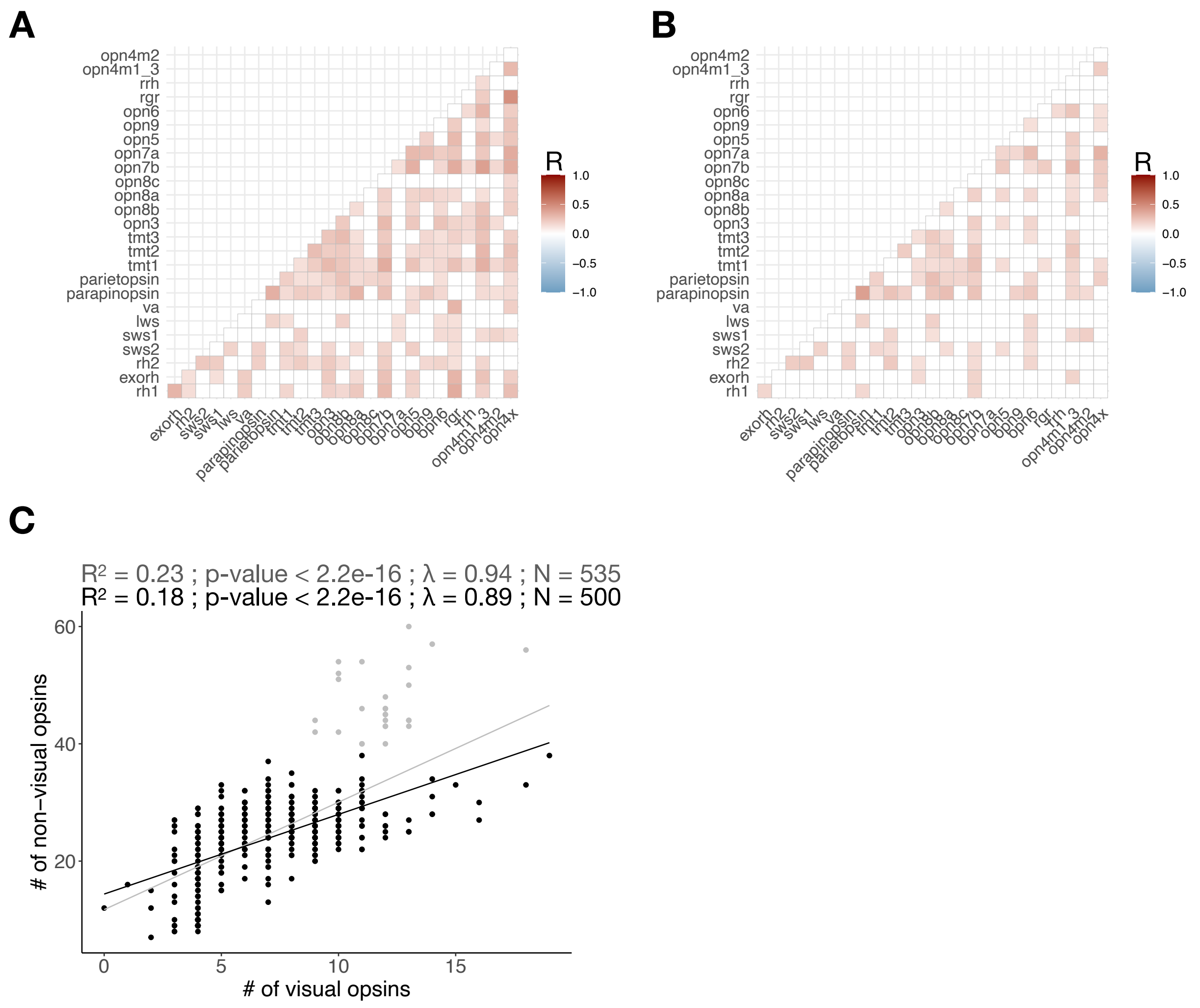


**Supplementary Figure 45.** (A) Correlation matrix of the number of genes per subfamily considering (A) all ray-finned fishes or (B) only diploid teleost species. Only pGLS with a Bonferroni corrected p-value < 0.05 are colored, and the color corresponds to the pGLS R value. (C) Correlation between the number of non-visual opsins and the number of visual opsins. pGLS $R^2$, p-value and λ and regression lines considering all ray-finned fishes (gray), or only diploid teleosts (black) are indicated. N represents the number of species considered in both pGLS.

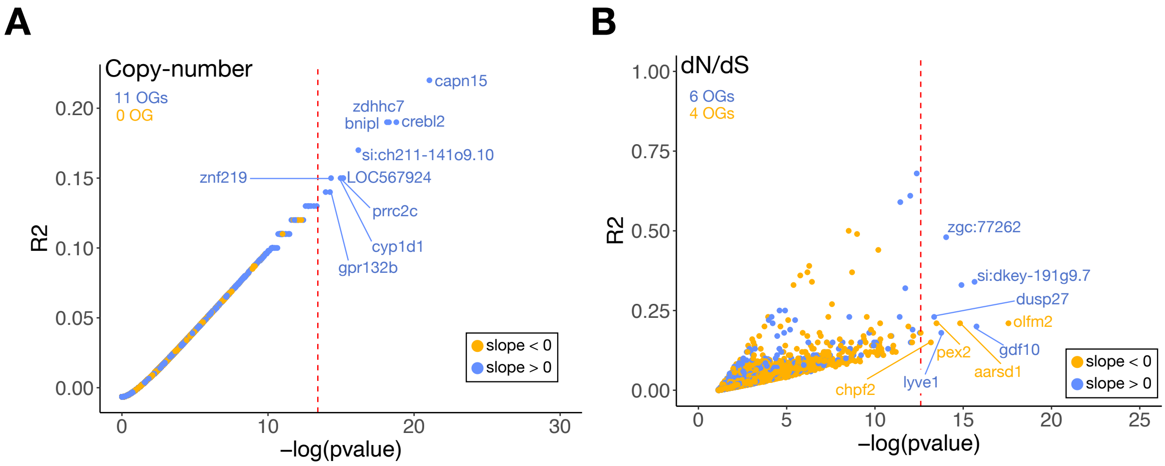


**Supplementary Figure 46.** Genome-wide screen for associations between the number of visual opsins and (A) the number of genes in 33,018 orthogroups and (B) the mean ω per OG, for 14,740 OGs. The dashed red lines correspond to the Bonferroni significance threshold. The number of positively correlated OGs (blue) and negatively correlated OGs (yellow) are indicated for both analyses. Names of the most significant OGs are indicated. The names assigned to an OG correspond to the name of one zebrafish gene name inside this OG. OGs with no zebrafish gene were not included in enrichment analysis and are not discussed in this study.

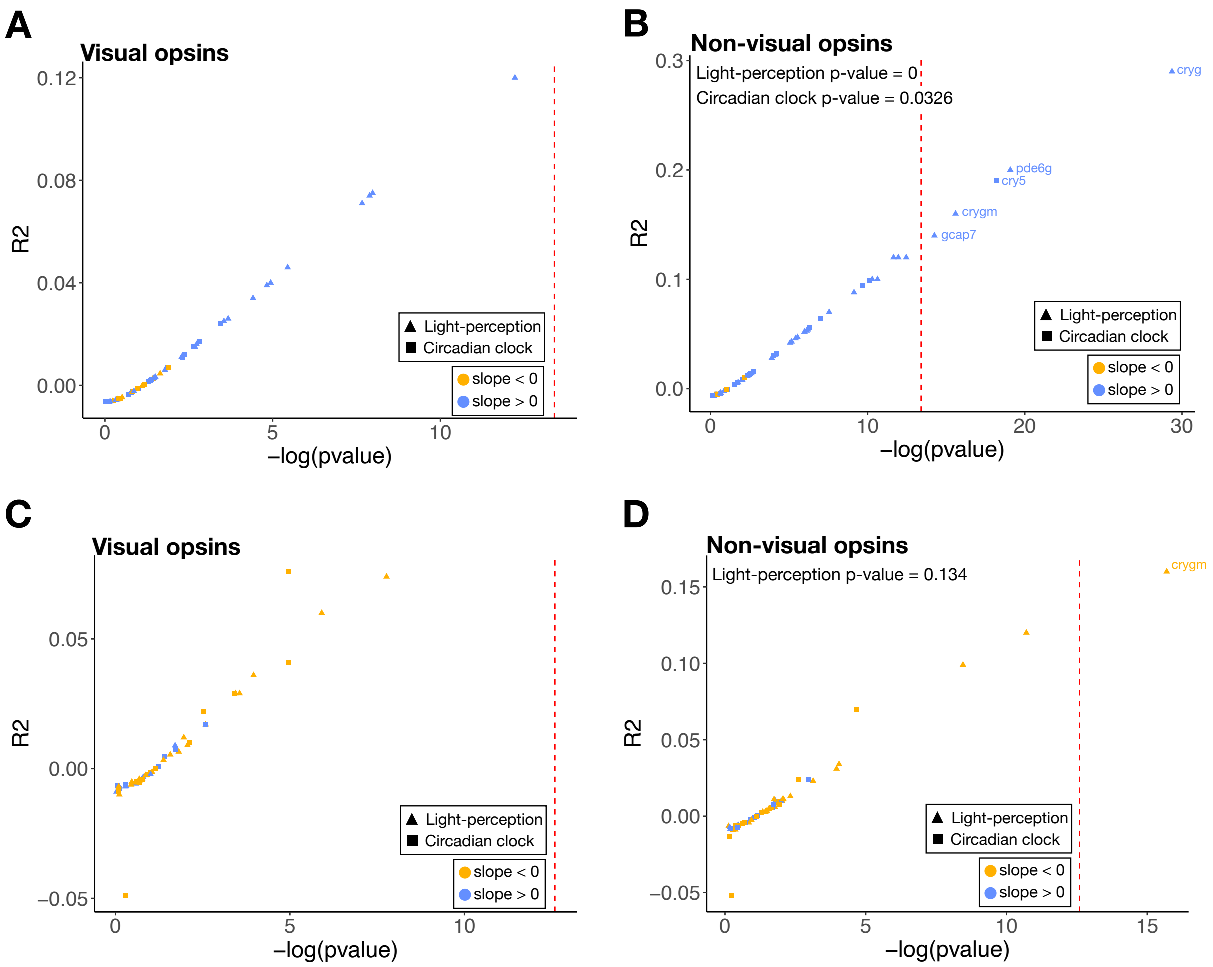


**Supplementary Figure 47.** Genome-wide screen for associations between the number of genes in 33,018 orthogroups and (A) the number of visual opsins or (B) the number of non-visual opsins. (C) and (D) represent the genome wide screen between the mean ω per OG, for 14,740 OGs, and the number of visual or non-visual opsins, respectively. Only genes involved in light-perception or in the circadian clock regulation are shown. The dashed red lines correspond to the Bonferroni significance threshold. Names of the significant OGs are indicated. The probabilities of finding 4 light-perception or 1 circadian clock OG associated to the number of non-visual opsins are indicated (p-value based on 10,000 simulations).

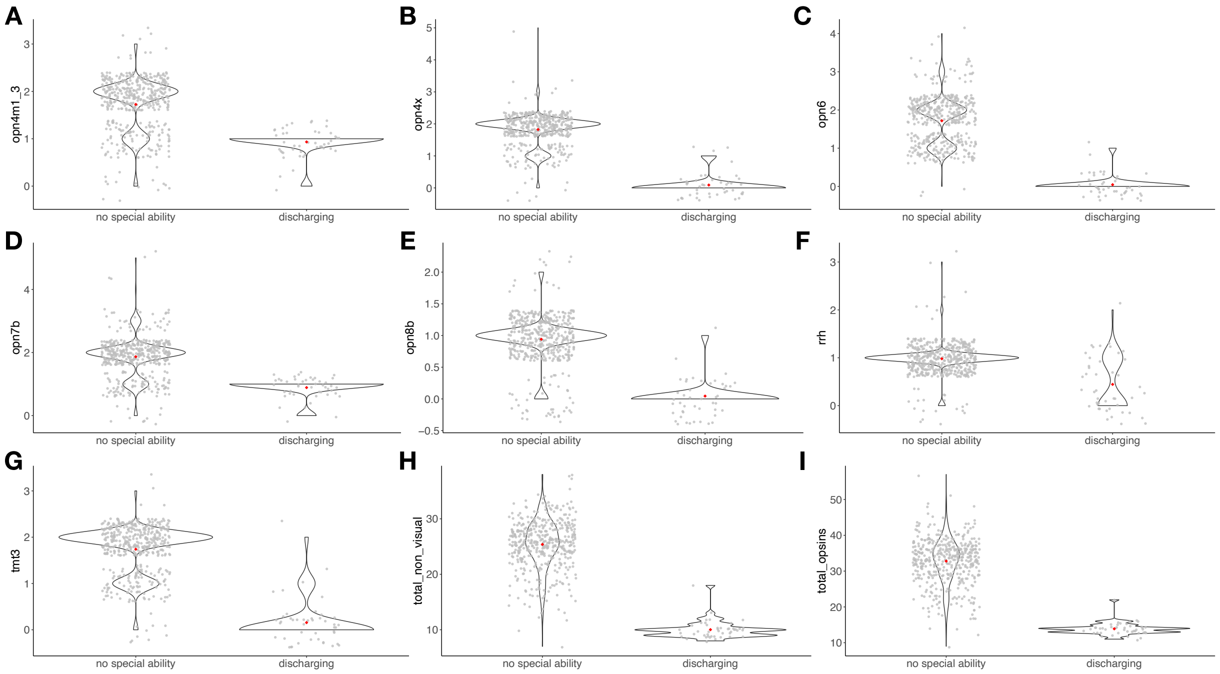


**Supplementary Figure 48.** Violin plots showing the number of complete genes in (A) *opn4m1/3*, (B) *opn4x*, (C) *opn6*, (D) *opn7b*, (E) *opn8b*, (F) *rrh*, (G) *tmt3* subfamily per electrogenic category. Violin plots for the total number of non-visual opsins and opsins (visual and non-visual) are represented in (H) and (I), respectively. The mean number of genes for each electrogenic category is indicated by a red diamond, while each species is represented by a gray dot. Only subfamilies significantly associated to electrogenic capacities are shown. The pGLS values of all subfamilies can be found in Supplementary data 1.

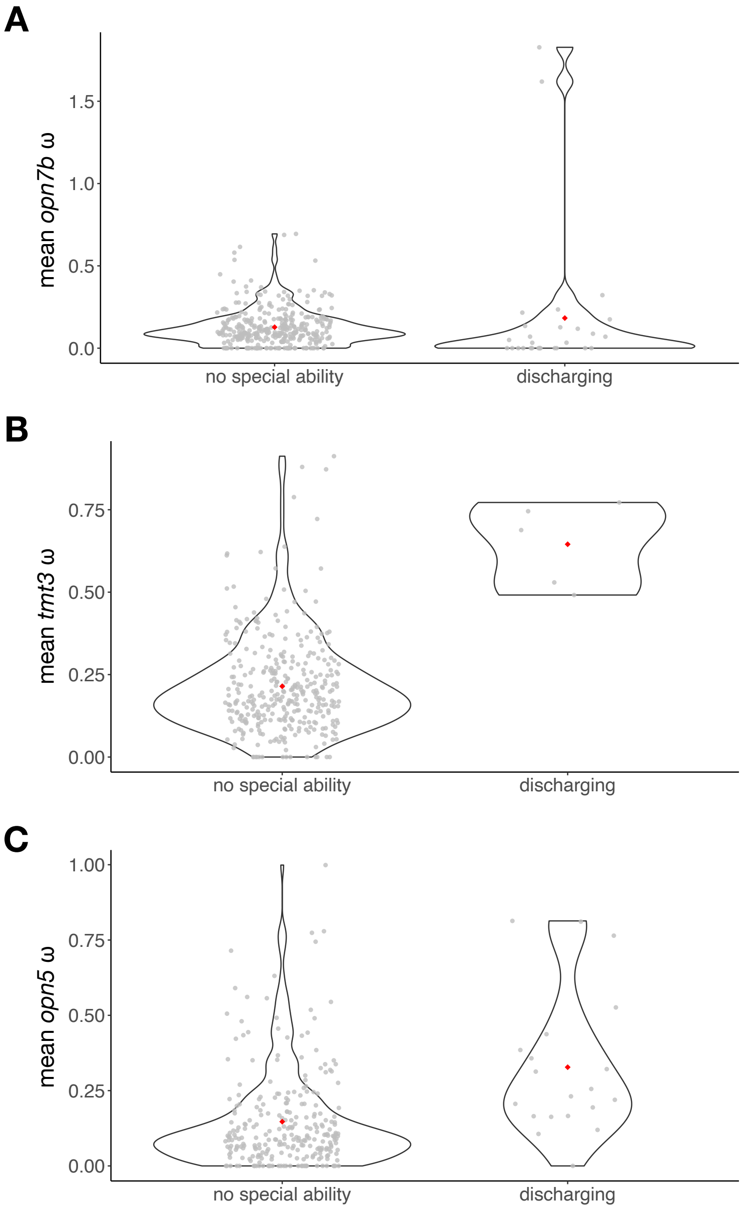


**Supplementary Figure 49.** Violin plots showing the mean ω of complete genes for (A) *opn7b*, (B) *tmt3*, (C) *opn5* subfamilies per depth category. The mean ω-value for each electrogenic category is indicated by a red diamond, while each species is represented by a gray dot. Only subfamilies significantly associated with electrogenic capacities are shown. The pGLS values of all subfamilies can be found in Supplementary data 1.

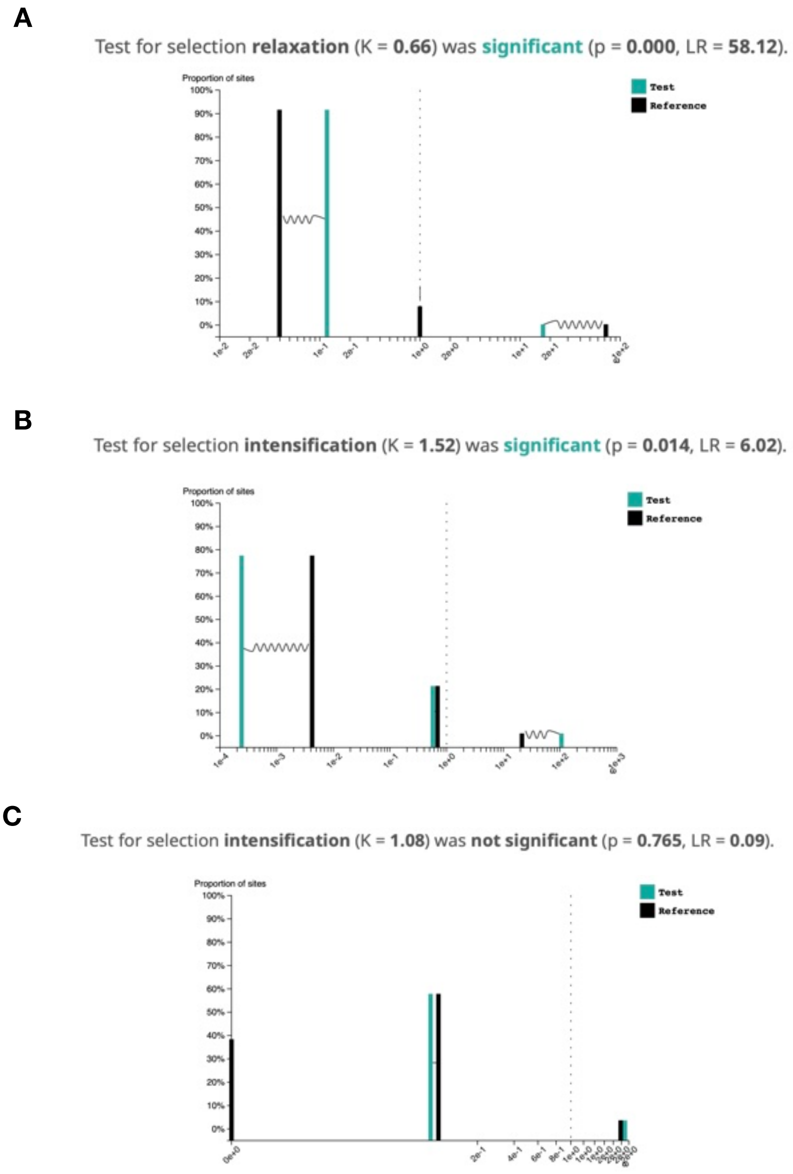


**Supplementary Figure 50.** Results of RELAX, assigning electric species opsins as test branches and all other terminal branches as foreground, for (A) *opn5*, (B) *tmt3,* and (C) *opn7b*.

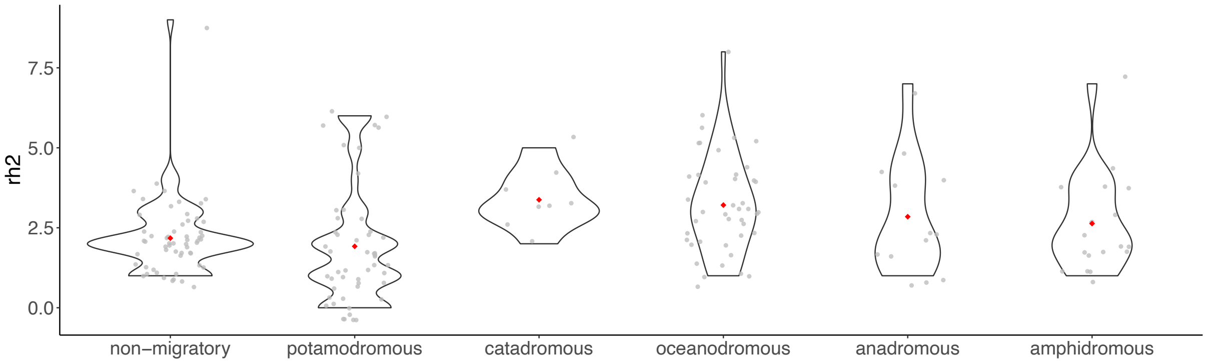


**Supplementary Figure 51.** Violin plots showing the number of complete genes in the *rh2* subfamily per migratory behavior category. The mean number of genes per migratory behavior category is indicated by a red diamond, while each species is represented by a gray dot. *The rh2* subfamily was the only subfamily with a copy number associated to the migratory behavior.

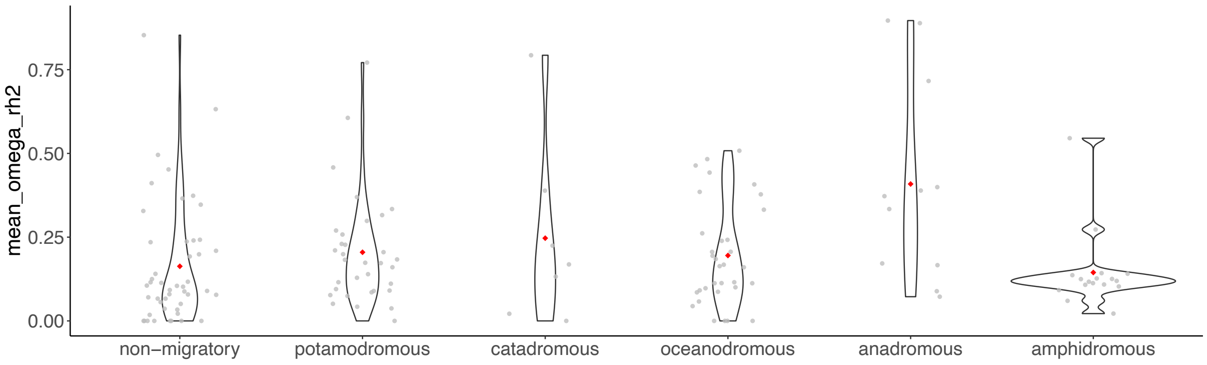


**Supplementary Figure 52.** Violin plots showing the mean ω of the *rh2* subfamily per migratory category. The mean ω-value for each category is indicated by a red diamond, while each species is represented by a gray dot. The pGLS values of all subfamilies can be found in Supplementary data 1.

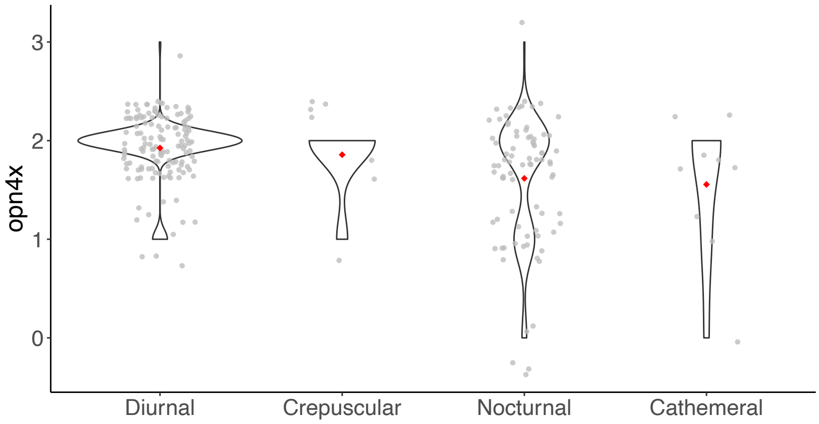


**Supplementary Figure 53.** Violin plots showing the number of complete genes in the *opn4x* subfamily per diel pattern category. The mean number of genes for each diel pattern is indicated by a red diamond, while each species is represented by a gray dot. *The opn4x* subfamily was the only subfamily with a copy number associated to the diel pattern.

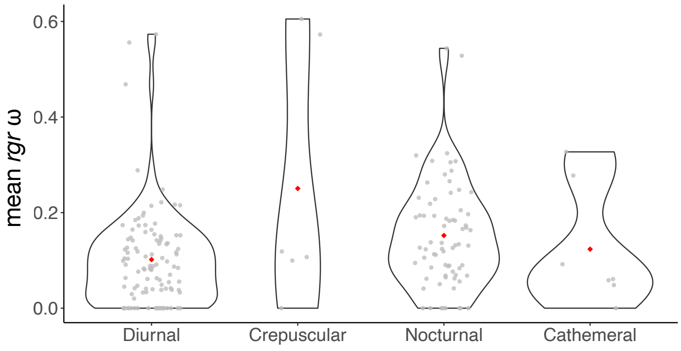


**Supplementary Figure 54.** Violin plots showing the mean ω of complete genes in the *rgr* subfamily per diel pattern category. The mean ω-value for each diel pattern is indicated by a red diamond, while each species is represented by a gray dot. *The rgr* subfamily was the only subfamily with a dN/dS associated to the diel pattern.

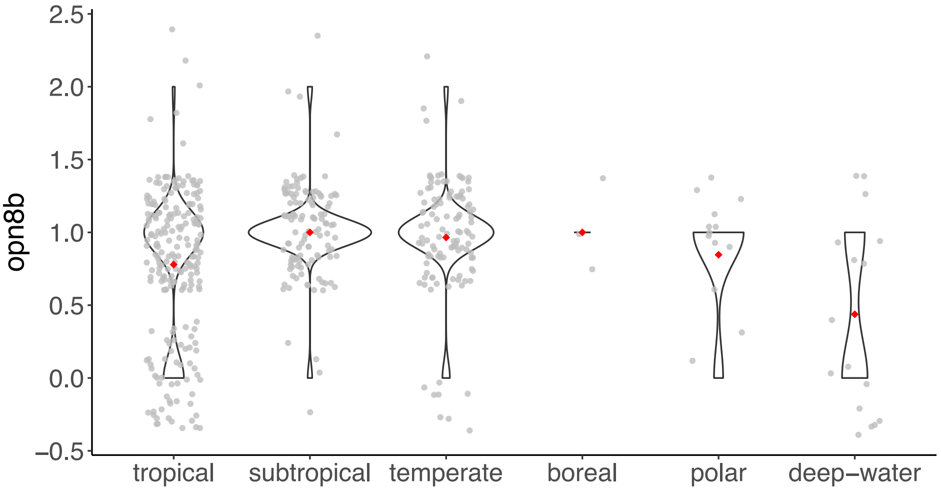


**Supplementary Figure 55.** Violin plots showing the number of complete genes in the *opn8b* subfamily per climate zone category. The mean number of genes for each climate zone is indicated by a red diamond, while each species is represented by a gray dot. The *opn8b* subfamily was the only subfamily with a copy number associated to the climate zone.

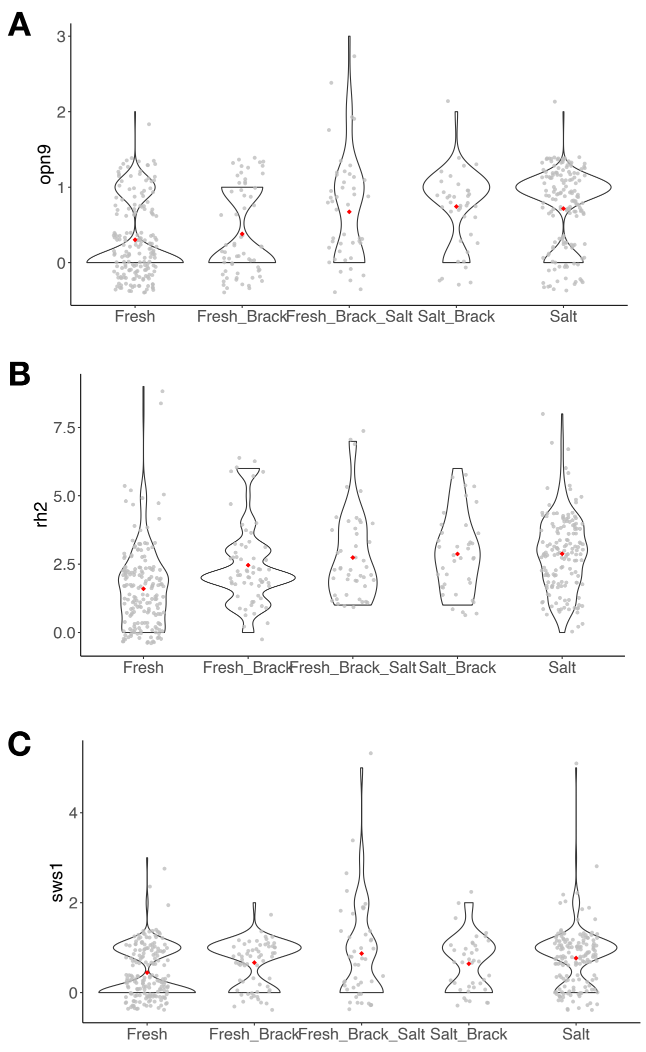


**Supplementary Figure 56.** Violin plots showing the number of complete genes in (A) *opn9*, (B) *rh2*, and (C) *sws1* per habitat category. The mean number of genes for each habitat is indicated by a red diamond, while each species is represented by a gray dot. Only subfamilies significantly associated to the habitat are shown. The pGLS values of all subfamilies can be found in Supplementary data 1.

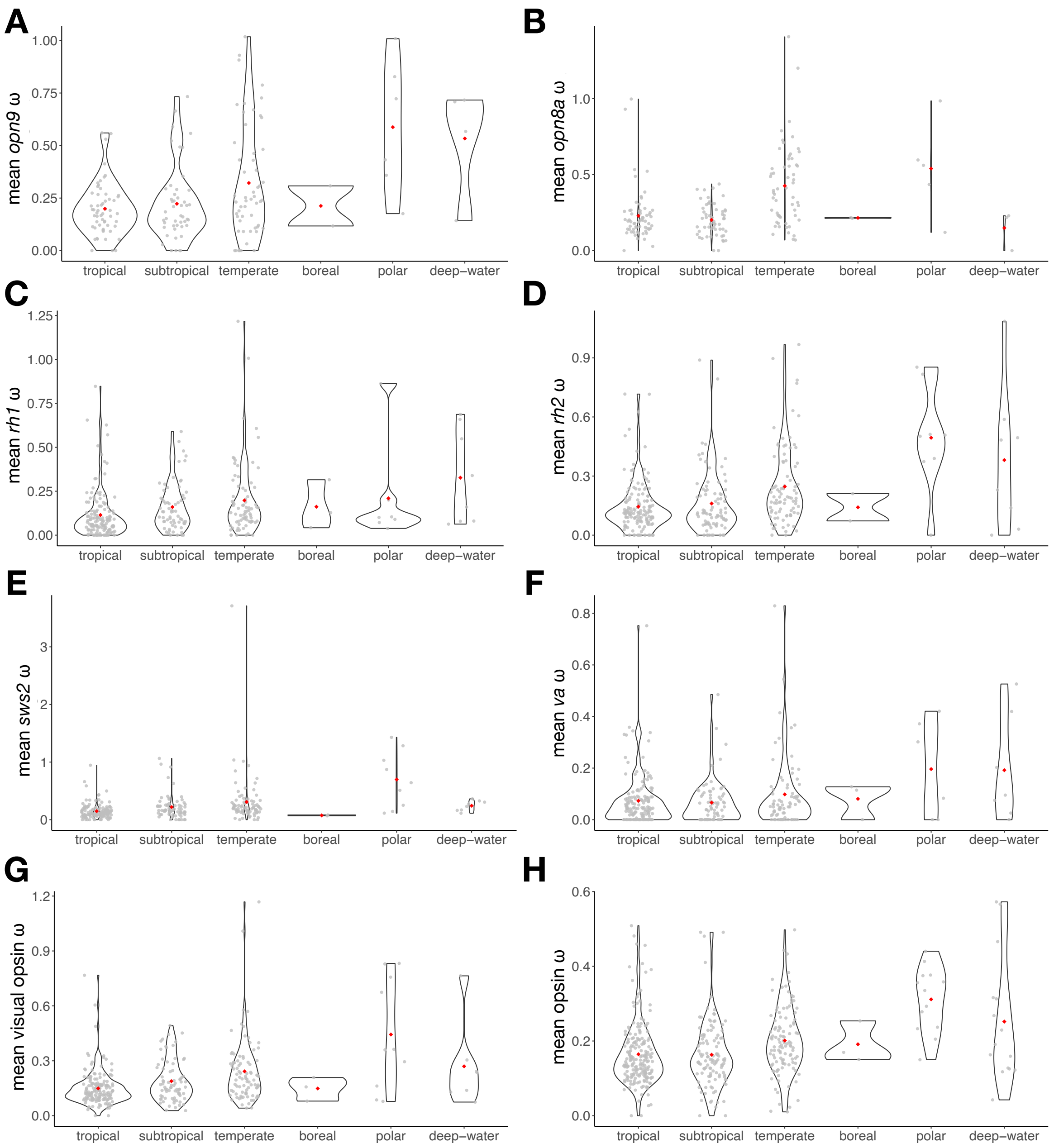


**Supplementary Figure 57.** Violin plots showing the mean ω of complete genes for (A) *opn9*, (B) *opn8a*, (C) *rh1*, (D) *rh2*, (E) *sws2*, (F) *va* subfamilies, (G) the entire visual opsin gene repertoire, and (H) the entire opsin gene repertoire, per climate zone category. The mean ω-value for each climate zone is indicated by a red diamond, while each species is represented by a gray dot. Only subfamilies significantly associated with the climate zone are shown. The pGLS values of all subfamilies can be found in Supplementary data 1. The high *rh1* ω observed for polar species is in agreement with a previous study, showing that *rh1* genes were under strong diversifying selection in Antarctic icefishes (15).

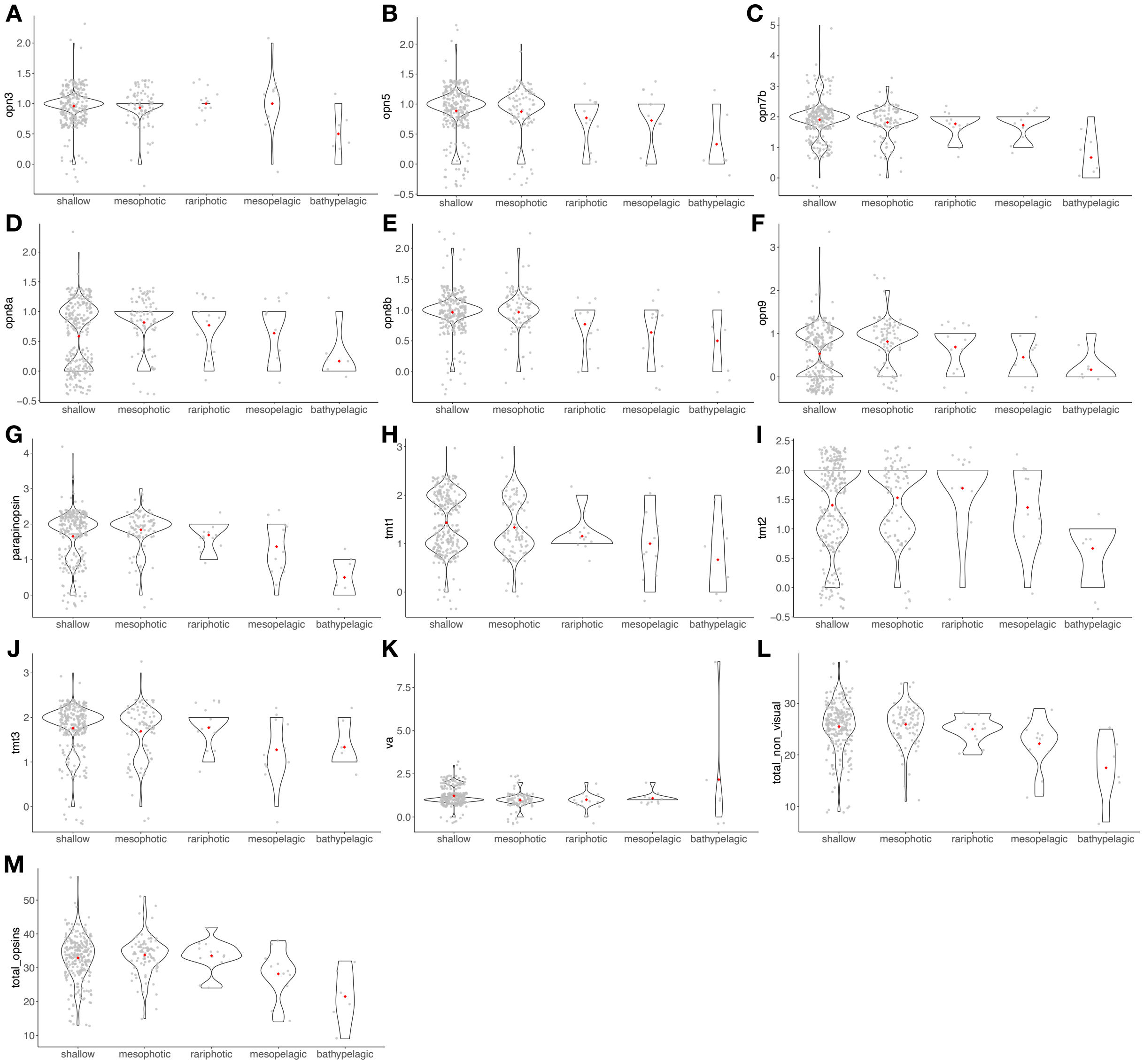


**Supplementary Figure 58.** Violin plots showing the number of complete genes in (A) *opn3*, (B) *opn5*, (C) *opn7b*, (D) *opn8a*, (E) *opn8b*, (F) *opn9*, (G) *parapinopsin*, (H) *tmt1*, (I) *tmt2*, (J) *tmt3*, (K) *va* subfamily per depth category. Violin plots for the total number of non-visual opsin genes and total opsins (visual and non-visual) are represented in (L) and (M), respectively. The mean number of genes for each depth category is indicated by a red diamond, while each species is represented by a gray dot. Only subfamilies significantly associated with depth are shown. The pGLS values for all subfamilies can be found in Supplementary data 1.

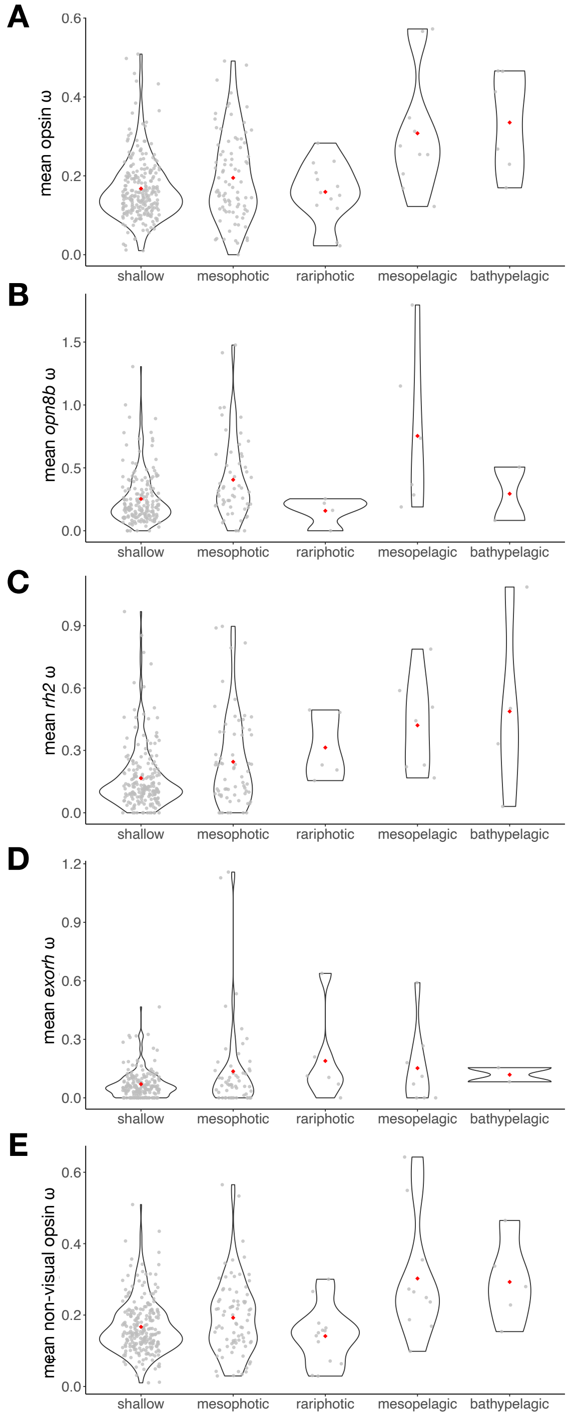


**Supplementary Figure 59.** Violin plots showing the mean ω of complete genes for (A) the entire opsin gene repertoire, the (B) *opn8b*, (C) *rh2*, (D) *exorh* subfamilies, and (E) for the entire non-visual opsin repertoire, per depth category. The mean ω-value for each depth category is indicated by a red diamond, while each species is represented by a gray dot. Only subfamilies significantly associated with depth are shown. The pGLS-values of all subfamilies can be found in Supplementary data 1.

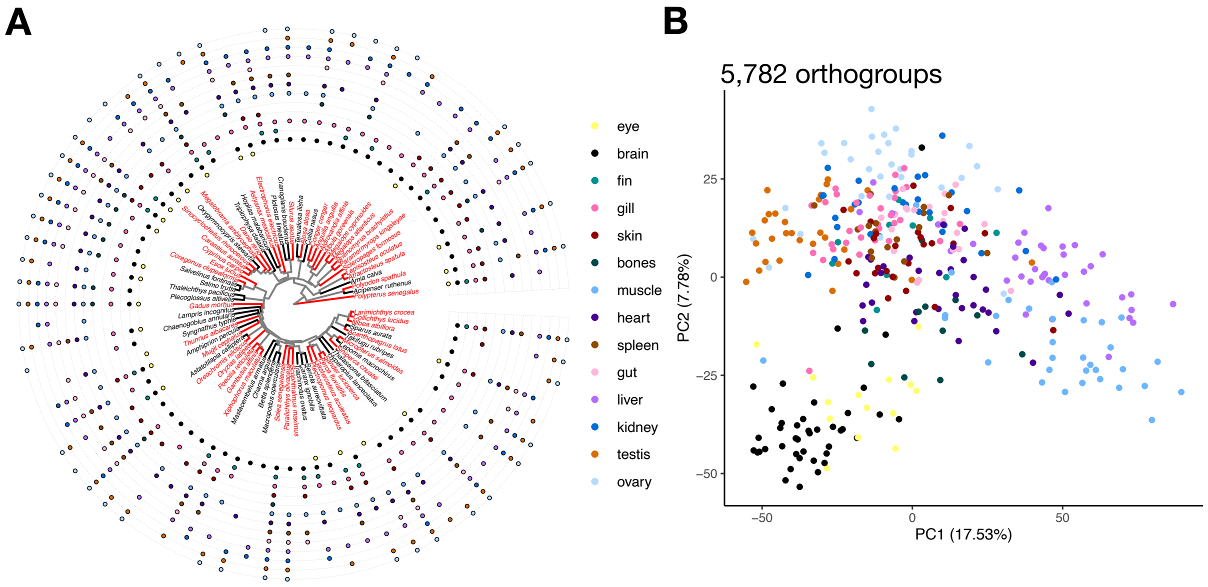


**Supplementary Figure 60.** RNA-seq analyses. (A) Phylogenetic tree representing the 75 ray-finned fish species for which at least one transcriptome was retrieved in the SRA database. Red branches correspond to species for which the genome is annotated on NCBI. The presence/absence of a tissue transcriptome for each species is indicated by the presence/absence of a dot, colored according to the tissue. (B) Principal component analysis (PCA) of samples from species with annotated genomes, using TPM values of 5,782 shared orthogroups. Samples are colored according to their tissue (358 samples, 14 tissues).

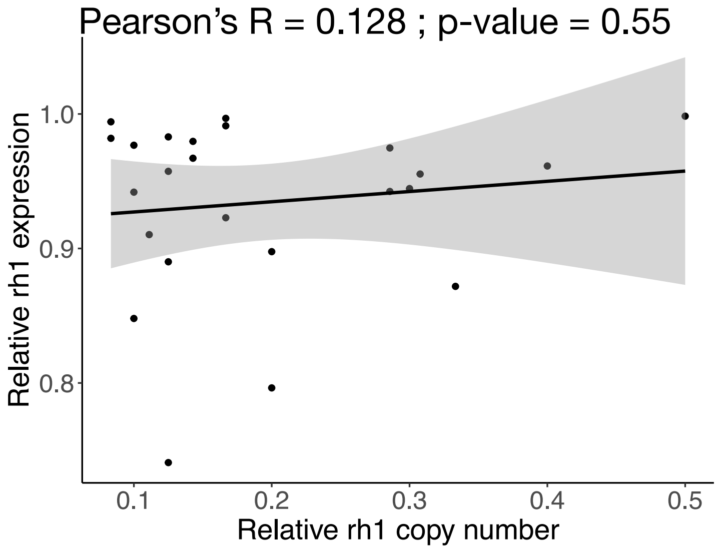


**Supplementary Figure 61.** Relationship between the relative expression of *rh1* and the relative *rh1* gene copy number (both relative to the entire visual opsin gene repertoire). The solid black line represents the linear regression line, and the grey area represents the confidence interval. Pearson correlation coefficient and p-value are indicated.

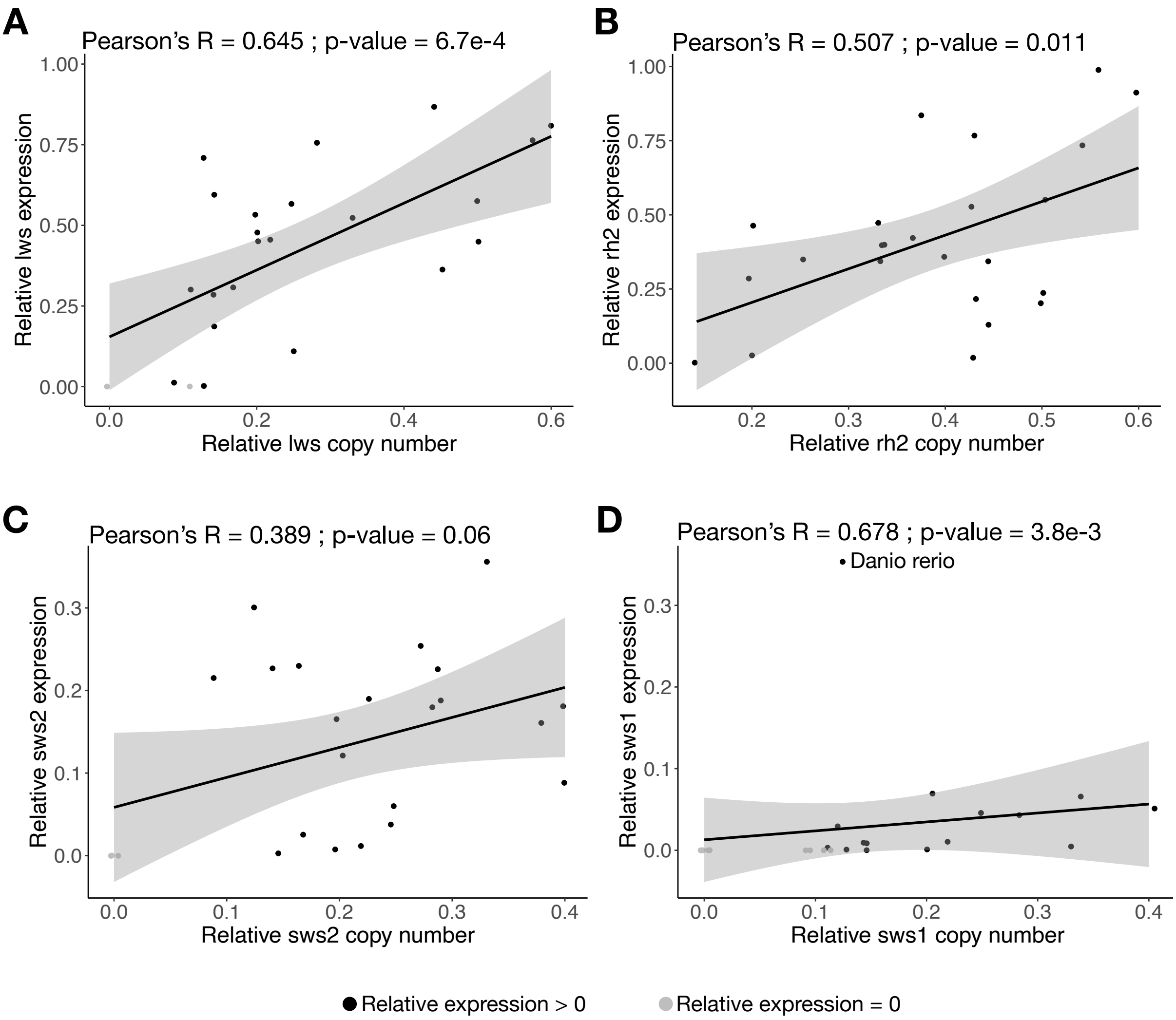


**Supplementary Figure 62.** Relationship between the relative expression of cone opsins and their relative gene copy number (relative to the entire cone visual opsin gene repertoire), for (A) *lws*, (B) *rh2*, (C) *sws2*, and (D) *sws1*. The solid black lines represent the linear regression lines, and the grey areas represent confidence intervals. Pearson correlation coefficients and p-values are indicated for each cone opsin. For *sws1*, we removed *D. rerio* from the correlation test, as it appeared as a clear outlier.

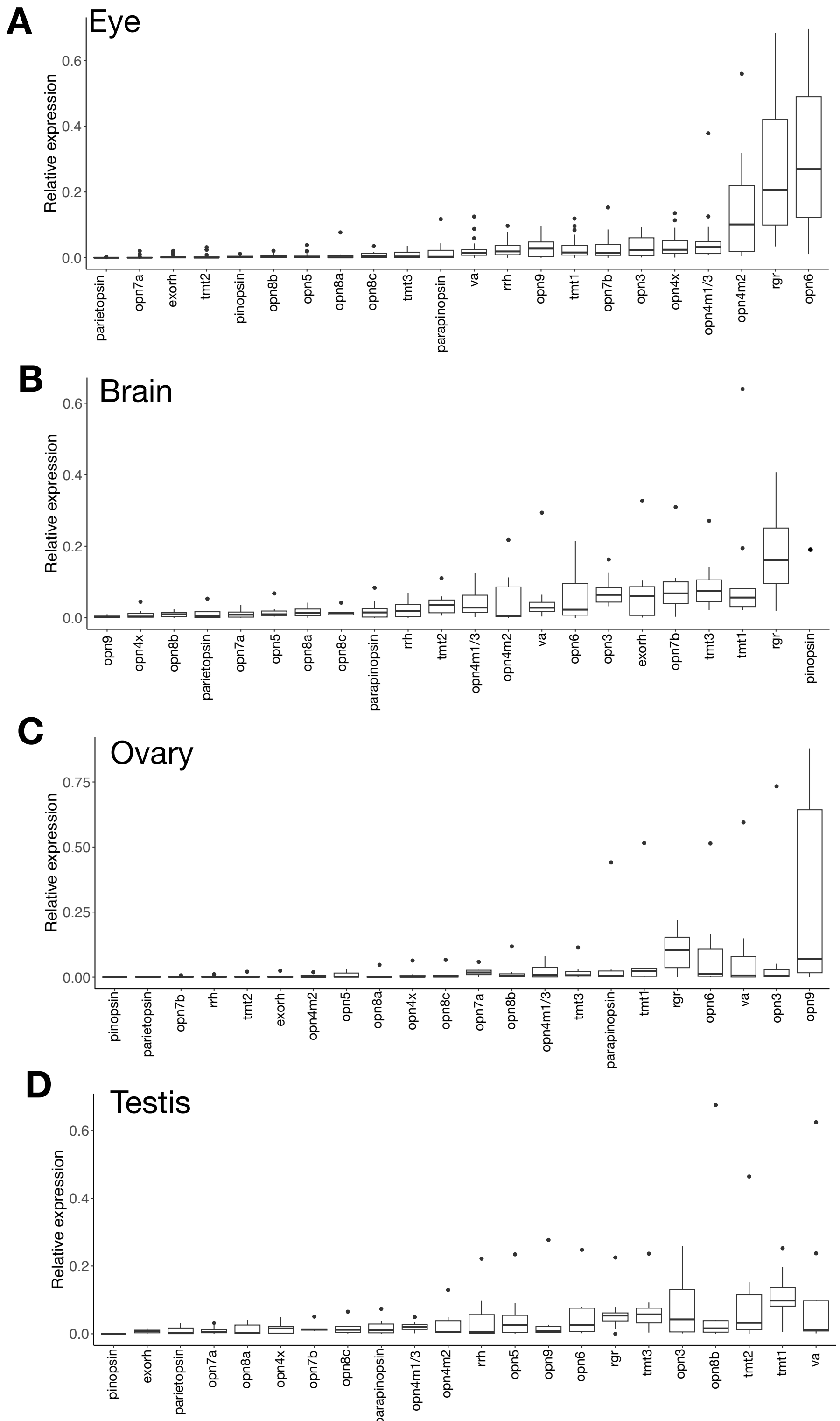


**Supplementary Figure 63.** Distribution of the expression of each non-visual opsin subfamily, relative to the entire non-visual opsin gene repertoire expression, in four tissues, (A) eye, (B) brain, (C) ovary, and (D) testis.

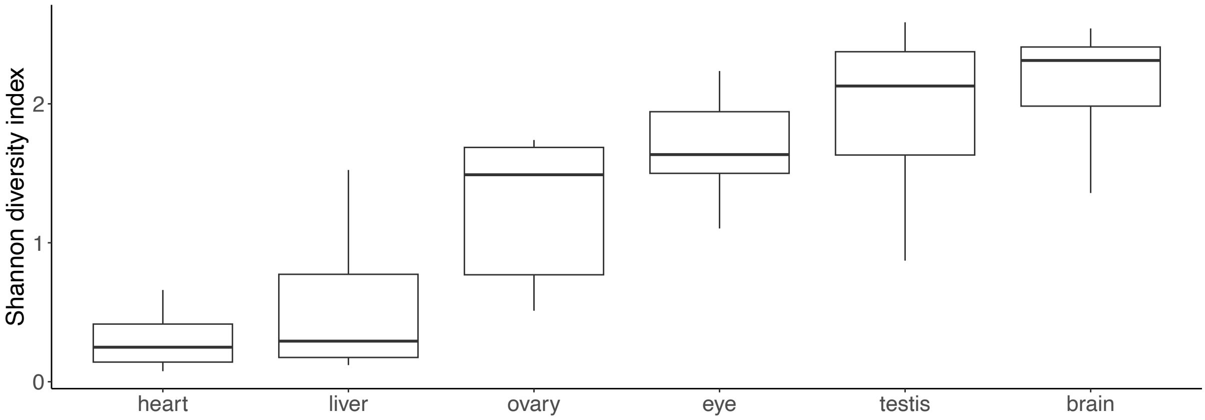


**Supplementary Figure 64.** Distribution of the Shannon diversity index for the 6 tissues investigated. Tissues are ordered according to the mean Shannon diversity index.

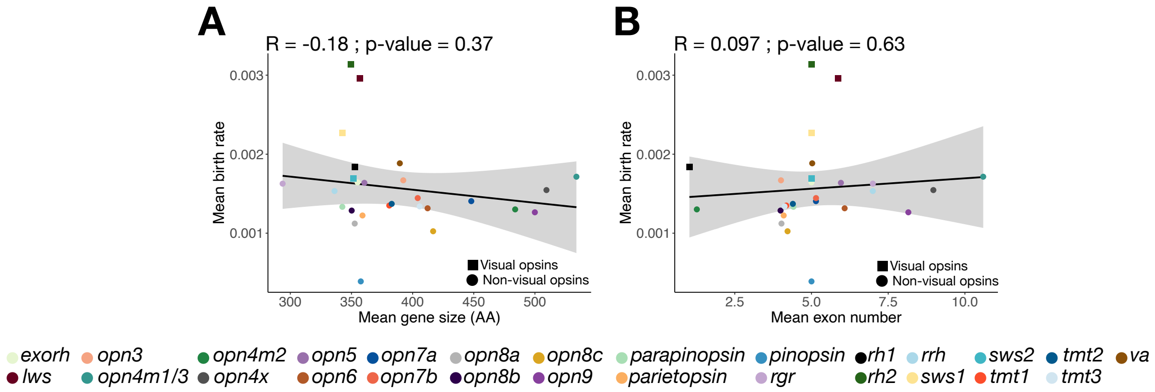


**Supplementary Figure 65.** Relationship between the mean birth rate per subfamily and (A) the mean gene size and (B) the mean number of exons. Pearson correlation coefficients and p-values are indicated. The solid black lines represent the linear regression lines, and the grey areas represent confidence intervals.

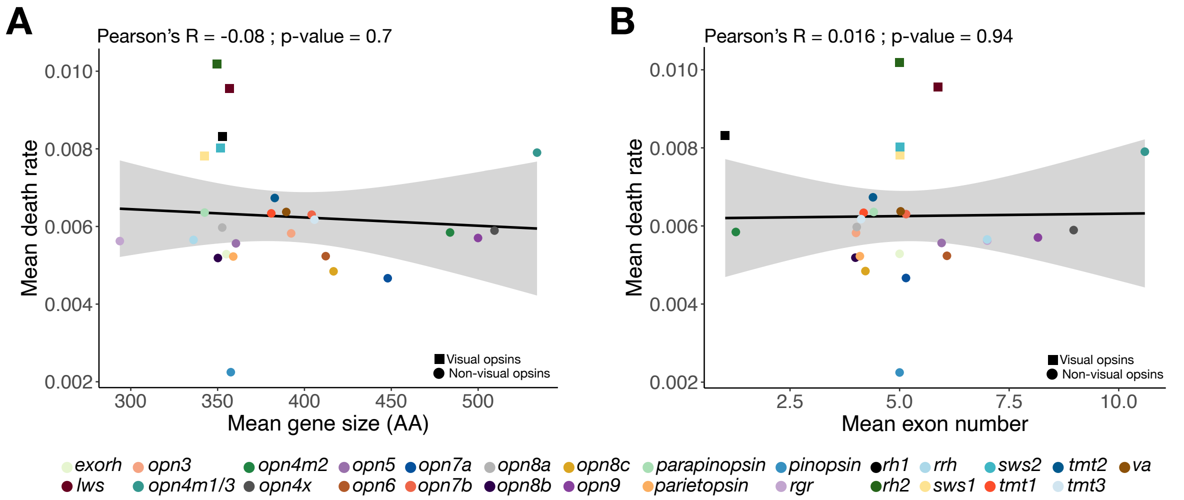


**Supplementary Figure 66.** Relationship between the mean death rate per subfamily and (A) the mean gene size and (B) the mean number of exons. Pearson correlation coefficients and p-values are indicated. The solid black lines represent the linear regression lines, and the grey areas represent confidence intervals.

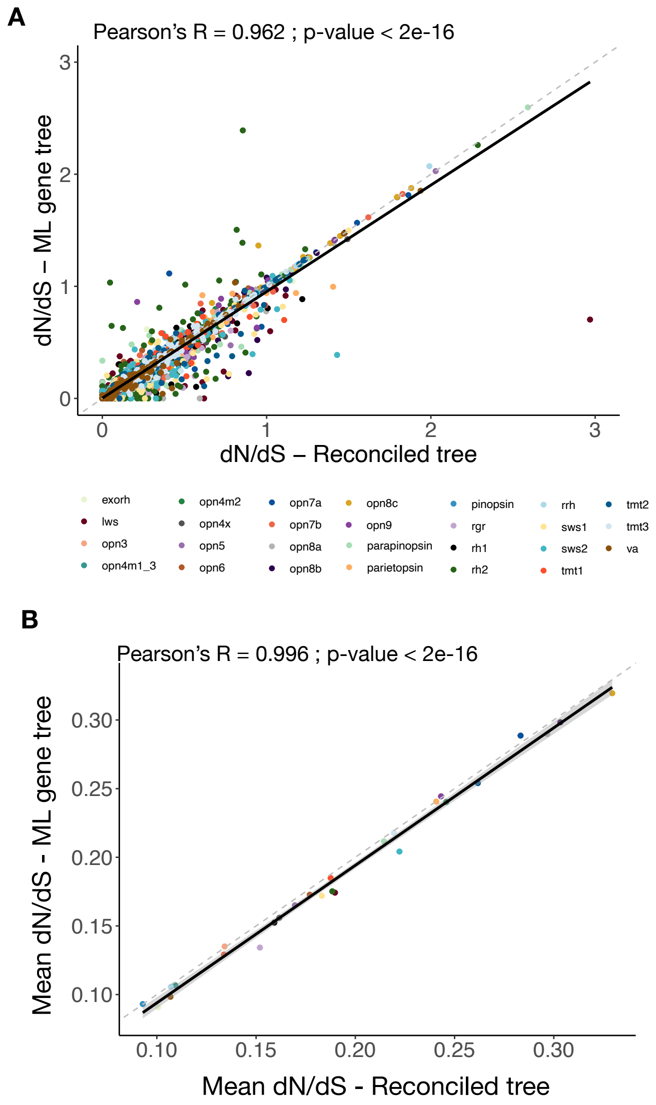


**Supplementary Figure 67.** Comparison of the dN/dS computed using maximum likelihood gene trees or reconciled gene trees. (A) Comparison of the dN/dS on all terminal branches. (B) Comparison of the mean dN/dS per opsin gene subfamily. Pearson correlation coefficients and p-values and linear regression equations are indicated. The solid black lines represent the linear regression lines, and the grey areas represent confidence intervals. Dashed lines indicate slope = 1 (x=y).

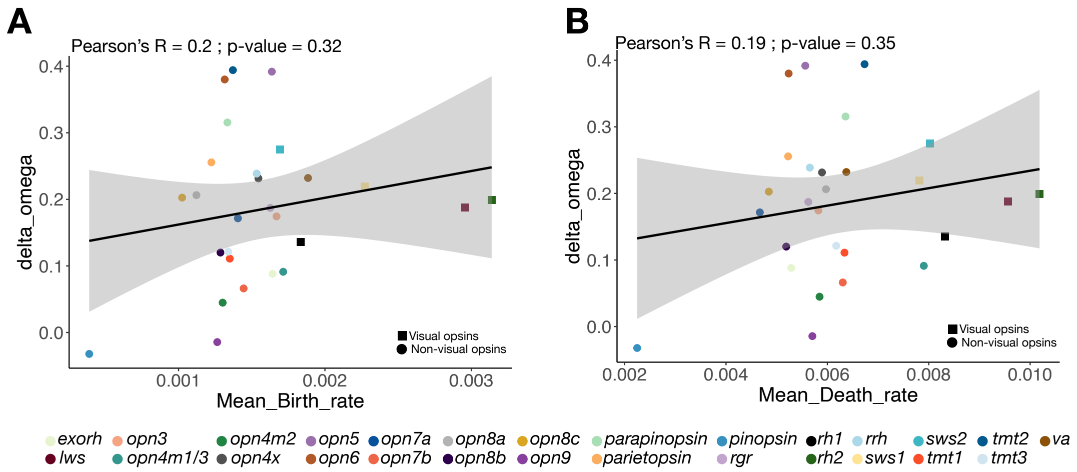


**Supplementary Figure 68.** Relationship between $\Delta_{\omega}$ ($\omega_{duplication} = \omega_{speciation}$) per opsin gene subfamily and (A) the mean birth rate and (B) the mean death rate. Pearson correlation coefficients and p-values are indicated. The solid black lines represent the linear regression lines, and the grey areas represent confidence intervals.

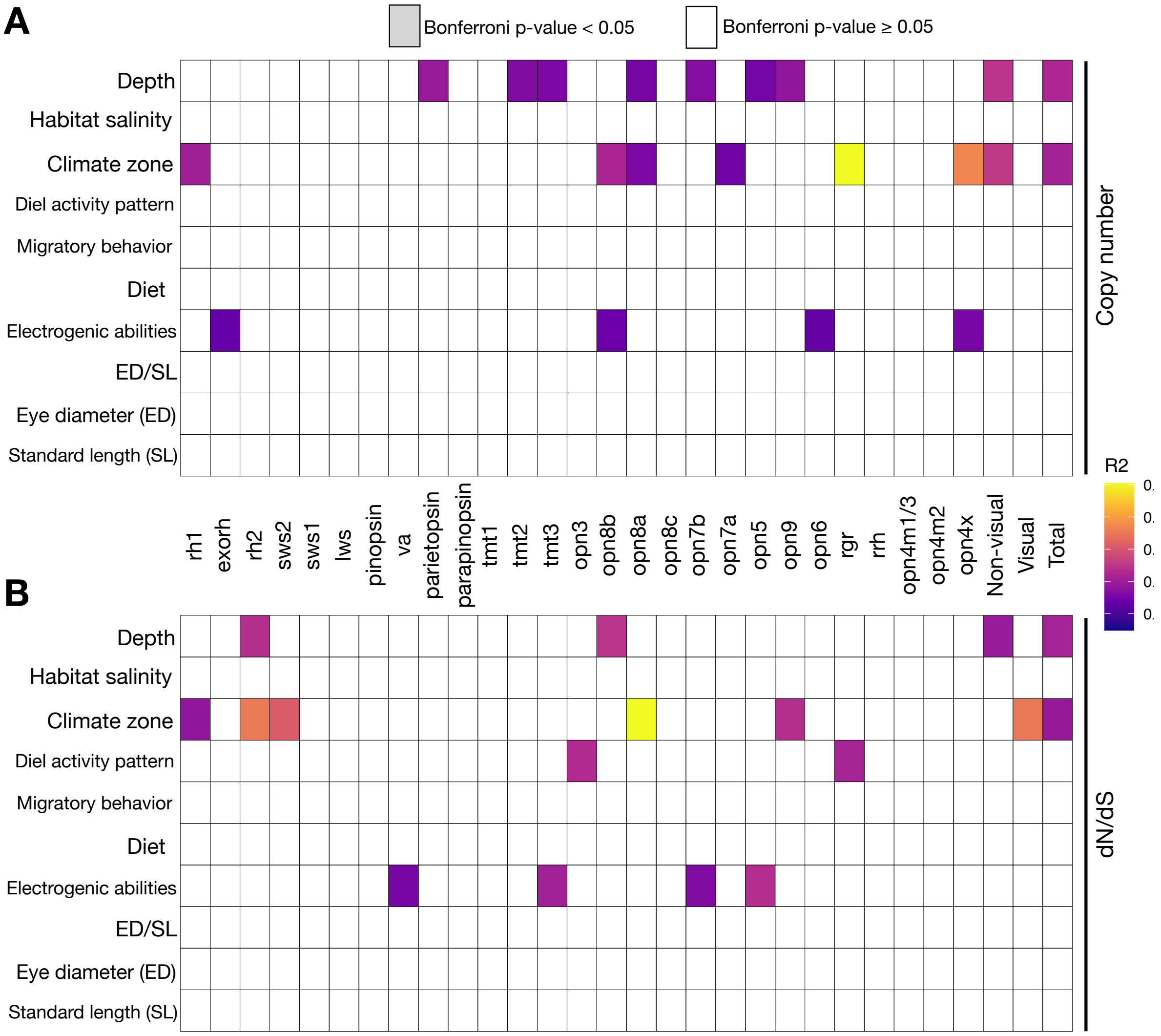


**Supplementary Figure 69.** Associations between eco-morphological factors and (A) the number of genes per opsin gene subfamily, or the total number of opsins, visual opsins or non-visual opsins. (B) the mean dN/dS per opsin gene subfamily, or the mean dN/dS of all opsins, visual opsins or non-visual opsins. For the mean dN/dS for the entire opsin gene repertoire or the entire non-visual/visual gene repertoire, we only considered species with at least three different opsins. Colored tiles correspond to significant associations (pGLS Bonferroni corrected p-value < 0.05) and the color depends on the pGLS $R^2$.

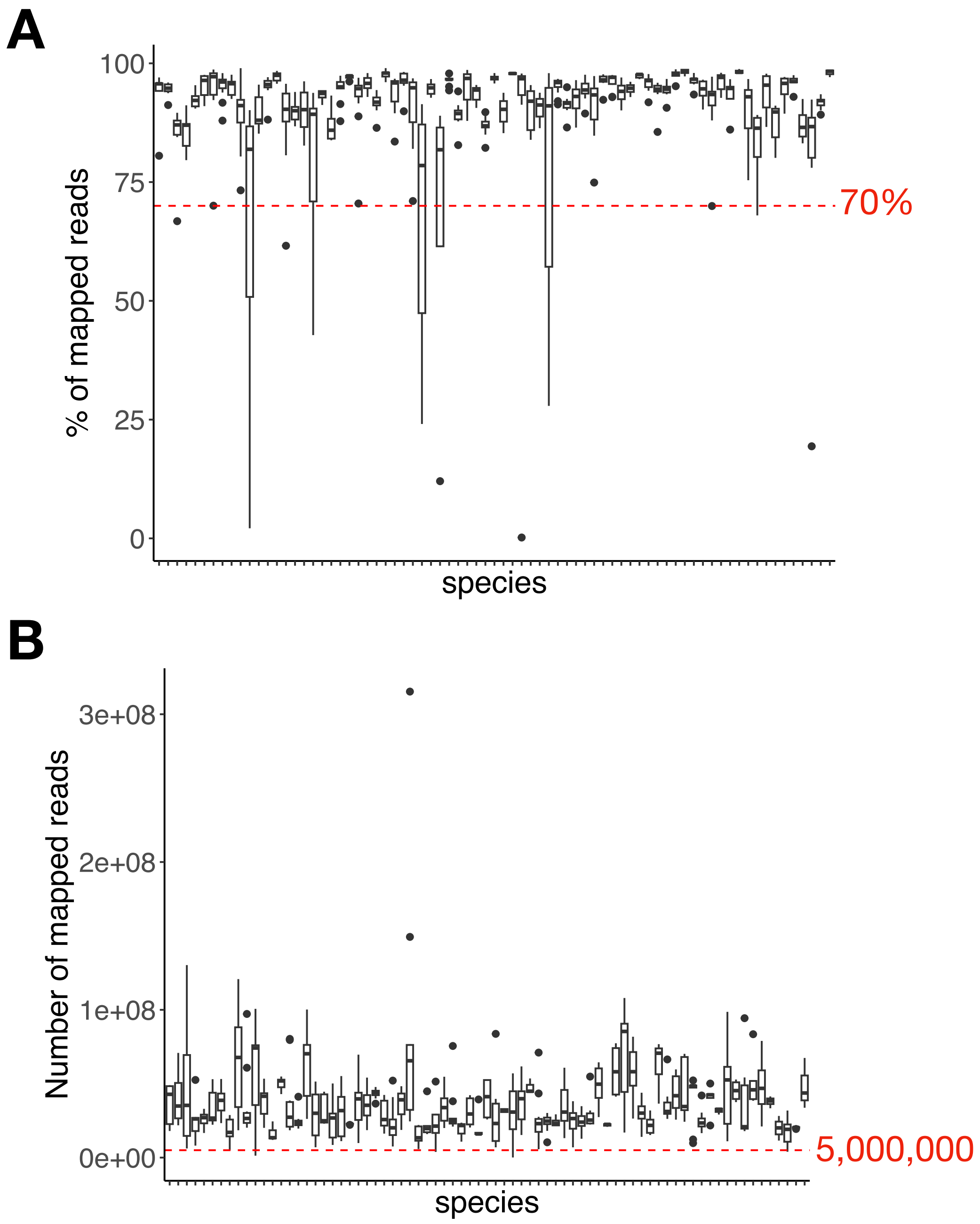


**Supplementary Figure 70.** RNA dataset quality filters. (A) Distributions of the percentage of mapped reads per tissue for each species. Samples with fewer than 70% of reads mapped to the genome were discarded. (B) Distributions of the number of mapped reads per tissue for each species. Samples with fewer than 5,000,000 mapped reads were discarded.

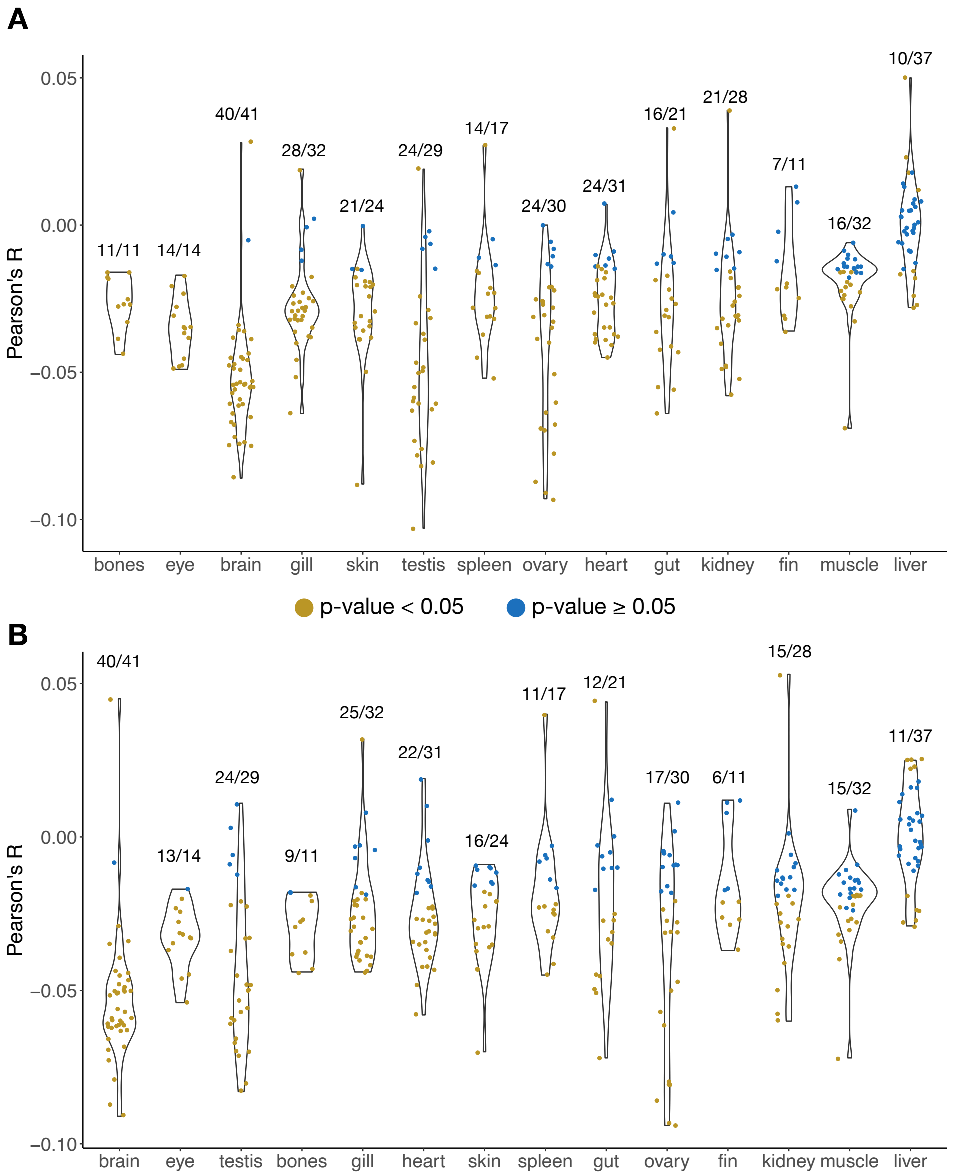


**Supplementary Figure 71.** For each sample, a Pearson correlation was performed between (A) genes TPM and ω and (B) OGs mean TPM and mean ω. For each tissue, samples with significant correlations are represented in orange and samples with non-significant correlations are represented in blue. The number of significant samples/total number of samples is indicated above each tissue. The vast majority of significant correlations were negative correlations. Results at the gene level (A) and OG level (B) were similar (see Supplementary Figure 72).

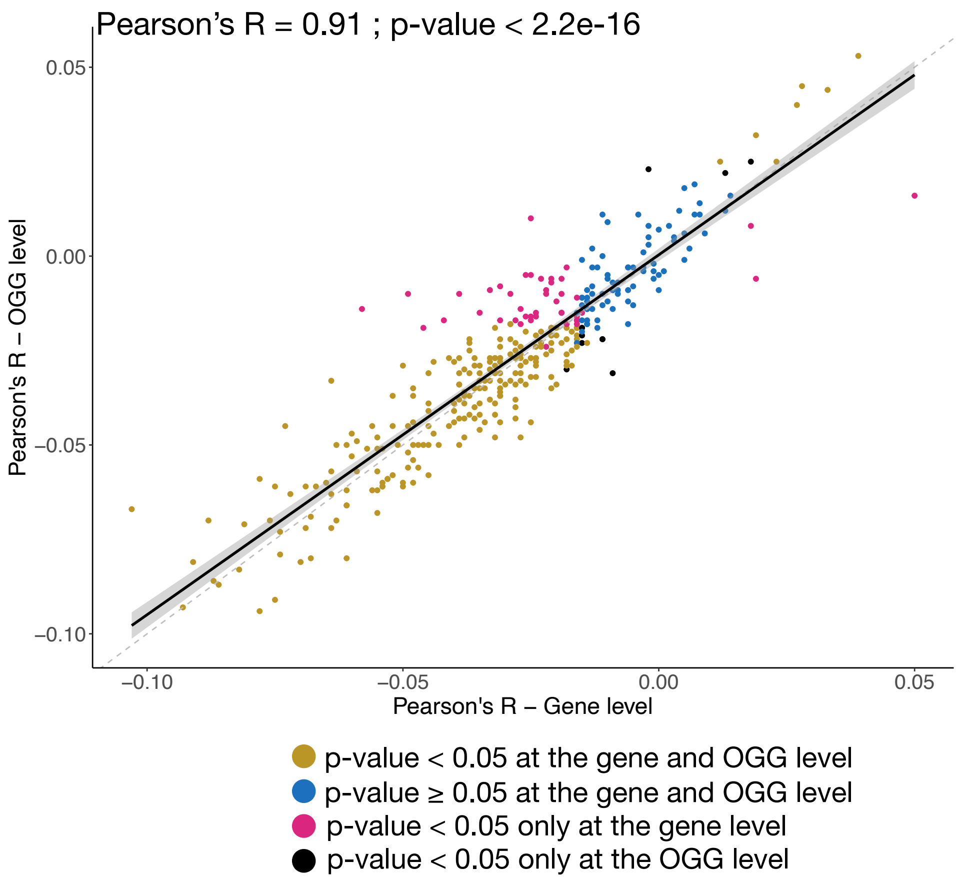


**Supplementary Figure 72.** Correlation between Pearson's Rs computed between TPM and ω at the gene level or at the OG level (taking the mean TPM and mean ω). Pearson correlation coefficient and p-value are indicated. The solid black line represents the linear regression line, and the grey area represents the confidence interval. The dashed line indicates slope = 1 (x=y).

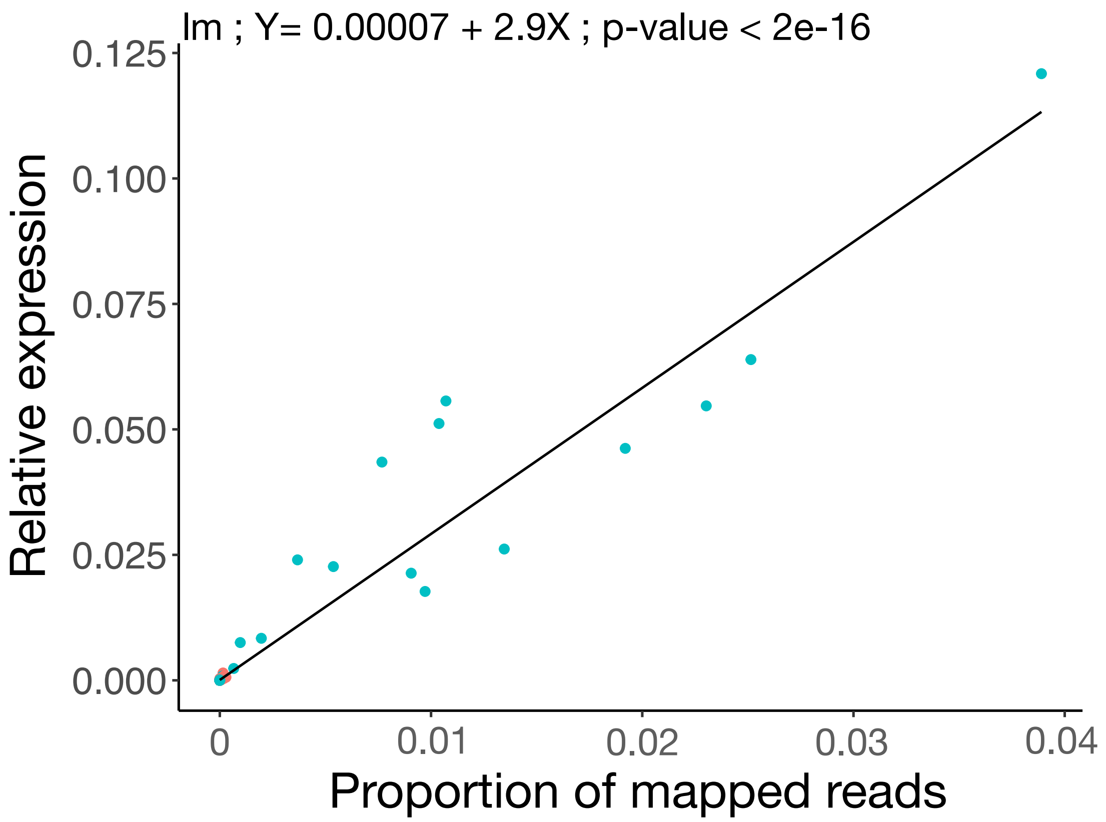


**Supplementary Figure 73.** Relationship between the relative expression of opsins and the proportion of mapped reads that mapped to opsins. The solid black line represents the linear regression line. The linear regression equation and p-value are indicated.
